# Neuronal architecture of the mouse insular cortex underlying its diverse functions

**DOI:** 10.1101/2025.10.23.683697

**Authors:** Bart C. Jongbloets, Yang Chen, Ian K. Gingerich, Michael A. Muniak, Tianyi Mao

## Abstract

The insular cortex integrates interoceptive and exteroceptive information to mediate bodily homeostasis, emotion, learning, and potentially consciousness.^1–4^ However, the cellular and circuit substrates governing the insula and other associative cortices are poorly understood compared to primary cortices. Here, we quantify the dendritic morphology together with electrical properties, local inputs, and/or projections of 1,093 insular pyramidal neurons. These neurons are mapped onto a quantitative anatomical model of the insula based on a Nissl-staining coordinate framework. Using improved algorithms, we define 21 morphological, 12 electrical, and 9 input neuronal types, and identify several morphological and input types that are unique to the insula. Further, we find that morphological properties constrain and often predict inputs, electrical properties, or projection targets. Several morphological types are differentially distributed between the functionally distinct anterior and posterior insula, providing the substrates for a quantitative demarcation between the anterior and posterior insular subregions. Surprisingly, certain neuronal types receive intra-insular inputs originating far beyond canonical cortical columns. Functionally, these connections bridge a long-range thalamus-to-amygdalar circuit that potentially links sensory information to valence. Our work establishes a structure-and-function foundation for investigating the insular cortex.

## Main

The insular cortex (IC), an evolutionarily conserved associative cortex, is an integration hub^5–7^ that orchestrates diverse functions, such as external and internal sensory perception, bodily homeostasis, learning, motivation, and self and emotional awareness.^1–4^ Dysfunction of the IC has profound clinical implications in mood disorders, schizophrenia, and addictive behavior.^1,8,9^ These diverse functions of the IC are in part attributed to topographic subregions along the anterior-posterior axis:^1,8,10^ the anterior IC is associated with cognitive functions including decision-making,^11–13^ emotional valence processing,^1^ and motivational control,^14^ whereas the posterior IC is associated with external and internal sensory processing.^4,15–17^ Interestingly, although distinct, functions of these two subregions of the IC are often related.^1,18^ For example, the anterior IC and posterior IC have been suggested to govern the subjective and objective dimensions of nociception, respectively.^18^ Further, temporally-synchronized electrical activity triggered by nociception arises within multiple locations of the IC along this posterior-to-anterior axis,^19,20^ implying the additional possibility that these related functions are integrated via intra-insular connectivity.^1,18,21^

Beyond the extensively studied anterior-posterior subdivision of IC function, additional heterogeneity also contributes to the various roles of the IC. Structurally, the IC has a non-canonical layer composition that contains agranular, dysgranular, and granular subregions (AI, DI, and GI, respectively), based on the absence, emergence, and full-presence of cortical layer 4 (L4).^3,22,23^ Functional diversity in the IC may also be associated with its differential long-range projections.^7,8^ Indeed, IC projections to the nucleus tractus solitarii (NTS) are associated with motivational vigor,^14^ whereas projections to the mediodorsal (MD) and parafasicular thalamus (PF) are thought to be involved in cognition, associative functions, and pain processing.^6,24^ In parallel, IC projections to the basolateral (BLA) and central amygdala (CeA) are linked to taste aversion and anxiety,^25,26^ and aversive state processing and satiety,^15,27^ respectively. There is also an ongoing debate about whether sweet and bitter taste percepts are processed in distinct anatomical locations of the IC.^28,29^ Nevertheless, despite these and other studies that linked the IC to behaviors^15,25–27,29^ and investigated the gross inputs and outputs of the IC,^5,6,24^ the neuronal substrates that underlie the heterogeneity of function and information processing within the IC remain poorly characterized.

Pyramidal neurons and their local connectivity form the backbone of cortical computation and are crucial for the functional specialization of cortical subregions.^30–33^ Comprehensive cell typing of pyramidal neurons based on dendritic morphology and intrinsic electrical physiology, along with anatomical properties such as layer location and projection targets, has fueled fundamental discoveries of how primary cortices function.^31,33–36^ However, thorough cell typing is not yet available for most associative cortices, including the IC. Furthermore, in addition to communicating with different upstream sources and downstream targets, distinct cell types may also form unique local connections to achieve their specific information processing in individual cortices. Yet, the intricate associations between local connections and other cell-type properties are typically not available comprehensively in primary cortices,^37^ and remain largely unknown for associative cortices such as the IC.

In this study, we systematically examine the cell type identities of pyramidal neurons in the mouse IC based on multiple modalities, including dendritic morphologies, electrical properties, and local inputs, from a dataset of 1,093 quality-controlled pyramidal neurons registered to a topographic atlas of the insula. We find that different cell types can exhibit distinct electrical properties, local inputs, and long-range output targets. In support of functional heterogeneity, local compositions of cell types strongly differ along the anterior-posterior axis of the IC and certain cell types exhibit non-canonical, laterally-extending local connectivity to facilitate intra-insular information propagation. Leveraging our multimodal dataset, we identify biologically relevant correlates of structural and functional hallmarks at the cellular level and provide tools to identify insular cell types in future studies. Further, our results indicate that the aforementioned intra-insular pathway bridges long-range thalamic inputs to the posterior IC with anterior IC outputs to the amygdala, potentially coupling interoception and gustation with taste learning and internal sensing of valence. Together, our study provides insights into the neuronal substrates of the functional diversity in the insula and analytical tools for cell type identification.

## Results

### Anatomical hallmarks of the insular cortex suggest a diverse circuit architecture

The IC contains subregions of unique cytoarchitecture marked by the gradual emergence of a granular L4— AI, DI, and GI—which may support its ability to integrate diverse input-output networks.^5^ To contextualize their underlying cellular types and local circuit connections, we generated an anatomically quantifiable framework of the IC. To register our large-scale dataset onto a precisely defined common reference that accurately captured the subregional and layer demarcations in the IC (Fig. 1a–d and Extended Data Figs. 1– 4), we leveraged distinctions in parvalbumin^+^ (PV^+^) neuron density between the AI and DI/GI (Extended Data Fig. 4a,b),^23^ as well as Nissl staining of cell nuclei, which is considered the ground truth for elucidating cortical layer organization (Fig. 1b). Nissl-stained sections were prepared in a modified horizontal orientation to preserve the functional anterior-posterior axis, annotated (Fig. 1a,c and Supplementary Fig. 1; Methods), and transformed into insular flatmaps using anatomical landmarks and a Laplacian operator (Fig. 1d and Extended Data Fig. 1; Methods). These flatmaps provided a quantitative common reference framework that enabled comparisons and averaging across sections and brains. Derived from these flatmaps, the insular long axis—the widest breadth of its bounds—also provided a close approximation for the functional gradient along the anterior-posterior axis (Extended Data Figs. 1–3; Methods). Strikingly, cortical thickness (pia to corpus callosum) of the AI nearly doubled from posterior (888 ± 35 µm; mean ± standard deviation) to anterior (1641 ± 235 µm) along a 5-mm stretch of the long axis (Extended Data Fig. 1e), which covaried with a near inversion of the relative thickness of superficial (L1–L3; posterior: 32 ± 4%; anterior: 54 ± 3%) vs. deep (L5– L6; posterior: 62 ± 4%; anterior: 34 ± 1%) layers (Extended Data Figs. 2 and 3; local claustral thickness included in calculation). Cortical thickness of the DI/GI increased only moderately toward the anterior side (17 ± 23% across a 3-mm stretch), primarily driven by a relative expansion of L5. Furthermore, the relative proportions of different cortical layers varied between AI and DI/GI (Extended Data Fig. 3c). As an illustration, we established subregion-specific layer demarcations (Methods) and normalized to the cortical thickness (AI: L1 = 13 ± 5%, L2 = 25 ± 7%, L3 = 43 ± 9%, L5 = 68 ± 2%, L6 = 87 ± 4%; DI/GI: L1 = 10 ± 3%, L2 = 22 ± 3%, L3 = 41 ± 6%, L4 = 54 ± 6%, L5 = 74 ± 3%, L6 = 92 ± 3%; mean ± standard deviation). Overall, the marked differences across subregions and along the long axis suggested the potential presence of specialized circuit architecture across subregions and functional axes, emphasizing the necessity to align our cell atlas within this three-dimensional topography.

**Fig. 1:**
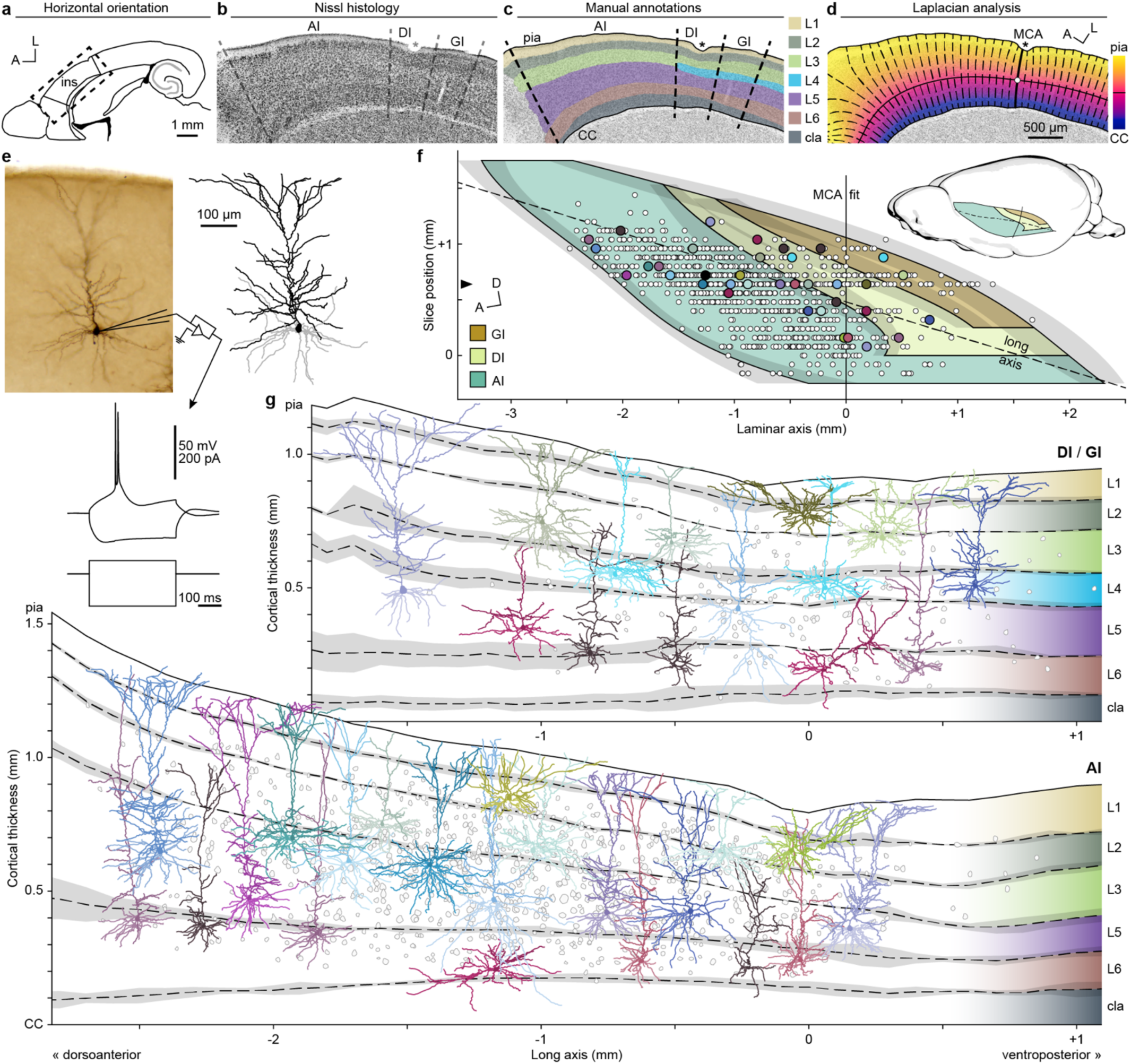
Anatomical hallmarks of the insular cortex suggest a diverse circuit architecture. **a**–**c**, Example horizontal section centered on the insular cortex with Nissl staining (**b**) and manual annotations (**c**; Supplementary Fig. 1). Boundary annotations (**a**) adapted from Franklin and Paxinos.^45^ Dashed lines: subregion boundaries; asterisk: middle cerebral artery (MCA) position; color blocks in (**c**): manual cortical layer delineations between the pia and corpus callosum (CC) boundary lines (solid). **d**, Solution to Laplace’s equation on the same section from (**b** and **c**). Thin black line: trajectory of laminar cortical axis; white circle: MCA-derived coordinate origin for section (Extended Data Fig. 1; Methods); solid streamline: MCA reference; dashed streamlines: Laplacian gradients at 100-µm intervals relative to MCA reference. **e**, Photomicrograph (top-left) of an example DAB-stained neuron from an acute slice following physiology characterization (bottom) and biocytin fill, with post hoc computer-assisted morphological reconstruction (top-right) showing apical (black) and basal (gray) dendritic branches. Traces (bottom) show physiology recordings in response to current injections. **f**, Topographic bounds of insular subregions (filled contours) and standard deviation of their laminar boundaries (gray shading) are represented on a 2D flatmap (Extended Data Fig. 1). Solid line: the MCA fit (Extended Data Fig. 1b); dashed line: long axis (Extended Data Fig. 1d). Black triangle on y-axis: position of example Nissl section from (**b**–**d**). Inset: approximate position of flatmap on whole mouse brain (adapted from Janke et al.^103^). **g**, Topographic distributions of recorded insular neuronal somas (gray outlines) and selected examples of reconstructed dendrites in the DI/GI (top) and AI (bottom) in their assigned cortical layers (Extended Data Figs. 2–4). Sampled cells from (**e**) and (**g**) are highlighted with circles of corresponding colors in (**f**). Layer boundaries calculated based on flatmaps from three animals are shown as mean (dashed lines) and standard deviation (gray shading) (Extended Data Figs. 1e and 3c). A, anterior; D, dorsal; L, lateral; AI, agranular insula; DI, dysgranular insula; GI, granular insula; cla, claustrum; ins, insula; CC, corpus callosum; L1-L6, cortical layers 1-6; MCA, middle cerebral artery.

### Generation of a position-mapped multimodal neuronal dataset

To investigate insular cell types, we recorded the electrical activities of neurons in acute insular brain slices in a modified horizontal orientation (Methods) using whole-cell patch clamp. PV^+^ neuronal density was used as a guide in acute slices to help target neurons in AI or DI/GI (Extended Data Fig. 4a,b). Recording electrodes containing biocytin allowed for post hoc staining to reveal dendritic morphology with high contrast (Fig. 1e; Methods). Using a Neurolucida microscope system, we reconstructed dendritic morphologies in their entirety (Fig. 1e and Extended Data Fig. 4c–f; Methods). We registered these reconstructions to adjacent PV- and Nissl-stained sections in our common reference framework for three-dimensional spatial matching. Soma locations were subsequently placed onto the average insular flatmap (Fig.1f and Extended Data Fig. 4g–i; Methods), highlighting comprehensive coverage across insular subregions (Fig. 1f) and cortical layers (Fig. 1g). Mapping our cell atlas to this common reference framework set the stage for investigations on the relationship of IC cell types with the anatomical and functional topography of the insula.

### Insular pyramidal neurons display distinct morphological types

To comprehensively identify insular pyramidal cell types, we classified neurons based on interpretable dendritic morphology features. We analyzed 958 high quality dendritic reconstructions (Supplementary Fig. 2) and extracted a curated list of morphological features built upon known properties in other cortical regions.^35,38,39^ These features included the shape and length of local dendritic arbors relative to the soma or pial surface, as well as additional components to capture spatial dendritic complexity (Supplementary Table 1). We then performed an iterative consensus cluster analysis (iCCA), with the initial round of clustering based on principal component analysis (PCA) transformed data (Methods). This approach allowed us to select non-redundant interpretable features, which resulted in high co-cluster scores to yield effective and robust clustering (Fig. 2a and Extended Data Fig. 5a–c; Methods). To assess whether we had achieved an adequate sample size of neurons, we repeated the iCCA on incremental dataset sizes of subsampled original data or on simulated data (Fig. 2b; Methods). Using both approaches, the number of identified clusters did not monotonically increase beyond 400 cells and was stable beyond 800 cells, suggesting that our data size was sufficient to capture putative insular cell types.

**Fig. 2:**
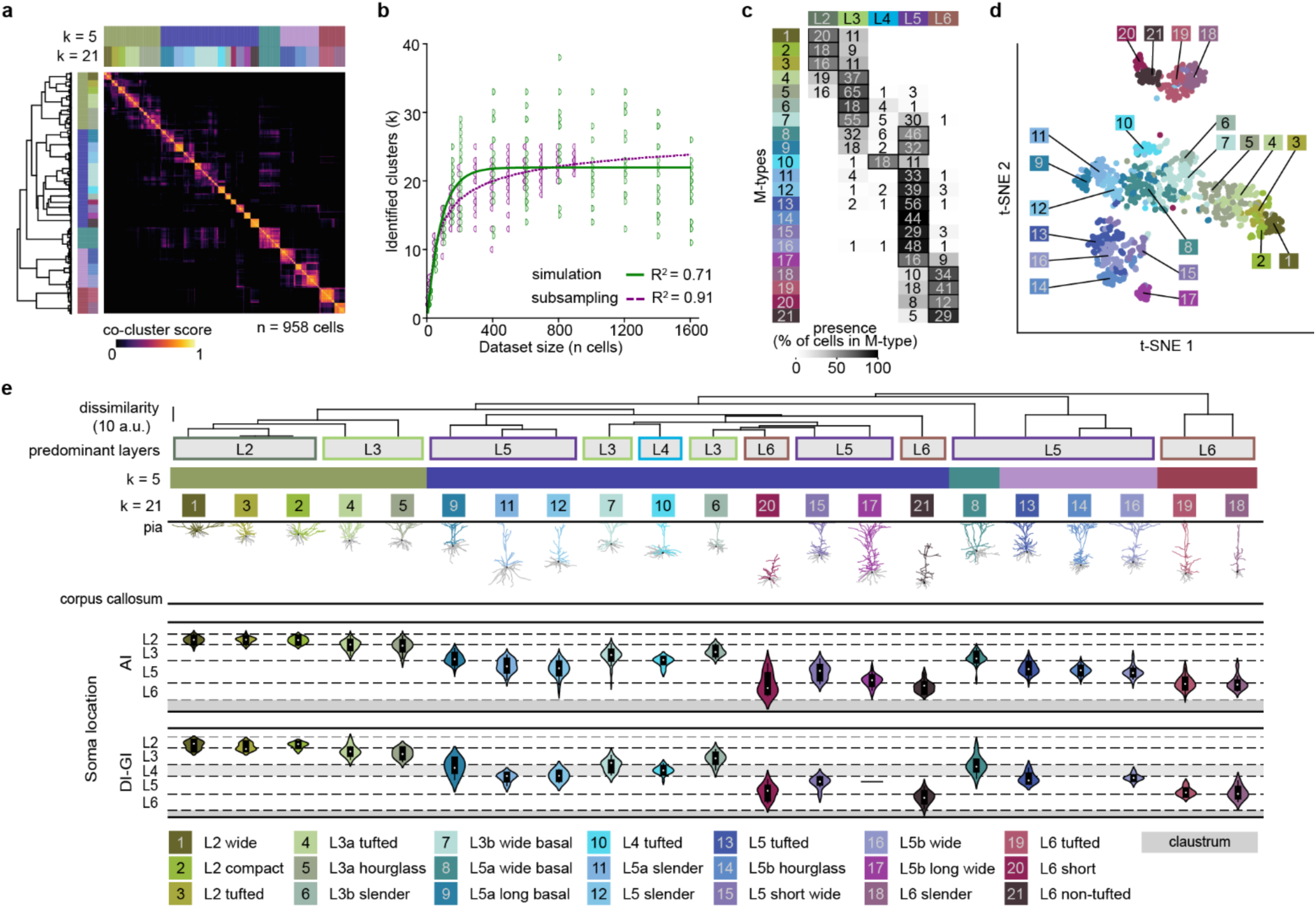
Insular pyramidal neurons display distinct morphological types. **a**, Hierarchical cluster analysis (HCA) on pair-wise co-cluster scores across 958 cells using the best performing morphological feature set. Dendrogram-based grouping is color-coded for 21 clusters (k = 21) based on optimal silhouette score, as well as for 5 clusters (k = 5). **b**, Number of identified clusters as a function of dataset size based on iterations of HCA on subsampled data (purple) or simulated data (green). Solid and dashed lines: logistic regression fit on subsampled and simulated data, respectively. **c**, M-type cell counts by layer. Horizontal colored boxes: layer identities. Vertical colored boxes: M-type aliases sorted by median soma location, color-coded corresponding to k = 21 results in (**a**). Number in gray boxes: cell counts per layer per M-type. Grayscale luminance indicates the percentage of counts per layer for each M-type. The predominant layer allocations are boxed (presence > 45%). **d**, t-SNE distribution of M-types. Colored boxes with numbers: M-types color-coded for k = 21 in (**a**). **e**, M-type predominant layer(s) and hierarchical dendrogram allocations (top) with their representative morphologies (middle) and soma distributions in the agranular (AI) and dysgranular/granular (DI/GI) insular cortex (bottom). M-type aliases are based on predominant layer(s) allocation and key morphology features.

Our morphology-feature based iCCA yielded 21 morphology types (M-types) (Extended Data Table 1), comparable to the number of M-types in other cortical areas.^35,38,39^ Reduced cluster numbers (e.g., k = 3, 5, or 14) resulted in groupings of cells that were highly heterogenous in layer location and morphology (Fig. 2a,e and Extended Data Fig. 6a). We named these M-types based on their predominant layer allocation and morphological characteristics (Fig. 2c–e and Extended Data Figs. 5d,e and 6). These feature-based cluster allocations were in general agreement (adjusted Rand index = 0.36) with the allocations (k = 20) obtained from the initial PCA-based analysis (Extended Data Fig. 5f; Methods). This confirmed that a reduced list of 29 out of the initial 67 interpretable features identified by our iCCA approach covered the most variability in the dataset (Extended Data Table 1). To validate that this feature shortlist was sufficient to classify M-types and that our cluster allocations were stable and generalizable, we trained random forest classifiers using subsets of the original data (75%) with the selected morphology features and achieved ∼84.8% accuracy (median F1-score, range 82.9 – 85.3%, n = 5 iterations, Extended Data Table 1) on the withheld dataset (Extended Data Fig. 5g). These classifiers are, therefore, reliable tools that require a minimal list of extracted features to allocate M-type labels to dendritic morphologies in the IC.

Analyses of M-types and their associated morphological features revealed several remarkable characteristics (Extended Data Fig. 6b). Certain M-types, for example, had total dendritic lengths extending over ten millimeters (*L5b long wide* [M-type 17] and *L5b hourglass* [M-type 14]; Extended Data Fig. 6b), which is rarely observed in other mouse cortical regions.^35,40,41^ Several other features, including long basal dendritic lengths, large maximal apical branch orders, and large numbers of apical nodes near somas, highlighted the vast extent and dendritic complexity of several neuronal types, e.g., *L5a long basal* (M-type 9), *L5b long wide* (M-type 17), and *L5 short wide* (M-type 15), respectively (Extended Data Fig. 6b). The elaborated postsynaptic compartments in these neuronal types may enable potential integration from complex inputs. Intriguingly, despite a saturating sampling depth for quantifying cell types (Fig. 2b), we did not find neurons resembling von Economo neurons (VENs) (Supplementary Fig. 2),^42^ which have been observed in higher order species and are implicated in conciousness.^1,8,43^ In addition, M-types mapped on a dimensionality-reduced representation using t-distributed stochastic neighbor embedding (t-SNE) showed separation of cell types within different layers (Fig. 2d). The separation of the larger L5 morphologies (M-types 14, 16, and 17)—primarily in the deeper L5 (Fig. 2e and Extended Data Fig. 5d)—from the smaller L5 morphologies (M-types 8 and 9) highlighted the presence of canonical L5a and L5b pyramidal neurons (Fig. 2d).^40,44^ Consistent with empirical notions of properties that are highly relevant in defining cortical pyramidal neuron types, soma location, total apical length, and the ratio of apical-over-basal dendritic height were among the most informative features in the classifiers (Extended Data Fig. 5e).

### Insular M-types are enriched in subregions and have preferential projection targets

Although a major theme of understanding functional diversity in the IC is spatial separation along its anterior-posterior axis,^1,8,10^ the precise demarcation is not well defined.^14,15,25,26^ As mentioned earlier, we observed a posterior-to-anterior broadening of cortical and layer thickness along the long axis, especially within the AI (Fig. 1g and Extended Data Figs. 1–3). Accordingly, we tested whether M-types of the IC were differentially distributed across its long axis. This was indeed the case (Fig. 3a–d). Most notably, several M-types within the same layer, for example *L5b hourglass* (M-type 14) and *L5b wide* (M-type 16), exhibited strong topographic preferences either towards dorsoanterior or ventroposterior sides (Fig. 3a,b and Extended Data Fig. 7a). To systematically quantify this tendency for each individual M-type, we first sought to objectively define an anatomical division of the AI (‘topographic split’). We mapped the soma distributions of all M-types along the insular long axis and performed pair-wise comparisons of each M-type with other M-types that were in the same layer (Fig. 3b and Extended Data Fig. 7a). Most pairs within L5 and L6 showed differential soma distributions along this axis (Fig. 3b and Extended Data Fig. 7a). A segregation point was calculated for each distinctly distributed pair (exemplified in Fig. 3a; Methods). The average of individual segregation lines resulted in a ‘topographic split’ line centered at −1,052 ± 48 µm (standard error; n = 18 segregation lines), yielding an objective division along the insular long axis (Fig. 3b,c), potentially related to the functionally distinct anterior and posterior insula.^14,15,25,26^ Unlike the DI/GI, the AI stretched across most of the insula’s anterior-posterior axis; we therefore used this topographic split to define dorsoanterior- and ventroposterior-AI. To facilitate future studies of these subdivisions, we registered the topographic split from our flatmap back onto our common reference frame (Extended Data Fig. 1a) and also registered the Franklin and Paxinos horizontal mouse atlas^45^ to this same reference frame (Fig. 3c; Methods). This allowed us to estimate the stereotaxic coordinates for targeting the dorsoanterior-AI (−2.56, 2.20, 2.55 mm; dorsal-ventral, anterior-posterior, and lateral-medial relative to bregma) and ventroposterior-AI (−3.76, 1.00, 3.80 mm) (Methods). Next, based on this topographic split, we re-evaluated which individual M-types were enriched in the dorsoanterior- or ventroposterior-AI according to this group-averaged division line (Fig. 3b; Methods). Eight M-types located in L2, L5, and L6 had statistically significant enrichment in either dorsoanterior- or ventroposterior-AI (Fig. 3d). The existence of distinct M-types in specific AI subregions implies specialized local insular subcircuits.

**Fig. 3:**
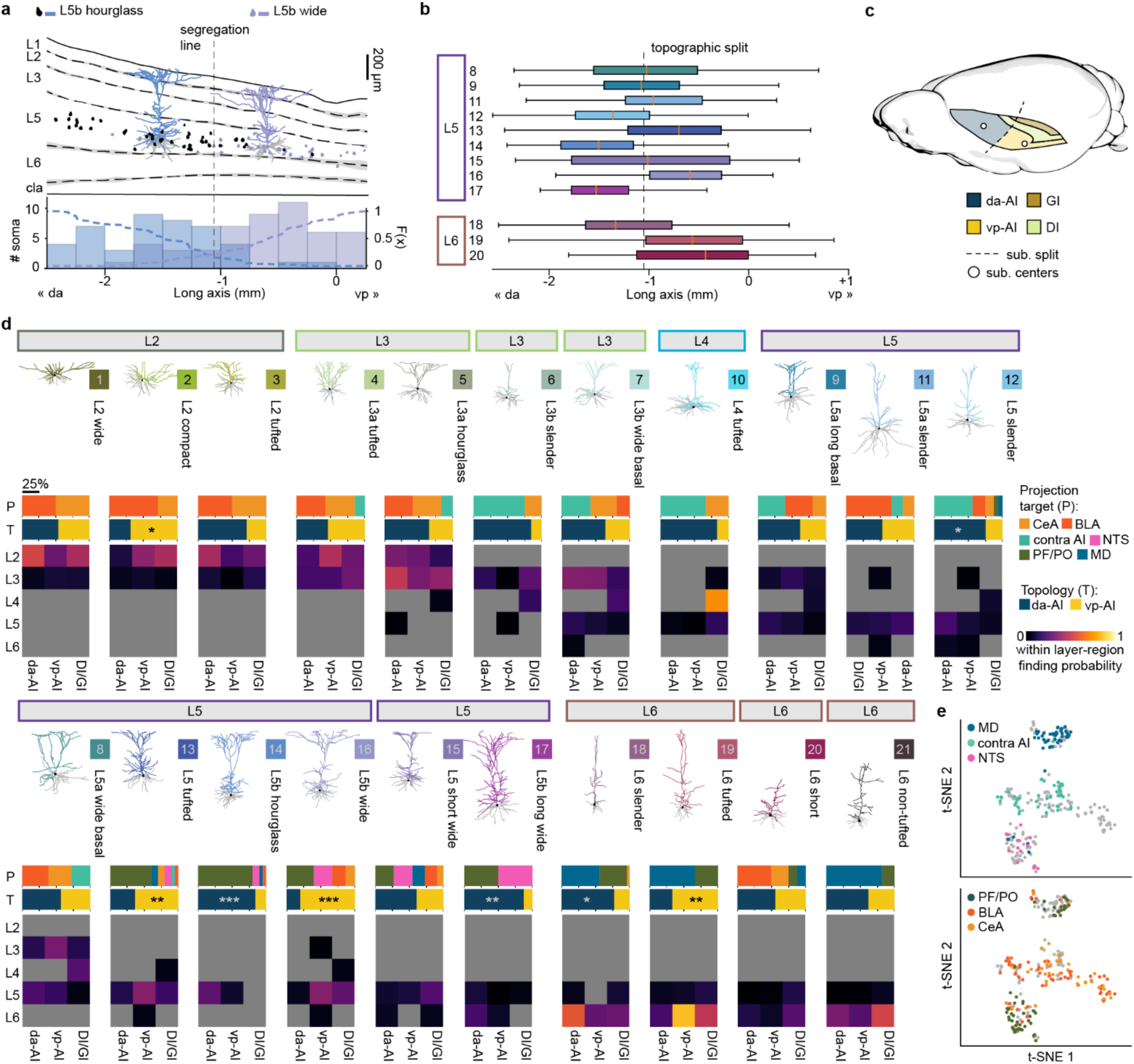
Insular M-types are enriched in subregions and have preferential projection targets. **a**, Soma distribution (top; Fig. 1g) and corresponding histograms (bottom) of two M-types (*L5b hourglass* and *L5b wide*). Colored bars and dashed lines (bottom): soma count (left y-axis) and cumulative distribution probability (right y-axis), respectively, along the long axis. Black dashed vertical line: segregation line at which the sum of cumulative distribution probability of the two M-types is the smallest. **b**, Boxplots of L5 and L6 M-types with significant topographic shift compared to another within-layer counterpart (Mann-Whitney U test with post hoc false-discovery rate Benjamini-Hochberg correction for multiple testing). Outliers not shown (Extended Data Fig. 7a for the full dataset). Topographic split (dashed line): average position of segregation lines from all significant within-layer comparisons. Orange ticks: median location for each M-type. **c**, Three-dimensional representation of the topographic split location (dashed line) within a model mouse brain. White circles: representation of suggested coordinates (see main text) to target dorsoanterior-AI and ventroposterior-AI subregions. **d**, Quantification of projection targets (P, top bar plot), topography (T, bottom bar plot), and within-layer and -region finding probabilities (bottom heatmap) per M-type. M-types are sorted based on predominant layer allocation and grouped by dendrogram hierarchy as shown in Fig. 2e. Representative dendritic morphologies were normalized to their local cortical thickness. Statistics on topographic distributions were performed using Fisher exact test with post hoc false-discovery rate Benjamini-Hochberg correction for multiple testing (Methods). Within-layer and -region finding probabilities: the number of neurons for a given M-type allocated to a specific layer and subregion (e.g., L2 of da-AI) divided by the total number of neurons in that layer and subregion. *: p ≤ 0.05, **: p ≤ 0.01, ***: p ≤ 0.001. **e**, Projection target identities revealed in t-SNE representations of the morphology dataset, as shown in Fig. 2d. da: dorsoanterior; vp: ventroposterior; BLA: basolateral amygdala; CeA: central amygdala; contra AI: contralateral agranular insula; MD: mediodorsal thalamus; NTS: nucleus tractus solitarius; PF/PO: parafascicular/posterior thalamus; cla: claustrum.

We next asked whether AI and DI/GI encompass different compositions of M-types. Indeed, *L5b hourglass* (M-type 14) and *L5b long wide* (M-type 17) cells were almost exclusively found in the AI, but rarely seen, if not completely absent, in the DI/GI (Figs. 2e and 3d). *L4 tufted* neurons (M-type 10) residing in the DI/GI resembled the typical L4 pyramidal neuronal morphology in primary visual and auditory cortices.^46,47^ Yet, spiny stellate cells, such as those in the somatosensory cortex, were not identified in the IC.^46,48^ Notably, a sparse presence of *L4 tufted* neurons (M-type 10) was also observed in the dorsoanterior- and ventroposterior-AI. In addition, *L3b slender* (M-type 6) and *L3b wide basal* (M-type 7) neurons located adjacent to L4 had morphological characteristics most similar to *L4 tufted* neurons (M-type 10) (Fig. 2d,e and Extended Data Fig. 5e) and were predominantly in the dorsoanterior-AI (Fig. 3d). Our topographic analysis, therefore, identified L4-like neurons and subregion-specific M-types in the AI.

Cortical pyramidal neurons relay information to specific brain regions via long-range axonal projections and their projection targets are often associated with morphological characteristics.^49,50^ To identify anatomical and morphological features of projection neurons in the IC, we traced and quantified the soma distribution of projection neurons across cortical depth using retrograde tracers injected into the CeA, BLA, MD, PF/posterior thalamus (PO), NTS, and contralateral AI (contra AI) in different animals. The soma distributions of contra AI, NTS, PF/PO and MD projection neurons showed clear layer preferences, whereas corticoamygdalar-projecting neurons were distributed across L2, L3, L5, and with a moderate presence in L6 (Extended Data Fig. 7b,c). To identify morphological characteristics of these different projection neurons, we analyzed the subset of our reconstructed morphologies that included projection target information based on retrograde tracer injections (n = 271 neurons). Mapping these onto the t-SNE morphology feature space revealed that neurons with the same projection target, such as NTS- or MD-projecting neurons, shared overlapping morphological traits (Fig. 3e). Projection targets also constrained M-types (Extended Data Fig. 7d). For example, MD-projecting neurons largely resembled *L6 tufted* (M-type 19) and *L6 slender* (M-type 18) M-types but rarely *L6 short* (M-type 20), which projected mainly to the amygdala. Notably, the majority of NTS-projecting neurons were *L5b wide* (M-type 16), primarily located in the ventroposterior-AI, and *L5b long* wide (M-type 17), primarily found in the dorsoanterior-AI.

Together, we comprehensively identified insular pyramidal M-types that were associated with specific layers, topographic distributions, and projection targets (Fig. 3d), which may underlie IC subcircuit diversity.

### Distinct electrical profiles associate with insular layers and M-types

To investigate the range of distinct neural processing of electrical signals in the insula, we analyzed the intrinsic electrical properties of 306 neurons in whole-cell current-clamp (Fig. 4 and Extended Data Fig. 8). Similar to our M-type analysis pipeline, we conducted iCCA starting with 29 electrical features, which captured intrinsic passive, rectifying, and spiking properties (Methods). We identified 12 electrical profiles (E-profiles) based on 16 informative features (Fig. 4a–d, Extended Data Fig. 8a–d, and Extended Data Table 1). Prominent features of specific E-profiles (Fig. 4a) included input resistance (Rin: low vs. high in E-profiles 1 vs. 11, respectively; Extended Data Fig. 8c,d) and the presence of rectifying properties (afterhyperpolarization: E-profiles 5 and 12, and hyperpolarization-induced voltage sag: E-profiles 8 and 9; Fig. 4d and Extended Data Fig. 8c,d). E-profile grouping based solely on spiking patterns resulted in three main electrical classes (E-classes): regular spiking, burst spiking, and spike adapting (Fig. 4a,b), as described in other cortices.^35,38,40^ Interestingly, in contrast to electrical property-based cell types identified in primary visual cortex,^35^ our burst spiking E-class (E-profiles 6 and 8) was strictly separated from other E-profiles by their fraction of short inter-spike interval and first spike interval (Extended Data Fig. 8b,d). Within each class, the E-profiles further defined physiological traits regarding passive and rectifying properties, for example by differentiating bursting cells based on the presence of a voltage sag (Fig. 4a, E-profiles 6 vs. 8). The E-classes occupied well-separated spaces in the dimensionality-reduced t-SNE plot, suggesting they are bona fide distinct categories (Fig. 4e and Extended Data Fig. 8c). The 12 individual E-profiles did not show clear layer predominance or topographic bias (Extended Data Fig. 8e), contrary to what was observed with M-types. However, the relative composition of E-profiles and -classes differed in each layer (Fig. 4f and Extended Data Fig. 8e). Notably, L2 neurons were mainly regular spiking, whereas L3 neurons displayed spike adaptation or regular spiking, but infrequently burst spiking behavior, and L5 neurons displayed all three main electrical classes. While L2 and L3 are often lumped together as one layer, our result suggested an interesting segregation of L2 and L3 in the IC based on electrical properties, which is consistent with our anatomical annotation (Fig. 2c,e). The differential layer presence of E-classes might contribute to different capabilities for relaying information in different layers of the IC.

**Fig. 4:**
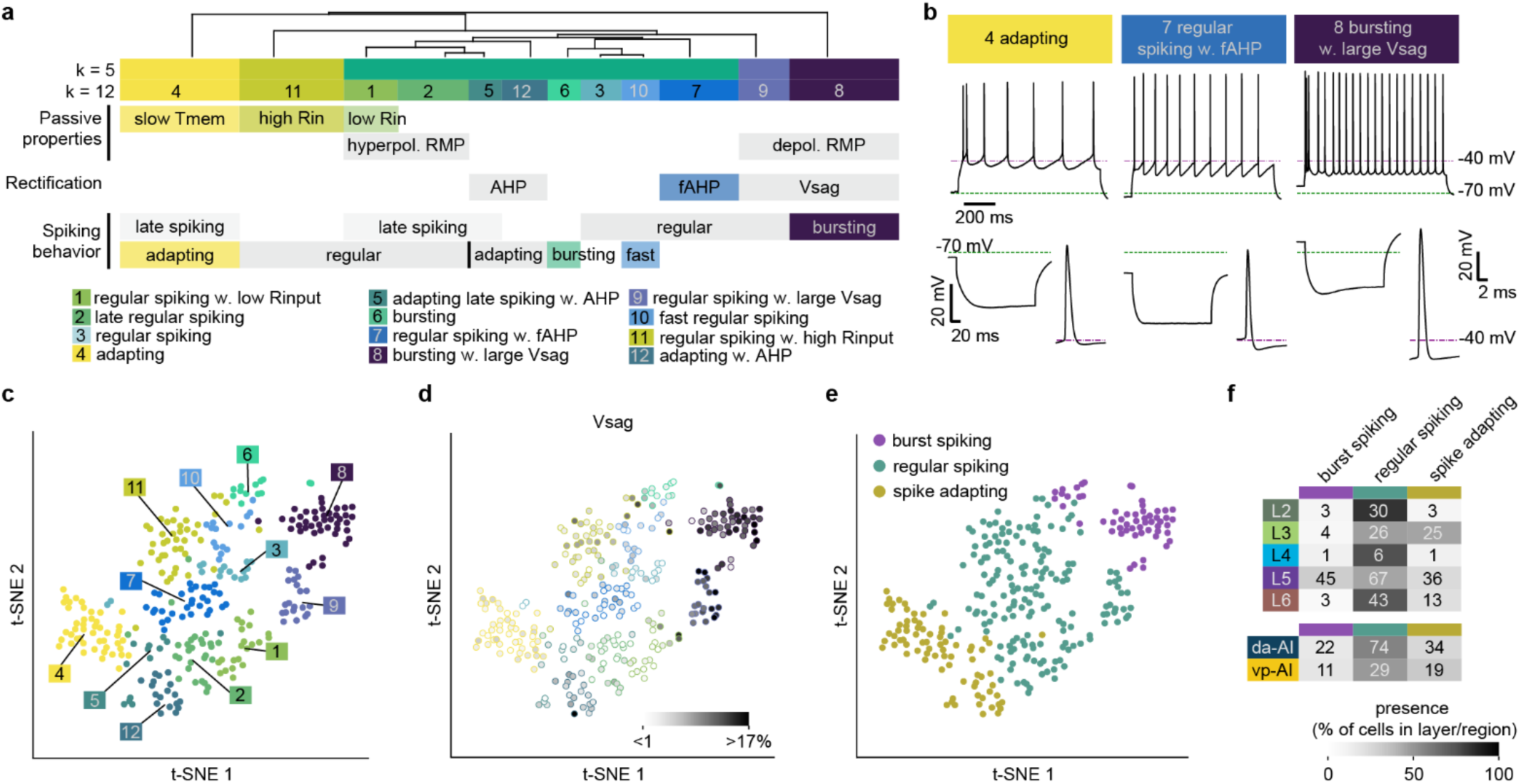
Distinct electrical profiles associate with insular layers and M-types. **a**, E-profiles and their most prominent electrical features. **b**, Representative voltage traces of three contrastive E-profiles, including action potential spiking patterns during suprathreshold stimulation (rheobase + 125 pA, top), responses to hyperpolarizing current injection (−300 pA, bottom left for each E-profile), and isolated action potentials near rheobase (bottom right for each E-profile). **c**, t-SNE distribution of E-profiles. Colored boxes with numbers: E-profiles color-coded for k = 12 in (**a**). **d**, Distribution of voltage sag, a representative electrical feature contrasting E-profiles. The parameter amplitudes are shown in grayscale luminance of the filled dots, mapped onto the t-SNE plot in (**c**). Edges of the filled dots are color-coded by E-profiles. **e**, Assignment of the three electrical classes (E-classes)—burst spiking, regular spiking, and spike adapting—mapped onto the t-SNE plot in (**c**). **f**, Contingency tables of layer (top) or AI subregion (bottom) distributions for the three E-classes. Grayscale luminance: the percentage of cells in an E-class allocated to a given layer or subregion. Numbers in gray boxes: n of observed cells.

### Insular pyramidal neurons receive non-canonical laterally-extending local excitatory inputs

The computational diversity of individual pyramidal neurons is shaped not only by anatomical, morphological, and electrical properties but also by local connectivity across layers. Towards this end, we systematically investigated the excitatory local inputs to IC neurons using laser scanning photostimulation (LSPS) with glutamate uncaging, presented as heatmaps indicating input origins and their strength (Extended Data Fig. 9; Methods).^51^ We grouped the input maps by the layer location of the postsynaptic soma. Representative (Fig. 5a) and averaged maps (Fig. 5b and Extended Data Fig. 9d–f, j) showed heterogeneous input origins and strengths onto the pyramidal neurons located in different layers. Strikingly, non-canonical local inputs extended laterally along the anterior-posterior axis, sometimes originating from the DI/GI (Fig. 5a). To summarize the flow of information across insular layers, we generated a local input diagram based on the absolute inputs originating from each layer, both within-column and laterally-extending (≤ or > 750 µm distance from the soma, respectively; Fig. 5c; Methods). This uncovered several general characteristics of local circuits in the IC. First, L2 and L3, although deemed superficial cortical layers and often grouped together,^34,52^ received differential local inputs (Fig. 5b,c and Extended Data Fig. 9d). Specifically, L3 received stronger inputs than L2, especially for inputs from L2, L3, and L6 (Fig. 5b and Extended Data Fig. 9d), although the normalized input patterns were similar between these two types of neurons (Extended Data Fig. 9e). These results indicated distinct roles of L2 and L3 in local AI processing, consistent with the suggestion of diverging L2 and L3 microcircuits in some primary cortices.^53–55^ Compared to superficial layers, the deep layers of the AI, L5 and L6, received marked inputs from the claustrum and at locations extending farther away from the soma, which were both distinctive compared to canonical microcircuits in primary cortices.^54,56^ In particular, L5 neurons received laterally extending inputs from all layers, but the strongest and most distal were from superficial layers, often posteriorly biased and up to 1.5 – 2 mm away from the soma (Fig. 5a–c and Extended Data Fig. 9d,e). In contrast to L5, L6 received the strongest laterally-extending inputs from deep layers in both anterior and posterior directions and received superficial inputs that originated mostly within-column (Fig. 5b and Extended Data Fig. 9d,e). Such interlaminar and lateralized local inputs to L6 are rare in canonical microcircuits in other cortices.^37,54,56^

**Fig. 5:**
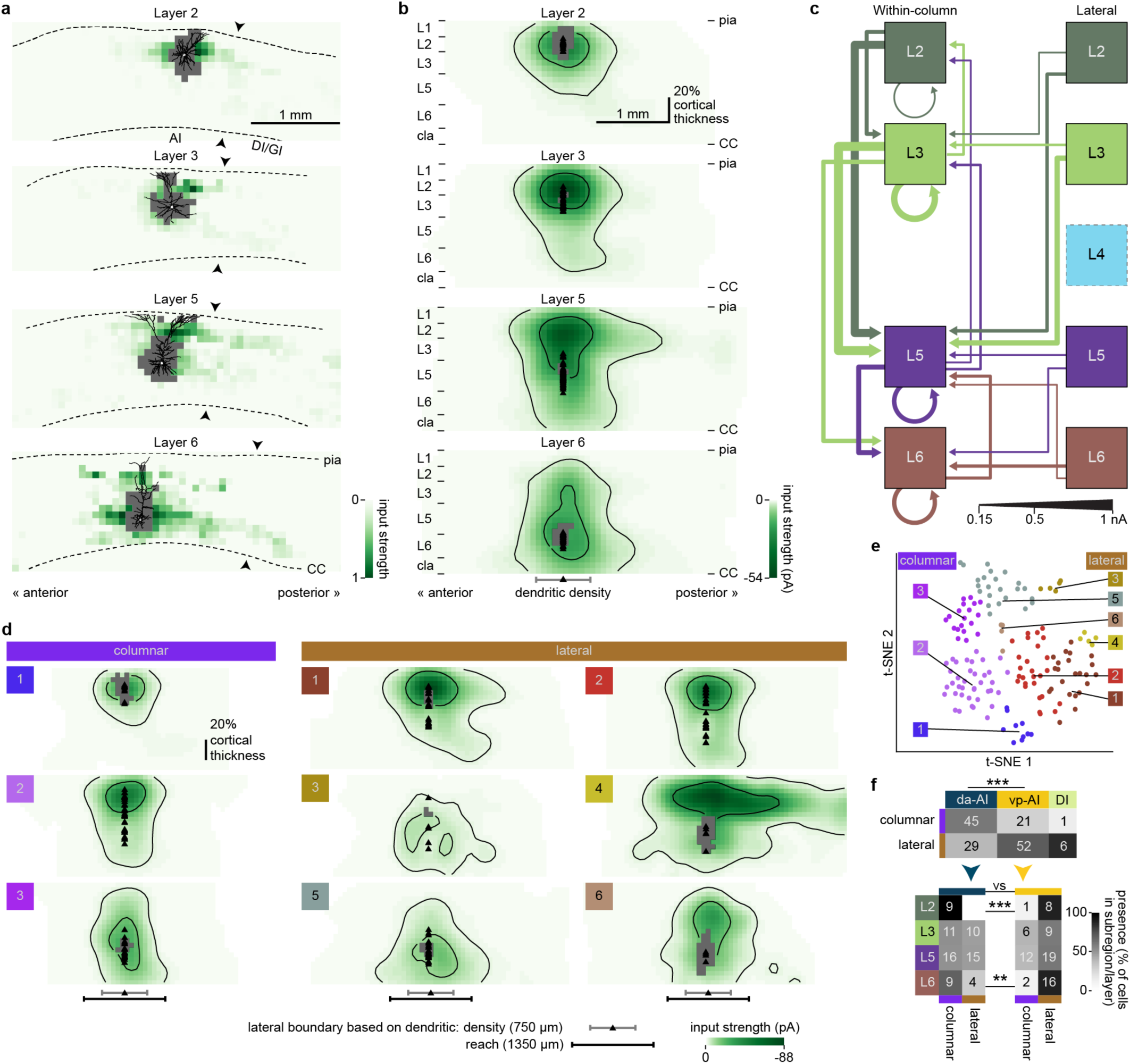
Insular pyramidal neurons receive non-canonical laterally-extending local excitatory inputs. **a** and **b**, Example (**a**) and average (**b**) local input maps of L2 (n = 18), L3 (n = 36), L5 (n = 64), and L6 (n = 35) neurons aligned by pia. The example maps are shown in a linear color scale; the average maps are Gaussian-filtered for visualization, shown in an exponential color scale to highlight distal inputs, and contoured at 15^th^ and 85^th^ percentiles of all responses (Methods). Arrowheads: estimated boundary between AI and DI/GI (left and right of arrowheads, respectively); gray pixels: direct response sites; black triangles: soma locations. **c**, Circuit diagram of local excitatory inputs to AI layers. Arrow thickness indicates mean input strength arriving at each layer. Only input strengths above 0.15 nA are displayed. Left: within-column inputs summarizing inputs confined within the dendritic density; right: lateral inputs summarizing distal inputs extending beyond the dendritic density (Methods). The L4 box is semitransparent because L4 is absent in the AI and is beyond the sampling range for some neurons. Note, L2 neurons received, on average, 0.412 nA lateral input from all layers combined (Extended Data Fig. 9j), but inputs from any individual layer did not exceed the display threshold. **d**, Average local input maps of the nine I-profiles, Gaussian-filtered for visualization, shown in an exponential color scale to highlight distal inputs, and contoured at 15^th^ and 85^th^ percentiles of all responses. Gray pixels: direct response sites; black triangles: soma locations. **e**, I-profile allocations revealed in the t-SNE-transformed local input feature space, color-coded as in (**d**). **f**, Contingency tables of the input classes vs. subregions (top) and layers (bottom). Grayscale luminance: the percentage of cells with either input class within a given subregion or layer. Numbers in gray boxes: n of observed cells (when n > 0). CC: corpus callosum.

Comparing microcircuits of all layers, intralaminar connectivity, although underestimated due to the exclusion of synaptic responses inseparable from direct responses, was evident within all layers and strongest in L3, followed by L5 and L6 (Fig. 5c). In addition, we identified reciprocal interlaminar connections, characterized by strong descending inputs (e.g., L2→L5, L3→L5) and relatively weak ascending inputs (e.g., L5→L2, L5→L3) (Fig. 5c). Similar to the agranular motor cortex,^54,57^ the strong ascending inputs to L2/3 observed in granular cortices^54,58,59^ were not observed in the AI, supporting the observation that local circuits of agranular and granular cortices are distinctive. Importantly, we confirmed that both intralaminar and interlaminar inputs can originate beyond the span of a typical cortical column (750 µm distance from the soma; Methods), with the strongest lateralized inputs coming from L2 and L3 to L5 (Fig. 5c).

To systematically analyze input patterns beyond laminar location,^58^ we identified nine prototypical input profiles (I-profiles) using our iCCA approach (Fig. 5d–f and Extended Data Fig. 10; Methods). Six out of nine I-profiles showed pronounced lateralized inputs, with more lateral inputs located beyond the dendritic density border (750 µm, Kruskal-Wallis H (8) = 80.77, p < 0.001) and sustained input probabilities located beyond the maximal dendritic reach (1350 µm; Extended Data Fig. 10d,e; Methods). Moreover, these six I-profiles were well separated in the t-SNE plot from the three I-profiles that displayed primarily local and columnar inputs (Fig. 5e), confirming the bipartite distinction between the columnar and lateral input classes.

Are the input patterns of IC neurons related to their layer or subregion locations? Indeed, similar to the subregional preferences of morphology-defined cell types, neurons located in the dorsoanterior- or ventroposterior-AI received predominantly columnar or lateral inputs, respectively (Fisher’s exact p < 0.001; Fig. 5f and Extended Data Fig. 10f,g). Interestingly, such predominance was only observed for L2 and L6, but not L3 and L5 neurons, likely reflecting the functional divergence of L2 and L6 neurons in dorsoanterior- and ventroposterior-AI, rather than a potential failure to preserve lateral inputs in brain slices. These results indicate that columnar or lateral integration preferentially occurs depending on the anatomical location of neurons.

### Multimodal analyses distinguish and predict specific insular cell types

#### Morphology and projection targets associate with specific electrical and input profiles

Progress in cortical cell typing based on a single modality has proven necessary but insufficient for understanding cortical cell types.^35,36^ To comprehensively associate multiple modalities that were used to define a cell type, we generated a visualization of all individual IC cells across all modalities and properties in our dataset (Fig. 6a; Methods) where cell types and modalities could be queried (MouseInsularAtlas; https://gitlab.com/maolab/mouseinsularatlas). Interestingly, morphological types and projection targets were strong indicators of electrical properties of a pyramidal neuron (Fig. 6b,c, and Extended Data Fig. 11a). Mapping M-type identities onto the t-SNE plot of electrical features indicated that several M-types, located within the same layer, shared electrical properties (Fig. 6b and Extended Data Fig. 11a). For example, a majority of *L5b long wide*, *hourglass*, or *wide* neurons (M-types 14, 16, or 17, respectively), displayed strong hyperpolarization-induced voltage sags and depolarized resting membrane potentials (E-profiles 8 and 9) (Fig. 4 c,d, and Extended Data Fig. 8d), while displaying two modes of spiking behaviors, bursting and regular spiking (Figs. 4e and 6b). These morphologies and related physiology were reminiscent of deep L5 extratelencephalic-projecting (ET) neurons found in sensory cortices.^40,50^ Mapping projection targets onto the t-SNE plot of electrical features showed that intratelencephalic-projecting (IT) neurons (e.g., cortical- and amygdala-projecting neurons) and ET neurons (e.g., thalamic-projecting neurons) had distinctive electrical properties (Fig. 6c), adding evidence to this notion observed in the primary cortices.^40,50^ Interestingly, although corticothalamic MD- and PF/PO-projecting neurons were located in overlapping cortical layers (Extended Data Fig. 7b,c), we found that only PF/PO-projecting neurons, but not MD-projecting neurons, resembled the aforementioned L5b neurons defined by their morphology and physiology, which might contribute to the functional differences of these two thalamic nuclei (Fig. 6b,c, and Extended Data Fig. 8d).

**Fig. 6:**
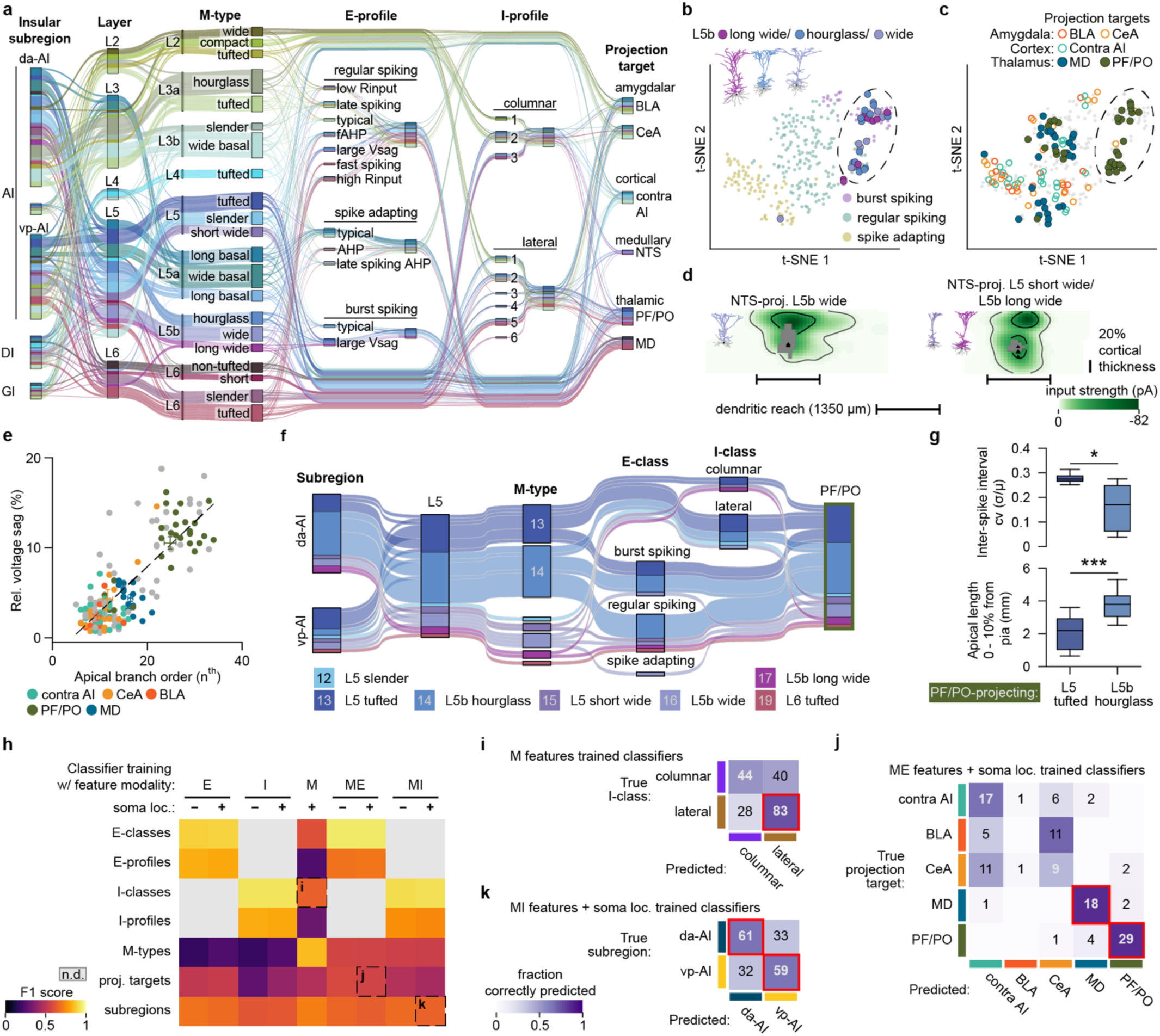
Multimodal analyses distinguish and predict specific insular cell types. **a**, Parallel categories diagram of all neurons and their allocations to subregions, layers, morphology-class (M-classes, based on predominant layer and cluster hierarchy from Fig. 2e), M-types, E-profiles, I-profiles, and projection targets. **b** and **c**, M-type (**b**) or projection target (**c**) identities mapped onto the t-SNE plot from Fig. 4c. Dashed ellipses highlight a group of L5b neurons with similar electrical properties that primarily project to the PF/PO. **d**, Two examples of averaged local input maps with either prominent columnar or lateral connectivity based on specific cell types defined by other modalities. The average maps are Gaussian-filtered for visualization, shown in an exponential color scale to highlight distal inputs, and contoured at 15^th^ and 85^th^ percentiles of all responses. Black lateral whisker bars: lateral boundary thresholds based on dendritic reach (1350 µm); gray pixels: direct response sites; black triangles: soma locations. **e**, Relative voltage sag as a function of apical branch order (Spearman correlation = 0.70), with projection target labeled. Dashed line: linear fit. **f**, Parallel categories diagram of L5 PF/PO-projecting neurons and their allocations to subregions, M-types, E-classes, and input classes (I-classes). **g**, Quantification of electrical (top, inter-spike interval cv, Mann-Whitney U test U = 44, p = 0.013) and morphological (bottom, apical length within 10% cortical thickness bin from pia, Mann-Whitney U test U = 20, p < 0.001) characteristic features for PF/PO-projecting *L5 tufted* (M-type 13) and *L5b hourglass* (M-type 14) neurons. **h**–**k**, Performance of random forest classifiers trained on specific uni- and multimodal feature sets to predict classes and types across different modalities. **h**, Classifier accuracy (F1 score, Methods). The F1 scores for classifier predictions of I-classes (**i**), projection targets (**j**), or subregions (**k**) are highlighted in (**h**) (dashed lines). **i**, I-classes predicted by M features. **j**, Projection targets predicted by morpho-electrical (ME) features and soma location. **k**, Subregions predicted by morpho-input (MI) features and soma location. For (**i**–**k**), purple luminance: percentage of neurons correctly predicted by classifiers for a given I-class, projection target, or subregion; red boxes: cell types well predicted by classifiers. da: dorsoanterior; vp: ventroposterior; BLA: basolateral amygdala; CeA: central amygdala; contra AI: contralateral agranular insula; MD: mediodorsal thalamus; NTS: nucleus tractus solitarius; PF/PO: parafascicular/posterior thalamus; loc.: location; n.d.: no data; proj.: projection.

Several morphological types and projection targets also showed prominent association with specific local input patterns (Fig. 6d and Extended Data Fig. 11b–e). First, *L5 short wide* (M-type 15) and *L5b long wide* neurons (M-type 17) received exclusively columnar inputs (I-profiles 2 and 3), whereas *L6 tufted* neurons (M-type 19) associated overwhelmingly with lateral inputs (I-profile 5) (Extended Data Fig. 11b,d). Second, IC neurons projecting to the PF/PO and MD thalamus, as well as the BLA, but not the CeA, contralateral AI, or NTS, displayed a significantly higher frequency of lateral inputs, compared to null (‘unknown’) neurons (Extended Data Fig. 11c; Fisher’s exact test p_PF/PO_ = 0.010, p_MD_= 0.040, p_BLA_= 0.021, p_CeA_= 0.111). Guided by our visualization (Fig. 6a), we found that remarkable preferences for columnar or lateral input patterns could also be deduced from combinations of morphology with projection target or morphology with subregion location. Notably, NTS-projecting neurons could be divided into two groups for their local inputs, depending on their morphology (Fig. 6d). These results indicate that pyramidal neurons defined by morphology and projection target may receive specific columnar or lateral inputs.

#### Specific insular cell types are separated by individually salient features

One advantage of our iCCA approach is that it is based on interpretable biologically-relevant features that quantify neuronal properties underlying a cell type. The availability of these quantitative features allowed us to address two fundamental questions: how do structural properties (e.g., morphology, projection) and functional properties (e.g., intrinsic electrical properties and local synaptic inputs) correlate, and what properties distinguish a cell type? First, we found multiple pairs of cross-modal features with significant monotonic association (Fig. 6e and Extended Data Fig. 12; Methods). For example, the relative size of voltage sag correlated with maximal apical branch order (Fig. 6e and Extended Data Fig. 12a), consistent with reported HCN channel gradients along apical dendrites of cortical neurons.^60^ This correlation also revealed a clear separation of PF/PO-projecting neurons from other projection neurons by their large complex dendritic arbors and strong voltage sags (Fig. 6e; see also Figs. 3d, 4d, 6b,c and Extended Data Fig. 7d). Beyond this example, morphological features also correlated with electrical and input features within all layers except L2 (Extended Data Fig. 12), indicating that the size and complexity of dendritic morphology may influence cellular physiology and local inputs. Second, we compared properties between cell types and identified salient features that distinguished them (Supplementary Fig. 3; Methods). For example, although both *L5 tufted* (M-type 13) and *L5b hourglass* (M-type 14) PF/PO-projecting neurons received strong lateral L3 inputs (Extended Data Fig. 13a), these two cell types were distinguishable by spiking properties and apical tuft sizes (Fig. 6f,g). Similarly, L3b–L5a amygdala-projecting cell types in different input classes were distinguishable by apical tuft length (Extended Data Fig. 13b–d).

#### Multimodal features predicted specific insular cell types

To quantify the contributions of different modalities to cell typing, we performed iCCA combining morphological features (Extended Data Fig. 5e) with either electrical (Extended Data Fig. 8b) or input features (Extended Data Figs. 10b, 14, and 15 and Extended Data Table 1; Methods). From this analysis, the morphology features turned out to be most informative in determining cell types, evidenced by their top rankings in the Gini-index during classifier training (Extended Data Figs. 14b and 15b). Interestingly, in contrast to primary cortices,^35^ using morpho-electrical features resulted in rather heterogeneous neuronal groups, meaning each group was often consisting of more than one M-type or E-profile, suggesting less co-variance between morphology and physiology categories compared to observations in the visual cortex (Extended Data Fig. 14g,h). Similar heterogeneity was observed when morpho-input pattern features were used in relation to morphology and input patterns (Extended Data Fig. 15h,i).

Practically, is a limited set of features sufficient to infer cell type identity based on the modality correlations observed above, particularly in combinations involving morphology? We trained independent instances of random forest classifiers with our interpretable features used in iCCAs to quantitatively assess whether they accurately assign neurons to their known types (M-types, E- and I-profiles and classes, subregions, and projection targets; Fig. 6h–k; Methods). We identified cross-modal classifiers that could predict a subset of cell types (Fig. 6h), although not all cross-modality-trained classifiers assigned cell types accurately. Prominently, classifiers trained on the feature sets that included morphology alone accurately predicted whether a cell received lateral inputs (Fig. 6i). When electrical or input features were combined with morphology, trained classifiers accurately predicted thalamic-projecting neurons (Fig. 6j) or subregion location (dorsoanterior-AI vs. ventroposterior-AI; Fig. 6k), respectively. These classifiers, along with high-performing within-modality classifiers (Fig. 6h) and the visualization tool (Fig. 6a), serve as valuable platforms for future work to predict cell type identity when certain modalities are unavailable and to link insular cellular properties to their functions.

### Posterior-to-anterior local inputs bridge a thalamo-cortico-amygadalar long-range circuit

To examine how IC cell types and their properties contribute to functional brain networks in the IC, we leveraged the surprising finding that cell types with asymmetrical laterally-extending input profiles (Fig. 5b,d) associated with specific projections (Extended Data Figs. 11c,e and 15g). This association raised the possibility that these local connections could integrate inputs and outputs of the functionally distinct anterior and posterior insular subregions. Behaviorally, the parvicellular ventroposteromedial thalamus (VPMpc) and amygdala are both required for food aversion.^61^ However, the complete circuits bridging these two functionally related regions are unknown. Anatomical evidence showed that the VPMpc and the nearby PF thalamus preferentially sent axons to the posterior IC whereas amygdala-projecting neurons were located in the anterior IC (Extended Data Fig. 16a–d).^5,7,62^ We hypothesized that a posterior-to-anterior intra-insular connection could be the missing link between the thalamic nuclei that relay gustatory and interoception information and the amygdala which mediates valence and behavioral choices based on such sensory information.

To test the functional connection of this circuit, we injected channelrhodopsin (ChR2) into the VPMpc/PF thalamus and performed loose-patch recordings of posterior IC pyramidal neurons while photostimulating VPMpc/PF axons. Indeed, VPMpc/PF projections drove action potentials in the posterior IC (Extended Data Fig. 16e,f; n = 14/29 neurons; see also Stone et al.^63^). To test whether VPMpc/PF signals are sent anteriorly via posterior-to-anterior local inputs, we recorded neurons in the anterior IC while performing channelrhodopsin-assisted circuit mapping^64^ in the posterior IC (Fig. 7a,b). A majority of neurons in the anterior IC responded strongly to optogenetic stimulations in the posterior IC, and minimally to stimulations in the anterior IC (Fig. 7c,d; anterior response < 10% of posterior response, n = 13/24 neurons). This result indicated that anterior IC neurons rarely received VPMpc/PF input directly, and instead, required posterior-to-anterior local inputs. The poly-synaptic nature of this thalamo-anterior IC circuit was supported by the delayed onset of anterior IC responses to VPMpc/PF stimulation (Extended Data Fig. 16g,h). In addition, the anterior-posterior distance between the responding anterior IC neurons and the VPMpc/PF axon stimulation sites in the posterior IC was within the range of observed posterior-to-anterior local input distances (Fig. 5b and Extended Data Fig. 16i). Thus, we concluded that the lateralized local inputs within the IC sent VPMpc/PF signals from the posterior to the anterior IC. Finally, we tested whether anteriorly located neurons receiving these poly-synaptic thalamic inputs project to the BLA. Indeed, IC neurons labeled by retrograde bead injections into the BLA responded preferentially to posterior photostimulation compared to nearby bead-negative neurons (Fig. 7e,f), indicating that interoceptive and gustatory thalamic signals were integrated and targeted toward the BLA via posterior-to-anterior local connections within the IC. Altogether, lateral, posteriorly-biased local inputs within the IC were required to connect a thalamic-cortical-amygdalar circuit linking the functionally-related IC subregions, VPMpc/PF thalamus, and BLA (Fig. 7g), providing physiological evidence for circuit substrates that may convert thalamic sensory signals arriving in the posterior IC to the amygdala-projecting cognitive signals in the anterior IC.

**Fig. 7:**
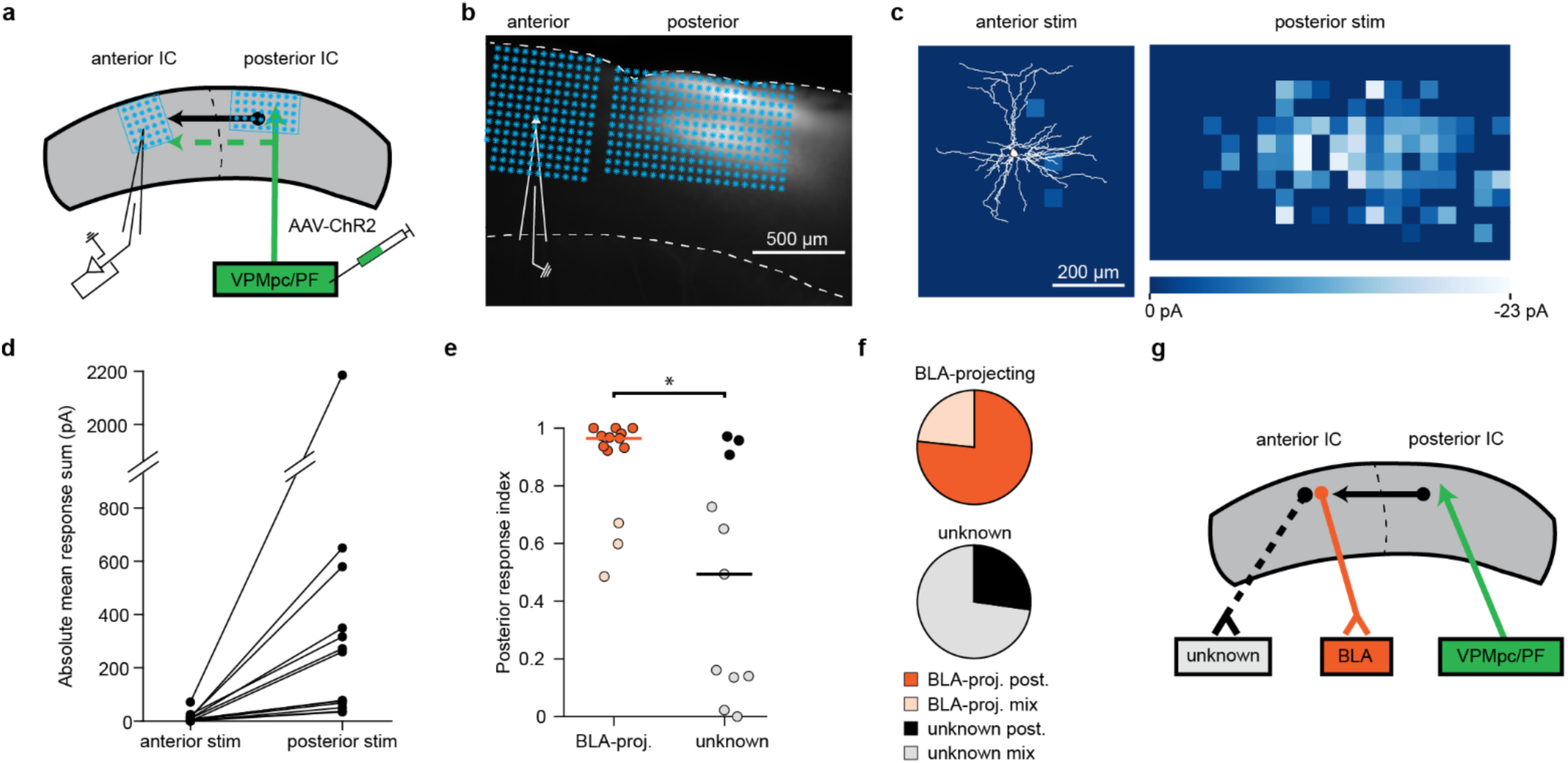
Posterior-to-anterior local inputs bridge a thalamo-cortico-amygadalar long-range circuit. **a** and **b**, Schematic (**a**) and example horizontal acute slice (**b**) of a recording in the anterior IC in response to ChR2 stimulations of VPMpc/PF axons in either the anterior or posterior IC. Blue dots: 50 × 50 µm grid of stimulation locations in either anterior or posterior IC. **c**, Example CRACM maps corresponding to anterior (left) and posterior (right) stimulations shown in (**b**). Reconstructed dendritic morphology (left) of the recorded neuron is overlaid. **d**, Quantification of CRACM maps in response to anterior vs. posterior stimulations. Sum of responses from neurons responding strongly to posterior stimulation and minimally to anterior stimulation are shown (n = 13) (anterior response < 10% of posterior response, Methods). **e**, Posterior response index of BLA-projecting neurons vs. neurons with unknown projection targets. Horizontal bars: median; dark color circles: neurons with posterior response index > 0.9; light color circles: posterior response index ≤ 0.9 (n = 13 for BLA-projecting neurons; n = 11 for neurons of unknown projection targets). *: p < 0.05 (Fisher’s exact test). **f**, Distribution of response types shown in (**e**). **g**, Schematic of the VPMpc/PF–IC– BLA long-range circuit. CC: corpus callosum; BLA-proj.: BLA-projecting; post: posterior; stim: stimulation.

## Discussion

Our multimodal dataset, consisting of over a thousand neurons, provides a comprehensive view of the cellular and circuit layout of the insula and sets the foundation for future functional studies. The cell typing analyses revealed several remarkable aspects of insular neurons, including large and elaborated dendrites, the anterior-posterior topographic distribution of M-types, layer-specific spike pattern distributions, and lateralized local inputs, along with the associations of these properties with specific projection targets. The quantitative anatomical analyses revealed dorsoanterior- and ventroposterior-AI subregions defined by topographical distributions of M-types and their associations with cell types, which allow users to identify locations most likely to contain a given type. Of note, our dorsoanterior-AI coordinates generally corroborate with the empirical anterior IC,^14,26^ while our ventroposterior-AI coordinates are anteriorly-shifted compared to the empirical posterior IC.^15^ This difference likely arose from the fact that the center of mass for the AI will be more anterior to the center of the mass of the entire IC (AI/DI/GI combined), and the latter has often been used for in vivo studies due to technical constraints in separating the AI/DI/GI subregions. Future work could harness the power of our anatomical framework and trained classifiers to both map and categorize cells using information obtained from, e.g., imaging or physiology experiments, to gain insights into putative circuit functions. As a representative application of our dataset, we identified that BLA-projecting neurons preferentially mediate long-range thalamus-to-amygdala connectivity via lateralized intra-insular microcircuits.

### Insular cell types and cross-modal associations

Cross-modal iCCA analyses and cross-tabulations of cell types highlighted both the broad heterogeneity in categorical cross-modal associations throughout the whole dataset and the existence of associations between specific cell types. The former prompts the hypothesis that modalities such as electrical and input properties may not be fixed for every given cell type defined by morphology, but rather could vary based on brain states. Despite the heterogenous cross-modal associations between M-types and E- or Ι-profiles, groups of M-types did occupy confined regions in the electrical and input feature space. For example, all L5b morphologies displayed large voltage sags with either burst or regular spiking behaviors (Fig. 6b). Such cross-modal associations were rooted in specific biological parameters: the more elaborate the distal dendritic arbor, the larger the voltage sags (Fig. 6e). This result points to the functional significance of cellular structures, because distal dendrites preferentially host HCN channels,^60^ and the regulation of HCN channel density is an important factor for synaptic integration at the dendritic level.^65^ Further, such physiological properties can be dynamic and modulated by neuromodulators, including dopamine, noradrenaline, acetylcholine, and serotonin.^66,67^ In addition, cell type-specific modulation of input strength by factors including neuromodulators may fine-tune local input distributions.^66,67^ Therefore, the E- and I-profiles obtained from ‘naïve’ mice in the present work may reflect the baseline electrical and input states from which cells can transition in response to neuromodulation.

With the emergence of cell typing datasets,^35,36,38,68^ it is imperative to perform systematic meta-analyses^69^ to understand common and subregion-specific cortical cell types and local inputs across cortical regions and species. Curiously, we did not observe VEN-like morphologies reported in other species including hominids despite our sampling depth, confirming previous surveys.^43,70^ However, we found a subset of *L5a wide basal* neurons (M-type 8) that have two apical dendrites, which could potentially be analogous to the “fork cells” found in the human insular cortex.^70,71^ Additionally, no stellate cells were identified out of the 37 L4 neurons recorded in the DI and GI, contrasting with their sparse or predominant presence in L4 of the rodent visual or somatosensory cortex, respectively.^35,46,48^ The growing availability of transcriptomic datasets,^72,73^ which often now include morphological information,^74–76^ provides platforms for linking all modalities presented here to transcriptional information using morphology. Practical challenges remain in the stratification of metadata and cell typing strategies (e.g., feature extraction and selection) across datasets.^77,78^ Therefore, our atlas-registered dataset with a reproducible clustering pipeline will facilitate future work reconciling existing whole-brain transcriptomic types^72^ with our multi-modal insular cell types.

### Characteristics of local insular computation

From an anatomical perspective, distinctive yet continuous AI, DI, and GI subregions piqued our interest in how information flows within and between these subregions. The AI, despite lacking a Nissl-defined L4, sparsely hosts neurons reminiscent of L4 morphologies (M-types 6, 7, and 10; Fig. 3d). Whether these neurons function as bona fide L4 neurons in an agranular cortex^79^ invites future efforts to test whether they express the L4 marker *Rorb,*^79^ send ascending inputs to L2/3, and receive thalamic inputs.^33,34^ Interestingly, strong descending inputs from L2/3 to L5 are conserved among the AI, the agranular motor cortex,^54,57^ and canonical granular cortices,^54,58^ but the strong ascending inputs to L2/3 observed in granular cortices^54,55,58^ were not observed in the agranular motor cortex^54,57^ or the AI. Although our dataset prioritized AI microcircuits, we did identify DI/GI to AI inputs, in line with previous anatomical and functional studies.^3,22,24^ Future studies on DI/GI microcircuits will reveal whether IC subregions with a six-layered organization employ distinct local computations compared to the AI, and whether IC subregions with different laminations connect as theorized.^80^

From a circuit perspective, one striking feature of the IC is that its lateralized local inputs (Fig. 5d) extend well beyond the width of canonical cortical columns.^81,82^ Although out-of-column cortical connections have been implied by axonal reconstructions and in vivo recordings,^83,84^ our dataset provides direct physiological evidence for the extensive lateral reach of intra-insular integration across subregions along its anterior-posterior axis, distinguishing the IC from other cortices.^53,54,56,57,59^ Our data also validated the long-sought connections between the anterior and posterior IC previously extrapolated from in vivo data,^1,20^ and demonstrated its necessity in connecting visceral and valence-related thalamic and amygdalar regions that communicate via the insula (Fig. 7).^25,26,61^ Because these inputs preferentially arrived from the posterior IC, this result aligned with the tract-tracing derived hypothesis that information flows from superficial layers to deep layers intracortically toward higher functional hierarchy^80,85^ and provides physiological evidence for the functional theory that the posterior IC sends feedforward information anteriorly.^1,10^

### Limitation and future directions

Several aspects beyond the scope of the current study can further enhance the comprehensiveness of our multimodal insular atlas. We usually recorded features across two or more modalities for each neuron, but never across all modalities simultaneously due to technical constraints and we used morphology as a gateway to connect to other modalities. In addition, we demonstrated one example of using our dataset to facilitate circuit mapping; more applications of our dataset can be carried out by future studies.

## Methods

### Animals

All procedures were approved by the Oregon Health & Science University Institutional Animal Care and Use Committee (IACUC). Male and female wildtype C57BL/6 mice (Charles River Laboratories, Wilmington, MA, USA) and PV-Cre, Sst-Cre, Gad2-Cre, Drd1-EGFP, Dat-Cre, Ai32, and Ai9 mice—all on a C57BL/6 background—were used (Tables S3 and S4). PV-Cre+/-;Ai9+/- mice were used to aid demarcation of the insula subregions during the establishing phase of the study. Mice were housed in group housing, given free access to food and water, and maintained on a 12-hour light/ dark cycle. Animal ages are specified in the following experiments.

### Atlas collection and histology

Four wildtype and six *PV-Cre^+/-^;Ai9^+/-^* mice (P49 ± 1 d, equal sex) were used to develop a reference atlas of the insular cortex. Mice were transcardially perfused with ∼50 mL 4% paraformaldehyde (PFA) in 0.1 M phosphate buffer (PB, pH 7.2). Each brain was dissected from the skull and post-fixed overnight in 4% PFA, then stored in phosphate buffered saline (PBS, pH 7.4) until sectioning. All subsequent steps were conducted at room temperature unless otherwise stated. To prepare for sectioning, each brain was dry-blotted, and the dorsal surface superglued onto a wedge (10° tilt, 2.5% agarose or aluminum) with the anterior side tilting upwards. This slicing method was used throughout the study to preserve the functional anterior-posterior axis of the insula as well as its dendritic morphologies and is referred to as ‘modified horizontal’ sectioning for simplicity. The brain was then embedded in 3% agarose, blocked, then sectioned in this modified horizontal orientation at 40 µm in PBS on a vibrating microtome (VT1200S; Leica Biosystems, Wetzlar, Germany). Alternating sections were either directly mounted onto slides (Superfrost Plus; Thermo Fisher Scientific, Waltham, MA, USA) for Nissl staining, or collected as free-floating sections in welled plates for parvalbumin (PV) immunohistochemistry.

Nissl staining was performed following a standard protocol^86^ and stained slides were coverslipped with Eukitt (#03989; Sigma-Aldrich, Burlington, MA, USA). For PV immunohistochemistry, sections were first rinsed 3× 10 min in Tris-buffered saline (TBS), then incubated with 3% H_2_O_2_ + 10% MeOH in TBS for 30 min to quench endogenous peroxidase activity, followed by another 3× rinse. Sections were then blocked for 1–2 h with 5% goat serum in TBS + 0.1% Triton X-100 (Sigma-Aldrich), then incubated overnight at 4 °C with mouse anti-PV primary antibody (1:1000; P3088; Sigma-Aldrich) in the blocking solution. The next day, sections were rinsed 3×, then incubated for 1–2 h with HRP-conjugated goat anti-mouse secondary antibody (1:500, #115-035-003; Jackson ImmunoResearch Laboratories, Inc., West Grove, PA, USA) in the blocking solution. Sections were again washed 3×, then incubated for 90 min with ABC (Vectastain Elite ABC-HRP Kit, PK-6100; Vector Laboratories, Newark, CA, USA), washed 3×, developed with nickel-intensified 3,3’-diaminobenzidine tetrahydrochloride (DAB, #5905; Sigma-Aldrich), and washed a final time. Lastly, sections were washed in 0.05 M PB, mounted on microscope slides (Superfrost Plus), and coverslipped with Eukitt.

### Atlas microscopy and annotation

All slides were imaged on an Axio Scan.Z1 (Zeiss, Oberkochen, Germany) using a 20× objective. Image data were subsequently processed in FIJI^87^ using custom scripts. Each brain was first reconstructed in 3D by importing ×16 downsampled images in serial order into TrakEM2,^88^ where they were semi-automatically aligned across the cutting axis (Supplementary Fig. 1 and Extended Data Fig. 1b). This operation was completed separately for Nissl- and PV-stained slides. For later steps, a merged atlas was also created for each brain by interleaving the two histological datasets in serial cutting order and applying a 4% enlargement to PV sections to correct for the average difference in shrinkage artifact between the two methods. All stained slide series were manually inspected for histological quality—i.e., absence of shrinkage and staining artifacts, and contiguous undamaged sections containing the insula. No qualitative differences were observed between *PV-Cre;Ai9* and wildtype samples.

For Nissl reconstructions, the three highest-quality *PV-Cre;Ai9* specimens were selected for further processing (Supplementary Fig. 1). Each section was manually annotated in TrakEM2 using a graphics tablet (Artist 22R Pro; XP-Pen, Shenzhen, China, or Cintiq 22HD; Wacom, Kazo, Japan) to segment the following conspicuous cortical features: (1) the pial surface, (2) the border between cortical layers 1 and 2 (L1/L2), (3) the superficial edge of the corpus callosum (CC), and (4) the pial indentation(s) corresponding to the middle cerebral artery (MCA) (Fig. 1 and Supplementary Fig. 1). Next, a 3D bounding box that contained the entirety of the right insular cortex was established to spatially restrict subsequent analyses. Using the rigid affine transforms established by alignment in TrakEM2, a full-resolution region of interest (ROI) was extracted from each section that intersected with this bounding box, thus creating a spatially coherent image stack through the insular cortex (Fig. 1b and Supplementary Fig. 1). For each section in the stack, a custom script utilizing the MorphoLibJ library^89^ automatically segmented the tissue from the slide background with occasional minor manual corrections when needed, and the image contrast was globally enhanced (normalized) using intensity data within the tissue mask.

Each image stack was loaded in Photoshop (v.23; Adobe Inc., San Jose, CA, USA) where two expert annotators each independently delineated the boundaries of cortical layers by painting a color-coded overlay using the Pencil tool. Subregional bounds (e.g., AI/DI border) were indicated (Supplementary Fig. 1). A histological atlas^45^ was used as a visual reference during annotation. A final consensus was reached through an averaging routine: identically annotated pixels were preserved by taking the intersection of the two overlays, after which an operational loop expanded each color-coded region into unallocated space by 1 pixel per iteration until the entire ROI bounded by the pial, CC, and outer-most subregional annotations was filled with coded pixels (Fig. 1c and Supplementary Fig. 1).

### Laplacian analysis

To register Nissl atlas locations into a reference coordinate space that accounts for the variable curvature and cortical thickness of the insula, we employed a Laplacian operator. Laplace’s equation was solved for the scalar field *ψ* enclosed between the two lines defined by annotations of the CC and L1/L2 boundaries described above in Nissl sections.^90–92^ It takes the form:

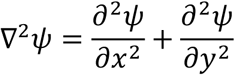

where *x* and *y* are the standard Cartesian coordinates in the Nissl section, ▽ the Laplacian gradient of the scalar field, and ∂ the partial derivatives of the Cartesian coordinates. The L1/L2 boundary was used in lieu of the pial boundary because the latter was often subject to minor deformations and folds during tissue mounting that could misrepresent local curvature. Laplace’s equation was iteratively solved using the Jacobi method, where the CC and L1/L2 boundaries were always fixed to 0.0 and 1.0, respectively, with standard iterative (*i*) relaxation taking the form:

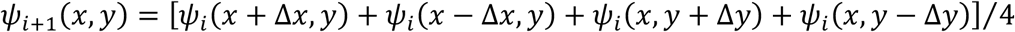

Following Lerch et al.,^91^ an additional “resistive” boundary was imposed to account for the fact that the CC and L1/L2 boundaries do not intersect. All pixels outside of the scalar field *ψ* enclosed between these lines were fixed to *NaN*, and with each iteration, calculations for edge pixels of *ψ* discounted contributions from any adjacent null pixels. The final equidistance field was then iteratively extrapolated in 1-px steps following two-point gradients until the entire ROI bounded by the pial and CC annotations was filled (Fig. 1d). Streamlines (i.e., steepest gradient descent paths) were then computed at any point within *ψ* between the pia and CC and were used to estimate local columnar orientation and cortical thickness.

### Insular flatmap

We transformed each brain into a 2D flatmap that topographically organized data with respect to (1) relative section along the cutting axis and (2) relative position along the laminar axis, which was defined as the mid-value (= 0.5) isocontour line of the Laplacian gradient *ψ* (Fig. 1d and Extended Data Fig. 1b,c) and approximated the anterior-posterior axis. To align data across the cutting axis in flatmap coordinates, a common reference was identified in the following way that could be located at a specific point along the laminar axis of each section. Surface reconstructions of each right hemisphere were visualized with ParaView (v5.10; Kitware, Inc., Clifton Park, NY, USA). Using affine transforms obtained by solving linear systems— each containing 21–25 matched fiducial coordinates between pairs of brains—we aligned these surfaces within a common 3D reference frame (Extended Data Fig. 1a). Applying these transforms to the MCA annotation coordinates revealed remarkable consistency in the placement of the trunk of the MCA proximal to its bifurcation across all three animals (Extended Data Fig. 1a). We therefore established a common reference as follows: for each mouse, take the MCA annotation coordinate from each section that falls within 1 mm ventral (proximal) to its point of bifurcation (n = 12 sections; Extended Data Fig. 1a,b). Second, calculate the streamline that intersects with the MCA annotation of each section, and take the coordinate of the intersection of this streamline with its respective laminar axis. Third, determine the plane that best fits these 24 coordinates (2 per section × 12 sections) in 3D space using singular value decomposition (Extended Data Fig. 1b). The point of intersection between this plane and the laminar axis of each section was defined as the section’s coordinate origin (Extended Data Fig. 1b,c, white circles), and all points/streamlines within a section were therefore plotted onto the flatmap in terms of their relative distance to this origin along the laminar axis. Cortical thickness on each flatmap was generated by sampling streamline lengths at 25-µm intervals along the laminar axis for each section across the cutting axis (Extended Data Fig. 1c,d).

To define the subregional bounds of the insula flatmap—anterior edge, AI/DI border, DI/GI border, and posterior edge—the intersections of each annotator’s lines with the laminar axis were identified, converted to relative positions along this axis, and averaged (Fig. 1b,c and Supplementary Fig. 1, black dashed; Extended Data Fig. 1d, white dashed). The coordinates of the cutting axis were also adjusted to make them relative to the ventral-most section that contained an annotation for L4 (Supplementary Fig. 1, dashed box; Extended Data Fig. 1d, white diamonds). The orientation of the long axis (Extended Data Fig. 1d, black dashed) for each insular flatmap was defined as the angle between the cutting plane and the major axis of an ellipse fit to the outermost boundaries of the insular flatmap; the short axis was orthogonal to this angle. This ellipse was calculated as the area having the same second order central moments as the insular flatmap region.

To compare data across brains, each flatmap was rotated about its 2D origin (defined by the L4/MCA origin; Extended Data Fig. 1d, white diamonds) such that the long axes were parallel across flatmaps and overlaid (Extended Data Fig. 1f). Average laminar boundaries were generated as follows: for each subregional boundary, coordinates from the three individual aligned maps were binned and averaged at 250-µm intervals along the cutting axis, then smoothed with a cubic spline (Extended Data Fig. 1g). The average insula was defined as the area between the average anterior and posterior bounds along the laminar axis. The average GI was defined as the area between the average DI/GI border and the posterior edge. The average DI was defined as the area between the average AI/DI border and the posterior edge, minus the GI area. The remaining area defined the average AI. An equivalent operation was applied to the error bounds (± 1 standard deviation) of each edge (Extended Data Fig. 1g, gray shading).

### Cortical layer analysis

Layer annotations were transformed to the flatmap space using streamlines spaced at 25-µm intervals along the laminar axis, as above (Fig. 1d and Extended Data Fig. 2). The pixel color/values of each annotated painting were queried at 0.1-µm steps along each streamline, yielding the thickness of each cortical layer at a given laminar position (Extended Data Fig. 2a,b). Dividing these values by the total length of each streamline (i.e., cortical thickness) produced normalized values. Thicknesses for all sections were then assembled within the 2D flatmap to highlight topographic variation (Extended Data Fig. 2c).

As each reference mouse brain required a slightly different rotation to its flatmap (Extended Data Fig. 1f), the individual 2D sample grids (Extended Data Fig. 2c) did not precisely overlap when overlaid upon one another. Therefore, a grand 2D average of normalized thickness for each layer was computed by binning and averaging values from each mouse brain at 100-µm and 250-µm intervals along the long and short axes, respectively, relative to the MCA origin, and finally averaging across mice (Extended Data Fig. 3a). This operation was run separately for bins assigned to AI or DI/GI (Extended Data Figs. 2c and 3a) due to the presence or absence of L4 annotations.

### Stereotaxic injections

Wildtype C57BL/6 mice (P21–P35) were anesthetized with 2% isoflurane and injected with adeno-associated virus (AAV) and/or fluorescent retrograde microbeads (LumaFluor, Inc., Durham, NC, USA) as previously described.^44^ For retrograde labeling of AI output neurons, 30 nL red retrograde beads (1:1 diluted in filtered PBS, Red Retrobeads IX) or green retrograde beads (1:1 diluted or undiluted, Green Retrobeads IX) were injected in up to two selected projection target regions per animal, namely contralateral AI, BLA, CeA, MD, PF/PO, and NTS. For retrograde labeling of thalamic nuclei projecting to the posterior AI, 30 nL red retrograde beads (1:1 diluted in filtered PBS, Red Retrobeads IX) was injected in the ipsilateral posterior AI. For optogenetic experiments, 30–40 nL of AAV2/1 ChR2-Venus (#20071; Addgene, Watertown, MA, USA) or AAV2/1 hChR2-TdTomatoH134R (lot V0314; Penn Viral Core, Philadelphia, PA, USA; also available from Addgene, #28017) virus was injected in the VPMpc/PF thalamus. Note that due to technical limitations, thalamic injections often targeted more than one nucleus. PF/PO injections were aimed at the PF, and partially labeled the adjacent PO region; VPMpc/PF injections were aimed at both VPMpc and PF, and sometimes led to minor spillover in neighboring nuclei (Extended Data Fig. 16c) which was negligible because they did not project to the IC (Extended Data Fig. 16a,b). Injection coordinates are measured in µm relative to bregma (unless noted otherwise) along anterior-posterior (AP), medial-lateral (ML), and dorsal-ventral (DV) axes as follows: contralateral AI (AP:1350 – 1700, ML: −2850 – −2900, DV: 3700 – 3800), BLA (AP: −1350, ML: 3100, DV: 5100), CeA (AP: −1050, ML: 2600, DV: 4750), MD (AP: −1050, ML: 250 – 300, DV: 3400), PF/PO (AP: −2050 – −2175, ML: 550 – 600, DV: 3400), NTS (AP: −6200, ML: 1150, DV from brain surface: 3900), posterior AI (AP: 750, ML: 3550, DV: 3900), and VPMpc/PF (AP: −2050 – −2175, ML: 550 – 600, DV: 3900).

### Electrophysiology recordings

For intrinsic property experiments, brain slices were prepared from P21–P107 animals (median = P38 for animals included in electrical feature-based iCCA, median = P40 for animals included in morphological feature-based iCCA); for LSPS, brain slices were prepared from P29–P59 animals (median = P38). Brains injected with retrograde beads and/or virus at P21–35 were sliced 3–14 days after bead injection or 14–28 days after viral infection of ChR2-expressing AAV. Wildtype C57BL/6 or *PV-Cre*^+/-^;*Ai9*^+/-^ mice were transcardially perfused with artificial cerebrospinal fluid (ACSF) cutting solution (in mM: 127 NaCl, 24 NaHCO_2_, 1.25 NaH_2_PO_4_·H_2_O, 25 glucose, 2.5 KCl, 2 CaCl_2_, 1 MgCl_2_) and decapitated to obtain modified horizontal brain slices as described above (VT1200S). Perfusion and slicing ACSF temperature was matched to subsequent slicing protocol. For intrinsic property recordings, brains were sliced at 300 µm thickness with warm (∼34 °C) ACSF cutting solution. For LSPS and ChR2 experiments, brains were sliced at 350 µm thickness with ice-cold choline cutting solution (in mM: 25 NaHCO_2,_ 1.25 NaH_2_PO_4_·H_2_O, 25 glucose, 2.5 KCl, 7 MgCl_2_·6H_2_O, 0.5 CaCl_2_; with 110 choline chloride, 11.5 sodium ascorbate, and 3 sodium pyruvate added freshly before the experiment). Whole-cell recordings were performed using a Multiclamp 700B amplifier (Molecular Devices, San Jose, CA, USA) with borosilicate pipettes (3–6 MΩ; Sutter Instruments, Novato, CA, USA) under an Olympus microscope (BX51W with 60×, 1.0 NA water immersion objective; Tokyo, Japan), and data were collected using Ephus software^93^ within MATLAB (Mathworks, Natick, MA, USA). A cesium-based internal solution was used for voltage-clamp recordings (in mM: 135 Cs-gluconate, 1 EGTA, 1.5 MgCl_2_, 10 HEPES, 2 Na_2_ATP, 0.3 GTP, 7.8 Na_2_·phosphocreatine, and 0.015 Alexa-594 (Invitrogen), adjusted to pH 7.35–7.4 using CsOH, ∼280 mOsm; with 3 QX-314·chloride (#2313; Tocris Biosciences, Bristol, UK) added before experiments), and a potassium-based internal solution was used for current-clamp recordings (in mM: 120 K-gluconate, 5 NaCl, 2 MgCl_2_, 10 HEPES, 1.1 EGTA, 4 Mg_2_ATP, 0.4 Na_2_GTP, 15 Na_2_·phosphocreatine, and 0.015 Alexa-594, adjusted to pH 7.25 using KOH, 290 mOsm). 3–5 mg/mL biocytin (B4261; Sigma-Aldrich) was included in the internal solutions for morphological reconstructions. For all experiments, brain slices were continuously perfused in ACSF (in mM: 127 NaCl, 24 NaHCO_2_, 1.25 NaH_2_PO_4_·H_2_O, 25 glucose, 2.5 KCl, 2 CaCl_2_, 1 MgCl_2_) with NMDA receptor blocker (R)-CPP (5 μM; 02-471-0, Thermo Fisher Scientific). Intrinsic property recordings were performed at near-physiological temperature (30–34 °C); LSPS and ChR2 recordings were performed at room temperature (∼23 °C). All recordings were performed in the right hemisphere. At the conclusion of each recording a microphotograph was captured using a low-magnification objective (4×, 0.16 NA; UPlanApo, Olympus) used for the estimation of cortical soma position and shrinkage factor.

Intrinsic property recordings consisted of three current-step protocols during which sweeps were recorded at 5 s inter-sweep intervals: (1) A current-voltage protocol with repetitive sweeps containing 200-ms-long square pulse current steps starting at −300 pA with 25 pA increments every fourth sweep until 625 pA was reached, or spike generation was occluded by depolarization-block. (2) A ramp-protocol with stimulation sweeps consisting of a 3-s-long rising current injection at a rate of 165 pA/s, repeated for ten sweep iterations. (3) A spike train-protocol with repetitive sweeps containing 1-s-long current steps starting at 0 pA with 50 pA increments every fourth sweep until 1050 pA was reached, or spike generation was occluded by depolarization-block.

### Biocytin labeling and staining

During whole-cell recordings, neurons were filled with internal solutions (see above) containing 3–5 mg/mL biocytin and subsequently visualized using DAB staining (adapted from Marx et al.^94^). Briefly, after recording, slices were fixed for 12 hrs in a 4% PFA solution and stored in PB until staining. Following 6 × 10 min washes in PB, slices were incubated 2 × 20 min in 3% H_2_O_2_ in PB solution to quench endogenous peroxidases. Slices were washed 4 × 5 min in PB and incubated for 1 hr at room temperature, followed by overnight incubation at 4 °C, and then again at room temperature for 1 hr the following day in PB containing 1% Vectastain ABC reagent (Vector Laboratories) and 0.1% Triton X-100 (Sigma-Aldrich). After 6 × 10 min washes in PB, slices were incubated for 30 min in cobalt-and-nickel-intensified DAB in PB solution (2 μL CoCl_2_ and 4 μl (NH_4_)_2_Ni(SO_4_)_2_ per 1 mg/mL DAB) at 4 °C. Slices were developed using 8 μl of 3% H_2_O_2_ in PB solution per slice for 15 sec – 2 min before being washed 3 × 10 min in PB with the final wash containing a 1:5,000 dilution of Hoechst 33342 (H3570; Thermo Fisher Scientific). Slices were then mounted using spun-down Aqua-Poly/Mount (#18606-5; Polysciences, Warrington, PA, USA) media and the coverslip was sealed with clear nail polish (Super Shine; Sally Hansen, Morris Plains, NJ, USA). Slices were dried and settled for a minimum of 2 weeks before morphological reconstruction.

### Morphological reconstruction and processing

Biocytin-labeled neurons were viewed under an Axio Imager.M2 microscope (Zeiss) at 4× magnification to assess filling and staining quality, as well as to identify cell locations. Cells were inspected for intact apical dendrites that extended all the way to the pial surface (excepting some L6 cell types), a dark biocytin fill that persisted to the distal parts of both apical and basal dendrites, and the presence of dendritic spines to confirm an excitatory neuronal identity. Neurons were reconstructed manually using Neurolucida (MBF Bioscience, Williston, VT, USA) at 40× magnification with an oil objective (EC Plan-Neofluar 40×, 1.3 NA; Zeiss). Cell bodies, apical dendrites, basal dendrites, and axons were traced and annotated. All processes were traced to their termination points, a truncation point, or location where staining could no longer be visualized. Section contours were traced in 3D. The pial edge was traced along the top and bottom surfaces of each slice. Additional contours were traced along the outline of multiple reference structures (lateral ventricles, pial surface, corpus callosum, fimbria, fornix, internal capsule, and outline of the right hemisphere) as landmarks.

Individual reconstructed cells were isolated from original Neurolucida files (DAT) containing multiple cells on the same acute slice and converted to SWC+ files using the HBP Neuron Morphology Viewer.^95^ Individual SWC+ files were then processed in Python (BrainCellSuite Python package; https://gitlab.com/maolab/braincellsuite_submission) to extract cortical thickness, correct for post-fixation slice shrinkage, and to set the coordinate origin at the soma center. The slice shrinkage was calculated as the ratio between pia-soma distance in the biocytin-stained section and pia-soma distance prior to fixation. The average of this post-fixation slice shrinkage was 0.96 ± 0.14 (mean ± standard error) and 0.49 ± 0.15 (mean ± standard error) across the X/Y plane and Z-depth, respectively. 3D morphology plots and flattened 2D images were then generated from these processed files (BrainCellSuite). 3D morphology plots were used to check reconstruction quality. Cells with apical dendrites that were truncated due to slicing or were missing apical and/or basal dendrite annotations due to poor staining quality were excluded from the dataset. Processed morphologies were then used for feature extraction.

### Cell assignments

Reconstructed cells were matched to our histological atlas of the insula (Extended Data Fig. 4c), thus allowing us to estimate a flatmap coordinate for each cell that consists of: (1) slice position along the cutting axis, (2) anterior-posterior position along the laminar axis, and (3) normalized cortical depth. Following post hoc recovery of cell morphology (Extended Data Fig. 4e,f), each DAB-stained slice was inserted into the PV-atlas for “Mouse #2”. The PV atlas was used because it also utilized DAB processing, and thus appeared texturally similar to the recovered acute slices. As acute slices were thicker (300–350 µm) than atlas sections (40 µm, every other; see above), each acute slice was compared to a range of atlas sections before inserting it into the middle of the matched stack within the atlas’s TrakEM2 project. Within TrakEM2, the slice was manually aligned to the reference atlas section. We prioritized the matching of features (e.g., pial and CC edges, MCA, etc.) that were proximal to the labeled cells to minimize the effects of tissue distortions that can accumulate with histological processing. Each acute slice was independently matched to the atlas and had no bearing on the placement of other slices.

Once aligned to the PV-atlas reference frame (Extended Data Fig. 4g), the coordinate position for the soma of each recovered cell was noted, then shifted −40 µm along the cutting axis to place it within the adjacent, interleaved Nissl-atlas reference frame (Extended Data Fig. 4h, open circles), yielding the first flatmap coordinate component: slice position. A streamline was then calculated that intersected with the soma (Extended Data Fig. 4h, vertical dashed lines). The crossing of this streamline with the mid-value Laplacian isocontour (Extended Data Fig. 4h, horizontal dashed line) yielded the second flatmap coordinate component: anterior-posterior position. Lastly, the soma location was corrected by shifting it along the streamline such that its normalized pia-to-CC position matched the measurement obtained from the acute slice (Extended Data Fig. 4h, filled circles), yielding the third flatmap coordinate component: cortical depth. When possible, this depth value was measured from pipette position during electrophysiology (e.g., Extended Data Fig. 4d), otherwise it was obtained following morphological recovery (e.g., Extended Data Fig. 4e). The first two dimensions of the flatmap coordinate were interchangeable with the long- and short-flatmap axes using a change of basis based on the 2D rotation of the “Mouse #2” flatmap about the MCA origin (see above, slice and anterior-posterior positions). Using their estimated flatmap coordinates, we assigned each reconstructed cell to both an insular subregion and cortical layer (Fig. 1f,g and Extended Data Figs. 3 and 4). A handful of cells (14/1090) whose estimated topographic positions fell just outside the bounds of the average flatmap, yet remained within the error margins, were assigned to the nearest subregion within the same slice position (Fig. 1f).

### Imaging and quantification of retrograde fluorescent bead distribution

To quantify the distribution of labeled AI output neurons along cortical depth, brains injected in the contralateral AI, BLA, CeA, MD, PF, or NTS with retrograde beads were perfused, sliced at 50 µm thickness, stained with Hoechst 33342 (1:5,000 dilution), mounted in aqueous mounting medium (FluoroMount-G, Sigma-Aldrich), and tile-imaged with a 10× or 20× objective on an Axio Imager.M2 microscope (Zeiss). For each brain, bead-positive cells were manually counted in FIJI.^87^ The relative cortical depth of each bead-positive cell was calculated by normalizing its distance to pia over the average cortical thickness in a ∼0.3 mm wide rectangular ROI surrounding the cell that covered the full cortical depth. The normalized distribution of bead-positive neurons was then averaged across slices for each projection target (Extended Data Fig. 7c; n = 3 mice for each group). To identify thalamic projections to posterior AI (Fig. 7 and Extended Data Fig. 16a,b), brains injected in the posterior AI with retrograde beads were perfused, sliced at 100 µm thickness, mounted in aqueous mounting medium (FluoroMount-G), and imaged under the bright-field and red fluorescence channels on an Axio Zoom.V16 microscope (Zeiss).

### Laser scanning photostimulation (LSPS)

LSPS was performed as previously described.^96^ Briefly, MNI-caged-L-glutamate (200 μM, #1490; Tocris Biosciences, Bristol, UK) was uncaged in confined areas by 1 ms collimated ultra-violet laser stimulation (Series 3500; DPSS Lasers Inc., Santa Clara, CA, USA), with 20 mW light power measured after the objective (XLFLUOR2X/340, 0.14 NA; Olympus Corporation, Tokyo, Japan). For each map, stimulation sites were positioned across a 75 × 75 μm-spaced grid, with light pulses given at each site (Extended Data Fig. 9a; beam size: ∼60 µm, full width at half maximum light intensity) in pseudo-randomized order that maximized distance between consecutive stimuli using galvanometer scanners controlled by Ephus software.^93^ Postsynaptic pyramidal cells were recorded in voltage-clamp at −70 mV, with map recordings repeated 2–4 times.

For each neuron, a local input map was generated by thresholding at 6× standard deviation of the baseline, averaging traces at each stimulation site across trials, and then integrating responses within a 75-ms window after laser onset. The map was then plotted as a heatmap and overlaid with the reconstructed morphology of the recorded cell after accounting for its post-fixation shrinkage factor (Extended Data Fig. 9b,c). For stimulation sites that overlap with the dendritic morphology, currents elicited by direct activation of glutamatergic receptors on the recorded neurons were separated from synaptic responses by their fast onset (20% peak response time < 10 ms) and were excluded from connectivity analysis. Next, to account for cortical curvature and differences in cortical thickness along the anterior-posterior axis, individual heatmaps were interpolated along each stimulation column between pia and corpus callosum to produce the same number of response pixels per column (n = 22 pixels) (Extended Data Fig. 9g). To generate the average input maps, interpolated heatmaps were either normalized to the maximum response of individual maps (Extended Data Fig. 9e) or kept as absolute mean without normalization (Fig. 5b,d, and Extended Data Fig. 9d), and grouped by soma layer location (Fig. 5b and Extended Data Fig. 9d,e). To average, individual maps were aligned by pia such that all somas were distributed by their cortical depth on the vertical axis and shared the same relative anterior-posterior location on the horizontal axis (Fig. 5b and Extended Data Fig. 9e), or additionally were aligned by soma such that all somas were also positioned to the same location at the center depth of each layer group (Extended Data Fig. 9d). After alignment and averaging, all pixels with sample repeats (n ≥ 2) were displayed, and pixels at the horizontal edges with single samples (n = 1) were hidden. For visualization only, averaged heatmaps were Gaussian filtered, contoured at 18^th^ and 85^th^ percentiles of all responses, and overlaid on heatmap displays (Fig. 5b,d and Extended Data Fig. 9d,e). To generate the circuit diagram (Fig. 5c), absolute mean input strength was calculated by summing the absolute inputs arriving from each input layer and dividing by the number of recorded postsynaptic neurons in the receiving layer; only inputs with mean strength > 0.15 nA were shown. To generate the probability map of input occurrence (Extended Data Fig. 9f), interpolated heatmaps were binarized by assigning responsive or non-responsive pixels with values of 1 or 0, respectively. The binary maps were then aligned by pia, grouped by soma layer location, and averaged across the same stimulation sites. To aid visualization, the 20% occurrence rate was calculated from Gaussian-filtered counterparts of the averaged binary maps and overlaid on top of the unfiltered averaged binary maps. Interpolated heatmaps were collapsed vertically or horizontally to 1D by summing along columns or rows, respectively (Extended Data Fig. 9g–i). Individual collapsed maps were grouped by morphology type (Extended Data Fig. 9h,i) and averaged. To calculate normalized input strength arriving at each AI cortical layer (Extended Data Fig. 9j), within-column and lateral inputs from each layer were first normalized to the total cortical inputs of each cell, and then averaged across cells of the same soma layer.

### Feature extraction

#### Cell morphology feature extraction

L-measure^97^ was used to calculate the numbers of stems, bifurcations, branches, and tips; as well as dendritic width, height, length, Euclidean and path distance, branch order, contraction, partition asymmetry, and local and remote bifurcation amplification. All remaining morphological features (Supplementary Table 1) were extracted using custom-written software (BrainCellSuite). For features that described dendritic length within a specified cortical laminar bin—i.e., a volume defined by an upper and lower border (Y-axis, infinite along X- and Z-axes) parallel to the pial surface—we summed the 3D segment lengths that fell within each specified bin. In cases where a dendritic segment traversed multiple bins, the segment was split at its intersection with each border plane; resulting sub-segments were included in the calculations for their respective bins. To calculate the overlap between apical and basal dendrites, we first established for each morphology a 3D density matrix with a voxel size of 18 × 18 × 18 µm based on the dataset averaged soma diameter of 18.07 µm. We then registered the total apical and basal dendritic lengths separately per voxel. We calculated the number of voxels that contained both apical and basal dendrites as a fraction of all voxels that contained a dendrite, and the total dendritic length within voxels which contained both apical and basal dendrites as a fraction of the total dendritic length. For the 18-µm lamina features we first collapsed voxels along the Y- and Z-axes—i.e., parallel to the pial surface—into laminar cortical bins and then calculated the same fractions as stated above. The voxel-based apical-basal overlap features were log_2_-transformed to reduce their strong non-normal distribution, skewness (0.96 and 0.96, respectively), and kurtosis (1.26 and 1.07, respectively).

#### Electrical property feature extraction

Custom-written software (BrainCellSuite), which included the importing of Ephus data files (XSG), was used. Feature details are described in Supplementary Table 1. A measured liquid-junction potential of 14 mV was not corrected for. Spike detection was performed on 1 kHz low-pass Bessel-filtered signal traces in which putative spikes were identified using the rise rate criteria ≥ 20 mV/ms. Spike characteristics (features) of the identified putative spikes were then extracted from the raw signal traces. Spiking behavior features were extracted from traces with a depolarizing 1 s-long current step at rheobase plus 125 pA.

Summary plots of electrical properties were made for each recorded cell, which included access and input resistance, resting membrane potential and delta membrane potential between start and end of recorded sweeps, action potential absolute peak and relative to spike onset, and noise and patch RMS (root mean squared; 1.5 and 50 ms periods, respectively). Large deviations across sweeps were manually inspected and individual sweeps were omitted when deviations indicated technical noise or loss of whole-cell mode. Cells were included for iCCA analysis when access resistance < 40 MOhm, action potential threshold ≤ −20 mV and amplitude ≥ 40 mV, and single action potential and spike train data was available.

#### LSPS map feature extraction

The interpolated averaged input maps that were normalized to the maximal input were used. Pixel-wise (dis)similarity measurements, such as mean squared error or structural similarity index,^98^ were attempted as distance matrices for the iCCA and were found to be confounded by the exclusion of direct excitation from recorded neurons. Instead, we extracted input features that resembled input strength and number based on local layer and soma-centric proximity allocations. To do so, the assigned location of each soma on our insular atlas was used as a reference point for registering its input map to the insular atlas, thus allowing us to allocate each input location of the interpolated input map to a cortical layer. Next, we allocated the columns of the input map to either proximal or distal input sites based on two levels of a lateral input border: The dendritic density-based (d-b) lateral border was based on the 2.5 – 97.5 percentile range of the summed dendritic densities of all morphologies included in this dataset, spanning 342 µm (5 proximal columns, rounded to 5 columns × 75 µm per column = 375 µm) laterally on each side of the soma. The dendritic reach-based (r-b) lateral border was based on the 2.5 – 97.5 percentile range of the summed most extreme dendritic reach of all morphologies in this dataset, spanning 675 µm (9 columns) laterally on each side of the soma. Using these two lateralization levels and layer allocations, we calculated relative input strengths and numbers of input locations (Supplementary Table 1). To calculate columnar input probability (Extended Data Fig. 10e) we converted the columnar collapsed input maps to a binary mask (i.e., columns with at least one input location yielded one). The input probability was calculated as the fraction of cells with an input location in the given column over the total sampled cells.

### Iterative consensus cluster analysis (iCCA)

iCCA was performed in two phases: First, on principal component analysis transformed datasets; second, on sets of biologically interpretable features (Extended Data Fig. 5a and Extended Data Table 1).

#### PCA feature-based consensus cluster analysis

During the first phase, we determined the number of putative clusters using PCA transformed data as the input data for consensus clustering. We established, per dataset modality (morphology, electrical properties, input source, or ‘cross-modal’ combinations thereof), a thousand iterations of 90% subsampled datasets with each PCA transformation. For each subsampled dataset, we determined the optimal number of principal components using column-wise K-fold cross-validation (MEDA toolbox).^99^ In short, a PCA-model containing a variable number of principal components was fit to each subsampled dataset and used to reconstruct the original subsampled dataset based on a single column (i.e., feature). An estimation error was calculated by comparing the reconstructed and original datasets (see Camacho et al.^99^). This process was repeated for all features and the resulting summed estimation error was used to identify the ‘optimal’ number of principal components for the subsampled dataset, defined by the lowest summed estimation error (Extended Data Table 1). Hierarchical cluster analysis (HCA, Euclidean distance and Ward linkage) was performed on each PCA-transformed subsampled dataset for which cluster allocations were determined using the CuTreeDynamics Python package.^100^ The resulting dendrogram allocated leaves (i.e., individual cells) to the clusters based on a set of criteria including minimal number of cluster members (n = 3 cells), intra-cluster cohesion, and inter-cluster distance, which were collectively controlled by one hyperparameter: ‘deepsplit’. To evaluate the cluster results, a pair-wise co-cluster score was calculated by taking the number of HCA iterations during which a cell pair was allocated to the same cluster (irrespective of the cluster identity across HCA iterations) relative to the total HCA iterations during which the cell pair was included (±900 iterations). The co-cluster score matrix therefore resembled a pair-wise similarity score for which a score of 1 indicated that the cell pair was always allocated to the same cluster. The inverted co-cluster matrix (1 – co-cluster matrix) was used as the distance input matrix for a second HCA (Ward linkage) to allocate all frequently co-clustered cell-pairs to the same cluster (i.e., consensus clustering). Based on this consensus clustering, we identified the most informative features that resembled distinct characteristics between pairs of clusters. To do so, we performed cluster pair-wise feature comparisons using the Mann-Whitney U test with a post hoc Bonferroni correction for multiple testing and selected all features (referred to as informative features, Extended Data Table 1) for which at least one cluster pair displayed significantly distinct feature distributions with the minimal fold-change criteria (morphological ≥ log_2_(1.75); electrical, input, or cross-modal ≥ log_2_(1.25)).

#### Biologically interpretable feature-based consensus cluster analysis

After generating a list of informative features from the above analysis, we performed the second phase of iCCA (Extended Data Fig. 5a) to identify clusters (cell types) based on biologically interpretable features. First, to reduce the number of highly-correlated and concentrated features (i.e., low coefficient of variation), we performed a grid search with unique feature sets using all combinations of criteria mentioned above for the Spearman correlation (≥ 0.70 – 1 with 0.05 increments) and the coefficient of variation (cv, ≤ 0.1 – 0.55 with 0.05 increments). With each unique feature set (Extended Data Table 1), we repeated the above-described consensus cluster analysis. In this part, however, instead of PCA-transformed data, the thousand iterations of 90% subsampled datasets were composed of z-score-transformed unique feature sets. The CuTreeHybrid ‘deepsplit’ hyperparameter level for the HCA on the individual subsampled datasets was kept the same as chosen for the PCA-transformed data; for the second ‘consensus’ HCA all hyperparameter levels were explored. To identify the feature set with the optimal consensus cluster result (based on the co-cluster scores), we compared the cluster results of all unique feature sets using the silhouette score while considering multiple ‘deepsplit’ hyperparameter levels (1–4), which produced increasing numbers of clusters. The feature set and ‘deepsplit’ parameter that yielded the highest silhouette score with a minimum number of clusters was deemed the optimal feature set and cluster allocation hyperparameter (Extended Data Fig. 5c and Extended Data Table 1). In addition to optimal cluster allocations determined by the highest silhouette score (e.g., k = 21 for M-types), we also showed a dendrogram cut level at k = 5 determined empirically for all iCCA analyses (Figs. 2a, and 4a and Extended Data Figs. 6a, 8a, 10a, 14a, and 15a) to display reduced clusters at a higher-level hierarchy and to reveal groupings of clusters allocated by the highest silhouette score.

To display the cluster results in a reduced dimensionality, we performed t-distributed stochastic neighbor embedding (t-SNE^101^; sklearn Python package). The z-score-transformed datasets of the selected feature sets were used as input data and t-SNE model parameters were either held constant (learning rate: 500; iterations: 5000; random initialization seed) or tuned (perplexity: 8–40; early-exaggeration: 1–10) to obtain the best possible visualization with low intra-cluster and high inter-cluster distances among cells.

### Simulation

Two approaches were taken to study the relationship between dataset size and number of identified clusters (Fig. 2b). First, a subsampling-based approach was used without corrections for data point imbalance, e.g., the frequency of cells located in the different cortical layers are different. We then assessed at which dataset size we could have stopped collecting data and obtained the same number of clusters, keeping all cluster analysis parameters fixed. Next, we assessed the upper bound of identifiable clusters and its relation to dataset size based on the data distribution. As mentioned previously, the sampling frequency of cells across cortical layers was not homogenous, e.g., deep L6 cells were less abundant in our dataset compared to L5 cells. To account for such imbalances in our simulation experiment, our second approach started by creating additional ‘fictive’ cells among these minority groups—located at supragranular and infragranular cortical depths—by imputing morphological feature data points using the synthetic minority over-sampling technique (SMOTE; imblearn Python package).^102^ The SMOTE-generated dataset was then fit with multiple Gaussian mixture-models (GMM, sklearn Python package; built with component-wise general covariance matrices; maximal number of expectation-maximalization iterations: 1000; number of initializations: 30) composed of varying numbers of components. Using the Bayesian information criterion (BIC), we selected the GMM with the fewest number of components and the lowest BIC score. The simulated datasets of varying sample sizes containing feature-based ‘fictive’ cells were generated using the selected GMM (6 components). Both the subsampled and simulated datasets were used for consensus clustering as described above. Like the original data analysis, the same feature set (Extended Data Fig. 5e) and CuTreeHybrid ‘deepsplit’ hyperparameter (level 2) during the HCA with 90% subsampled data was used (Fig. 2a). The consensus cluster analysis for each tested dataset size (subsampled datasets: 10, 20, 50, 100, 150, 200, 300, 400, 500, 600, 700, 800, or 900 cells; simulated datasets: 10, 20, 50, 100, 150, 200, 400, 600, 800, 1000, 1200, 1400, or 1600 cells) was performed for 20 independently initialized instances. The recorded cluster number (Fig. 2b) was based on either a fixed ‘deepsplit’ hyperparameter (level 2; subsampled datasets) or the maximal possible cluster number based on all four assessed ‘deepsplit’ levels (simulation datasets). The cluster number vs. dataset size results were modeled with a five-parameter logistic regression that takes the form:

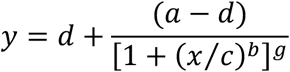

Model parameters were obtained using non-linear least squares to fit to our data (simulated datasets: *a* = - 1.967, *b* = 0.9569, *c* = 1.964×10^10^, *d* = 21.91, *g* = 1.105×10^8^; subsampled datasets: *a* = 0.8220, *b* = 1.967, *c* = 18.45, *d* = 30.12, *g* = 0.1750).

### Classifier training

Random forest classifiers (sklearn Python package) were trained on 75% of the dataset using untransformed features from morphological, electrical, or input modalities, or cross-modal combinations thereof (Fig. 6h). Classifier performances were assessed (F1-score, the harmonic mean of precision and recall) using the remaining unseen 25% of the dataset. For performance quantification we trained and assessed random forest classifiers over five independently initialized instances. For the ranking of feature contribution displayed in z-score-transformed heatmaps (Extended Data Figs. 5e, 8b, 10b, 14b, and 15b), we used the Gini-index based on random forest classifiers trained on the full dataset as the readout. We confirmed that the out-of-bag (OOB) performances of the full dataset-trained classifiers were similar to that of the classifiers trained on 75% of the dataset (OOB_full_ – OOB_75%_ ≤ 7%). The Gini-index describes the total normalized reduction of the Gini impurity by a given feature, i.e., a high Gini-index indicates greater feature contribution to the classifier performance.

### Topographic analysis

To estimate the topographic split line based on spatial distribution of M-types (Fig. 3a–c), we first determined individual optimal segregation lines between each M-type pair located in the same layer (based on their predominant layer assignment; Fig. 2c) along the long axis of the insula. These segregation lines were defined as the location at which two M-types, in a pair-wise comparison, were best distinguished based the distribution of their somas along the long axis. To establish the segregation line for a pair of M-types, we first used the relative positions of median soma location to designate each M-type to either the dorsoanterior or ventroposterior side of the long axis. Cumulative probability plots of soma locations were generated for both M-types: for the dorsoanterior-biased M-types (e.g., Fig. 3a: *L5b hourglass*), F(x) = 1 was located at the dorsoanterior-most position and decreased towards F(x) = 0 at the ventroposterior-most position, whereas ventroposterior-biased M-types were oriented in the opposite direction (e.g., Fig. 3a: *L5b wide*). The location along the long axis with the lowest sum of both cumulative probabilities was defined as the segregation line between this pair. A topographic split for the entire dataset was calculated using the average of all segregation lines that met the following criteria: (1) the distributions of soma locations between the two M-types were significantly shifted along the long axis (Mann-Whitney U test with post hoc false-discovery rate Benjamini-Hochberg correction), and (2) the segregation line was located between the median soma locations of the two M-types. The topographic split line was located very anteriorly relative to the DI and GI—only two mapped cells in the DI were located anterior to this split line. Therefore, only cells mapped to the AI in our dataset were allocated to either the dorsoanterior-AI (dorsoanterior-AI) or ventroposterior-AI (ventroposterior-AI), based on their relative position to the topographic split line along the long axis (Fig. 3d). Statistics on topographic distributions of these dorsoanterior-AI and ventroposterior-AI cells were performed. To test whether M-types had a predominant topographic localization to either dorsoanterior-AI or ventroposterior-AI we compared the frequency of both allocations (dorsoanterior-AI/ventroposterior-AI) within each M-type to the frequencies of all other M-types residing in the same 95^th^ percentile range of the cortical depth using the Fisher exact test with post hoc false-discovery rate Benjamini-Hochberg correction. The local control population consisted of all AI cells, excluding the cells allocated to the tested M-type, located within the same cortical depth range (2.5 – 97.5 percentile range) as the tested M-type.

The within-layer and -region (L2 dorsoanterior-AI vs. L2 ventroposterior-AI) finding probabilities (Fig. 3d) were calculated by dividing the frequency of specified M-type cells allocated to a given layer-region combination (e.g., L2 dorsoanterior-AI) by the total number of cells allocated to that layer-region.

### Cross-modal feature and cell type intersectional analysis

To identify highly correlated cross-modal features we performed Spearman correlation analysis on a curated list of features. This list included all features identified during the iCCA for morphological, electrical, and input cell type analyses (Extended Data Figs. 5e, 8b, and 10b), and included number of apical stems, resting membrane potential, adaptation index, and mapped locations in the insular flatmap. Note that the untransformed absolute versions of morphological and input features were used for Spearman correlation analysis, as opposed to the relative versions used during iCCA. Correlation analysis was only performed on combinations of features belonging to two different modalities, p-value was based on permutation test with post hoc false-discovery rate Benjamini-Hochberg correction (n = 1911 feature combinations, Extended Data Fig. 12).

To identify characteristic features of cell types we performed statistical feature distribution comparisons across a list of curated pair-wise cell type combinations (n = 309 combinations). The cell type combination list was constructed based on morphological, electrical, local connectivity (input), and topographic classifications (n = 102 unique cell types, with n ≥ 3 cells per cell type). The same feature list, excluding mapped locations in the insular flatmap, was used to perform the feature distribution comparisons (Mann-Whitney U test with post hoc false-discovery rate Benjamini-Hochberg correction, n = 6900 tested comparisons). Only features available for the specified cell type combination (n ≥ 3 cells per cell type for a given feature) were tested and displayed. For each feature comparison the log_2_(fold-change) was color-coded and the bubble size was displayed based on p-value thresholds, corrected for multiple testing (Supplementary Fig. 3).

Cross-modal cell type classifications were displayed using a parallel categories plot (plotly Python package). Vertical locations of categories were manually adjusted for display.

### Channelrhodopsin-assisted circuit mapping (CRACM)

Using brain slices from mice injected with virus expressing ChR2 in the VPMpc/PF thalamus, anterior AI neurons were recorded in voltage clamp at −70 mV, while ChR2-expressing thalamic axons in the AI were stimulated by a 473 nm laser (CL473-025, Crystal Laser, Reno, NV, USA) in two 50 × 50 μm-spaced stimulation grids (Fig. 7a,b). One stimulation grid was positioned in the posterior AI covering the dense expression zone of VPMpc/PF axons. The other stimulation grid was positioned in the anterior AI covering the dendrites and soma of the recorded neuron. Stimulation trials (complete grid) were repeated 3–5 times and corresponding responses were averaged across the same stimulation sites. To quantify the strength of multisynaptic integration (posteriorly positioned grid) vs. monosynaptic response (anteriorly positioned grid), we calculated the posterior response index (Fig. 7e), where Σ responses to posterior stimulation is the sum of all averaged responses to stimulations in the posterior AI, and Σ responses to anterior stimulation is the sum of all averaged responses to stimulations in the anterior AI:

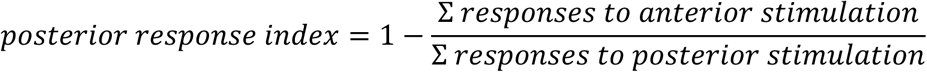

### Statistics

No statistical methods were used to predetermine sample size; the group sizes used in each experiment are comparable to those generally employed in the field. Multiple cells were obtained per mouse (median 3 cells per mouse, range 1 – 13, Supplementary Table 3). Data collection and analysis was not performed blind to the conditions of the experiments. Criteria for excluding observations from analysis are listed in appropriate method sections. Unless otherwise stated, mean ± s.e.m. was used to report statistics. The statistical tests used, definition of n, and multiple comparisons correction where appropriate are described in the figure legends and methods. All statistical tests were two-sided with significance defined as alpha = 0.05. Statistical analyses were performed in R, Matlab, or Python.

## Acknowledgements

We thank Tess Lameyer for excellent technical support in the early phase of the project, and Drs. Kylie McPherson and Jian Qiu for contributing, respectively, 16 and 7 recorded cells during their research stay at the Mao Laboratory. We thank Dr. Stefanie Kaech Petrie and the Advanced Light Microscopy Core facility at Oregon Health & Science University for Axio Zoom imaging. We thank Dr. Haining Zhong and all members of Mao and Zhong Labs for discussions throughout this project. We thank Drs. Corette Wierenga, Haining Zhong, and Gary Westbrook for critical feedback on the manuscript. This work was supported by the Netherlands Organisation for Scientific Research (NWO Veni Research Grant: VI.Veni.192.231 to B.C.J.), a Lacroute Fellowship (Y.C.), BRAIN Initiative awards (National Institute of Health, United States) RF1MH120119 (T.M.), R01NS104944 (T.M.), and RF1NS133599 (T.M.), as well as an R01 grant R01NS081071 (T.M.).

## Extended Data

**Extended Data Fig. 1:**
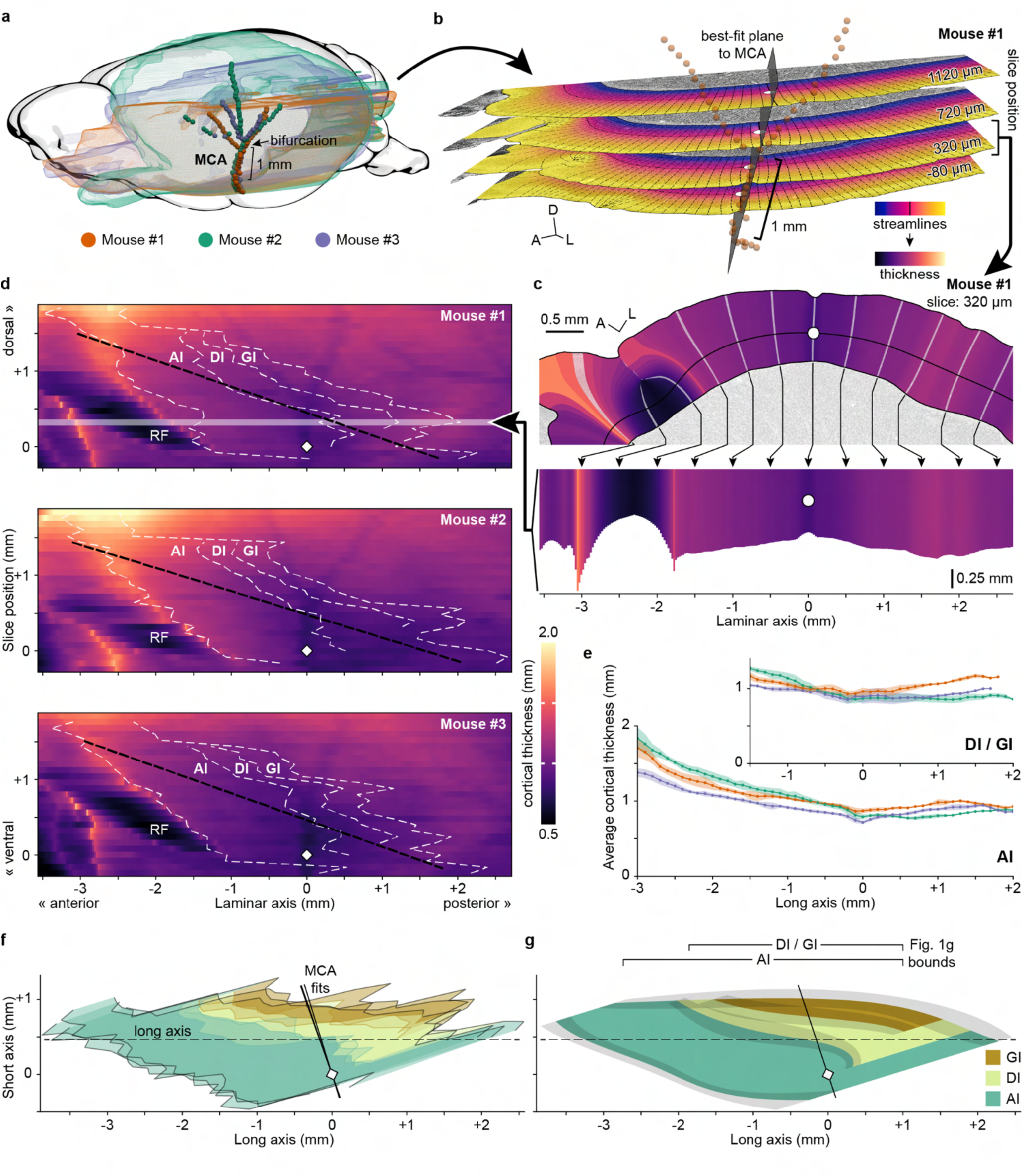
Construction of an average insular 2D flatmap. **a**, 3D reconstructions from brain sections in Supplementary Fig. 1 are registered to a single coordinate space. The excellent spatial alignment of the proximal MCA across three animals (Mouse #1, #2, and #3; colored spheres), along with its proximity to the insular cortex, motivated its utility as an anchoring point for organizing spatial data. Model brain adapted from Janke et al.^103^ was used for illustration only. Visualization was mirrored to left hemisphere for consistent display orientation. **b**, Four example sections from Mouse #1 (Supplementary Fig. 1) shown in 3D space with Laplacian streamline overlays (Fig. 1d). White circles: the points of intersection between the Laplacian isocontour lines and the vertical 2D plane (gray) fit to the proximal MCA annotations (orange spheres; Methods). **c**, Estimated cortical thickness across the laminar axis on an example section from Mouse #1. Arrows (displayed at 0.5 mm intervals) show the relative mapping of 25-µm wide streamline bins (white lines) from the physical section (top) to a ‘flattened’ laminar axis plot (bottom). White circles: MCA-derived coordinate origin. **d**, Individual flatmaps of the right insular cortex for the three animals from (**a**). Each flatmap is a grid of 25-µm horizontal × 80-µm vertical voxels. The 2D coordinate origin (white diamonds) corresponds to the MCA position on the ventral-most Nissl section containing layer 4 (Supplementary Fig. 1). Vertical axis (i.e., ‘cutting axis’) represents tissue sections; horizontal axis represents laminar distance from MCA origin (Fig. 1d). Luminance indicates cortical thickness (i.e., streamline length in (**c**)). The relatively dark zone at the lower-left of each map reflects the curvature of the rhinal fissure (RF). White dashed lines: annotated boundaries of insular subregions; black dashed line: long axis of an ellipse fit to the outermost boundaries of the insular annotations; thick arrowhead and semitransparent bar: row in Mouse #1 corresponding to data shown in (**c**). **e**, Mean cortical thickness in the AI (bottom) or DI/GI (top) for each mouse, binned along the long axis. Shaded regions: standard deviation. **f**, Overlay of insular flatmap subregion boundaries from all three mice. Data are aligned by rotating each flatmap about its MCA coordinate origin (white diamond) such that the long axis (dashed line) is horizontal. Solid black lines: trajectories of the best-fit planes to each animal’s MCA. **g**, An average 2D flatmap of the insula was generated by spatially averaging and smoothing (cubic spline; Methods) each set of lines from (**f**). Gray shading: smoothed standard deviation. A, anterior; D, dorsal; L, lateral; AI, agranular insula; DI, dysgranular insula; GI, granular insula; MCA, middle cerebral artery; RF, rhinal fissure.

**Extended Data Fig. 2:**
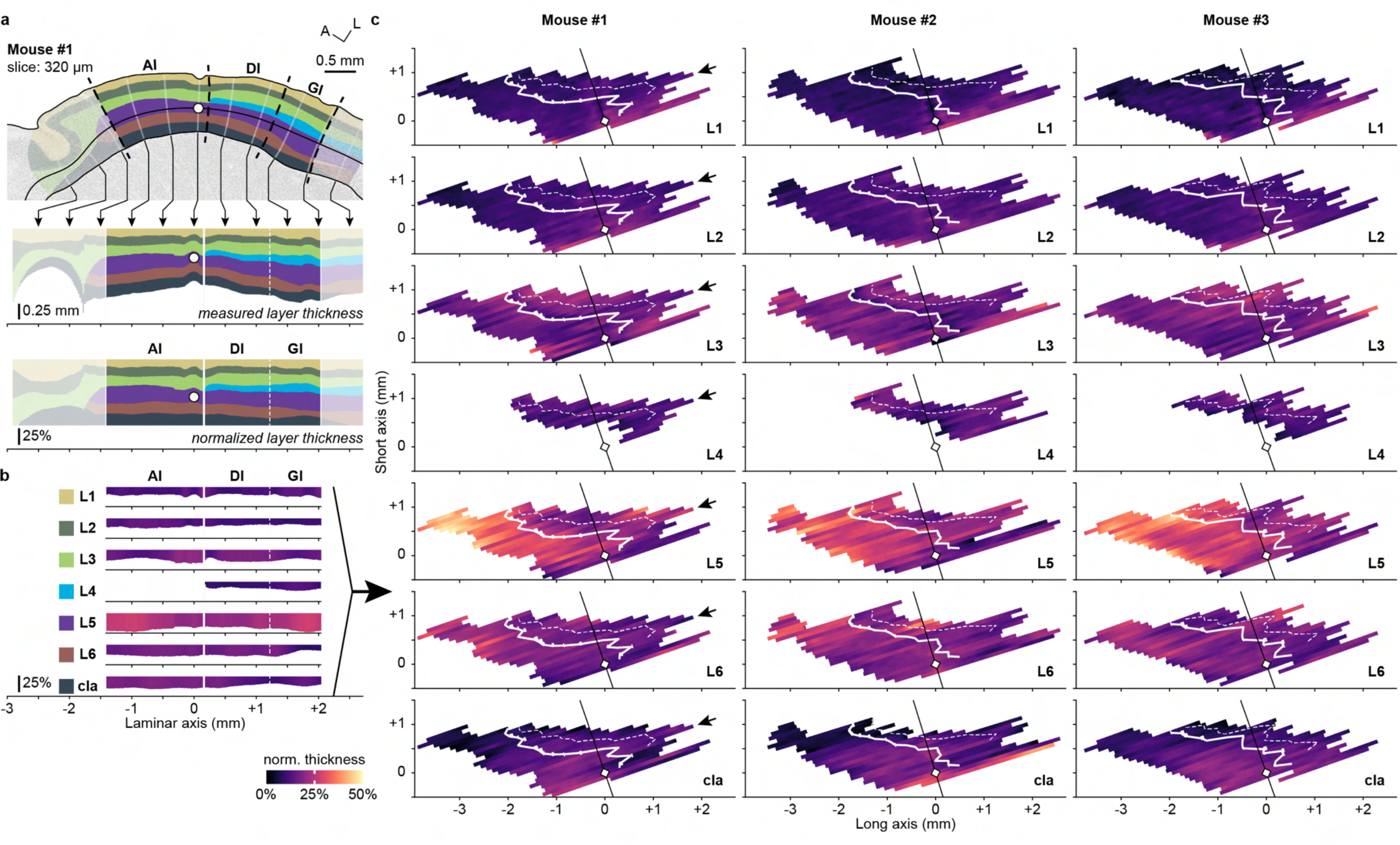
Quantification of insular cortex layer thickness. **a**, Example process of defining layer boundaries and calculating relative cortical depth. The layers were first manually annotated (top; Fig. 1c and Supplementary Fig. 1). The section was then ‘flattened’ according to streamlines (middle; Fig. 1d) and normalized by cortical thickness (bottom; Extended Data Fig. 1c and 1e). Arrows (displayed at 0.5 mm intervals) show the relative mapping of 25-µm wide streamline bins from the physical section (top) to a ‘flattened’ laminar axis plot (middle/bottom). White circles: MCA-derived coordinate origin. **b**, Visual representations of the change in normalized thickness for each layer from (**a**) along the laminar axis. Luminance indicates normalized cortical thickness. **c**, 2D flatmaps of relative layer thickness for each reference mouse. White diamond: MCA origin; black line: best-fit plane to MCA; thick white line: AI/DI border; thin dashed white line: DI/GI border; arrowheads: row in Mouse #1 corresponding to data shown in (**a** and **b**). A, anterior; L, lateral; AI, agranular insula; DI, dysgranular insula; GI, granular insula; MCA, middle cerebral artery; cla, claustrum.

**Extended Data Fig. 3:**
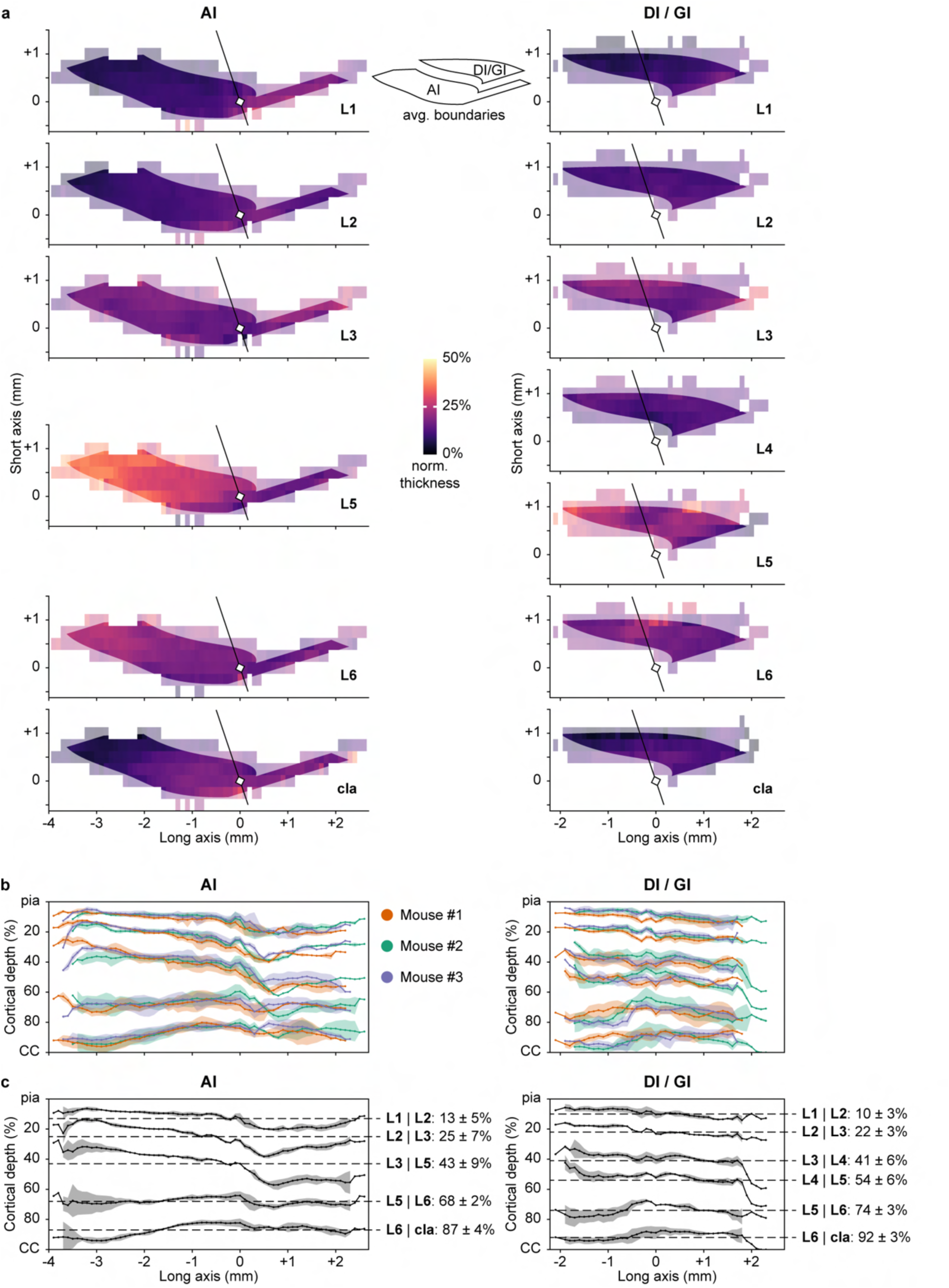
Mean layer thickness across the insular cortex. **a**, 2D flatmaps showing mean normalized layer thickness for AI (left) and DI/GI (right), respectively. Values from individual flatmaps (Extended Data Fig. 2) were binned at 100-µm and 250-µm intervals along the long and short axes, respectively, relative to the MCA origin (white diamond). Luminance indicates normalized cortical thickness. Shaded areas: the average boundary of each subregion (Extended Data Fig. 1g); black line: best-fit plane to MCA. **b**, Mean normalized position of layer boundaries (Extended Data Fig. 2) binned along the flatmap long axis separately for AI (left) and DI/GI (right) for each mouse. Shaded regions: standard deviation. **c**, Grand mean of (**b**) for AI (left) and DI/GI (right). Dashed lines: subregion-averaged cortical depth values (shown at right), obtained by subsequent averaging across the long axis. AI, agranular insula; DI, dysgranular insula; GI, granular insula; MCA, middle cerebral artery; cla, claustrum; CC, corpus callosum.

**Extended Data Fig. 4:**
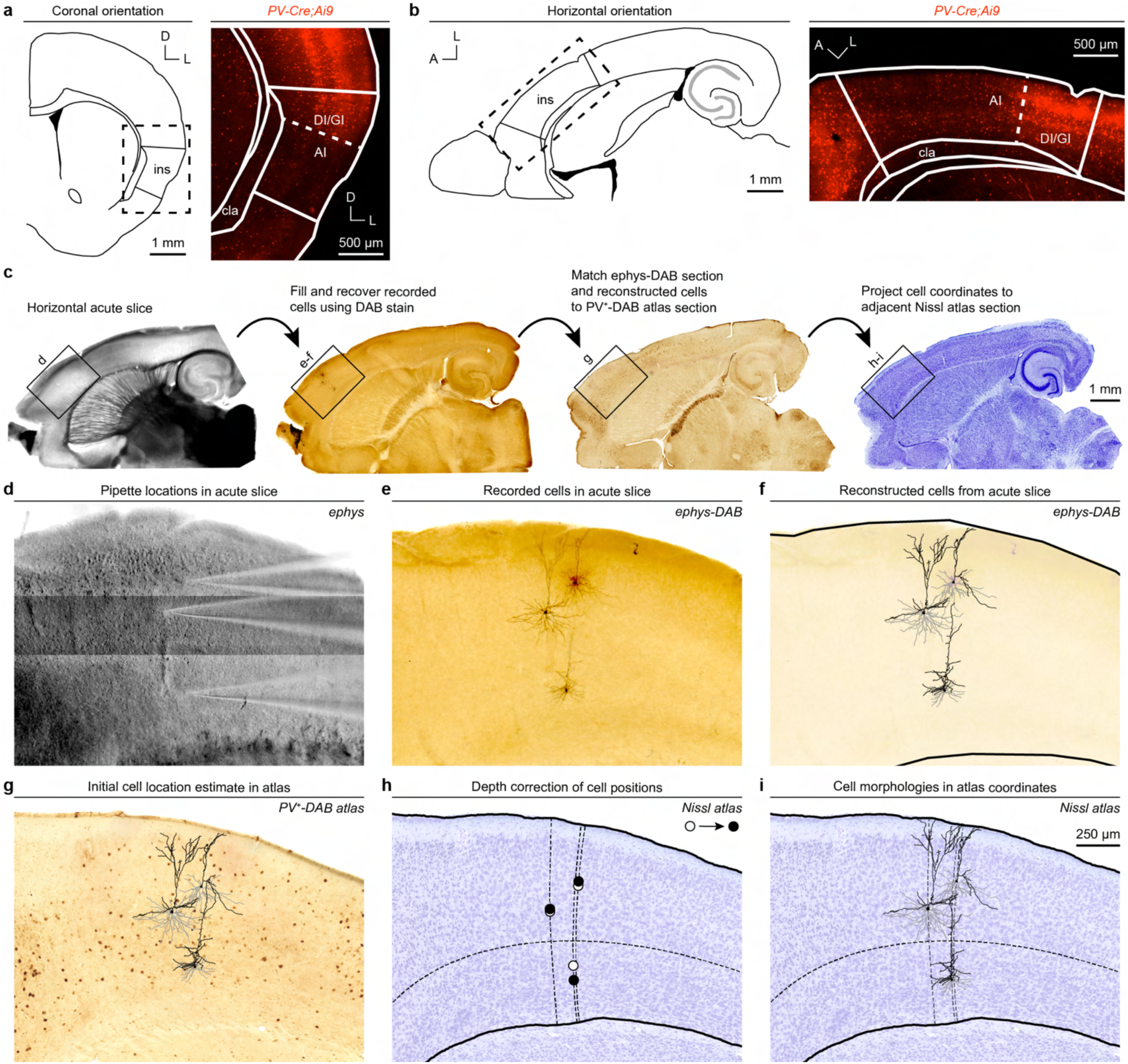
Registration of recorded cells from acute slices to the Nissl atlas. **a** and **b**, Example coronal (**a**) and horizontal (**b**) sections from *PV-Cre^+/-^;Ai9^+/-^* mice highlighting the paucity of PV^+^ somas in the AI. Boundary annotations adapted from Franklin and Paxinos.^45^ Dotted boxes on schematics: image locations. **c**, Overview of registration procedure. **d**, Photomontage of pipette locations for three separate electrophysiology recording sessions from the same acute slice. **e**, Recovery of dendritic morphologies from (**d**) following DAB staining. **f**, Reconstructions of cells from (**e**). Solid lines: pia (top) and CC (bottom) annotations. **g**, Estimated positions of cells from (**e** and **f**) on matched PV^+^-DAB atlas section following alignment. **h**, Estimated positions of cells from (**e** and **f**) on Nissl atlas section following ventral projection from adjacent PV^+^-DAB atlas section, both before (open circles) and after (filled circles) correction of cortical depth (Methods). **i**, Cell morphologies in final atlas coordinates. Vertical dashed lines: streamlines intersecting cell positions; horizontal dashed line: mid-value Laplacian isocontour.

**Extended Data Fig. 5:**
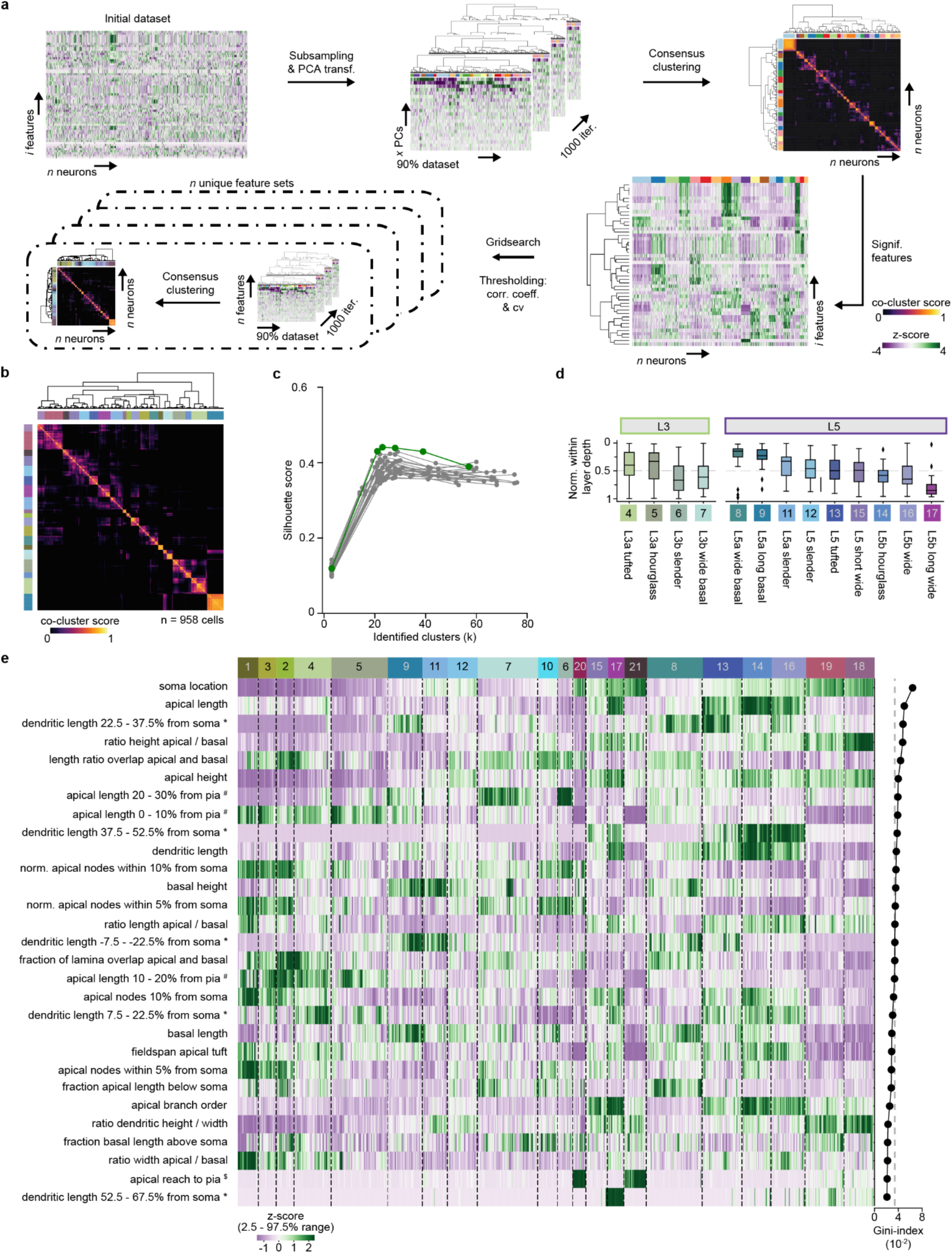
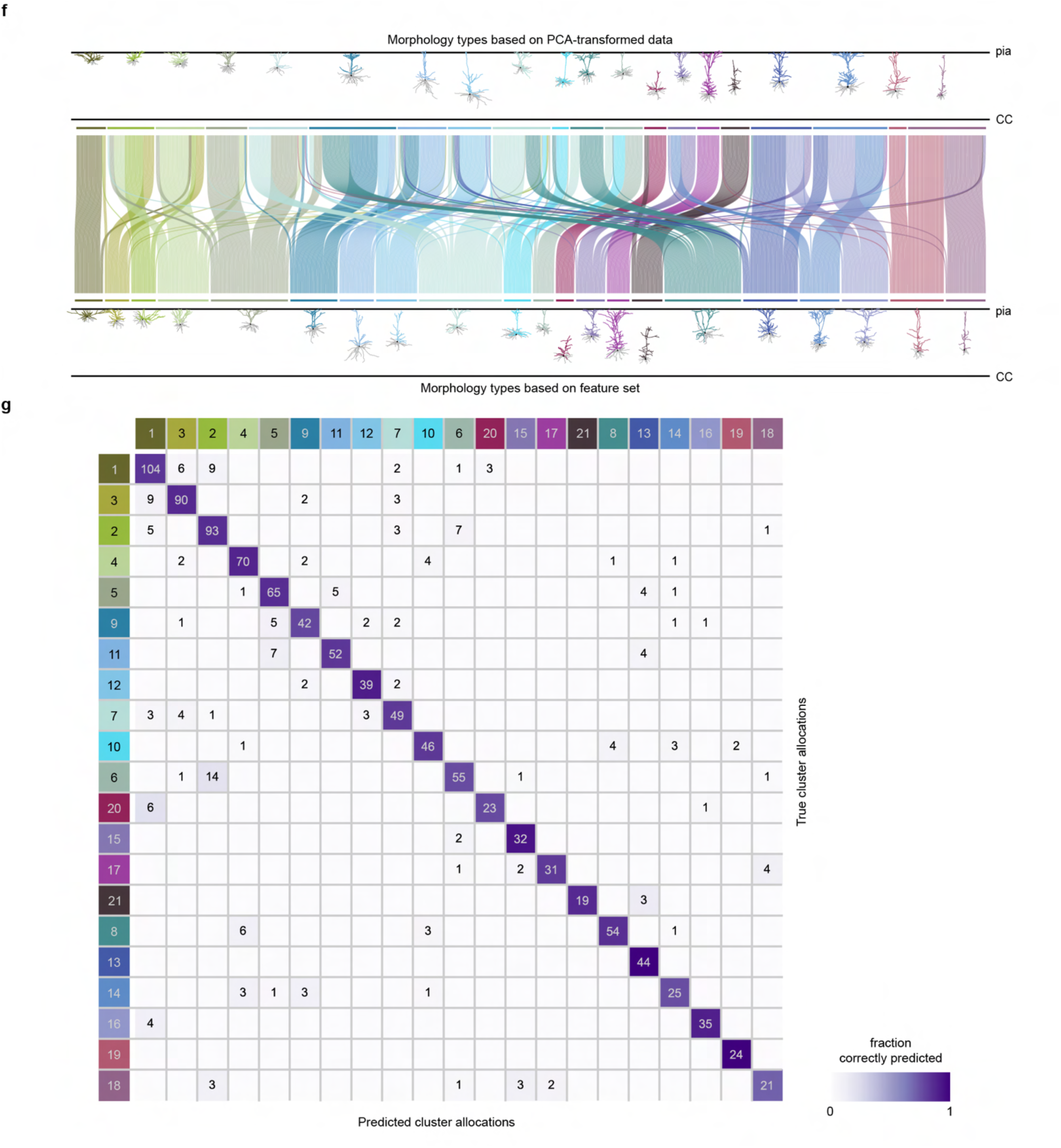
Iterative consensus cluster analysis pipeline and evaluation of its performance. **a**, Analysis pipeline of iterative consensus clustering analysis (iCCA) using neuronal properties (i.e., features; Methods). **b**, Hierarchical cluster analysis (HCA) on pair-wise co-cluster scores across 958 cells using PCA-transformed morphology dataset initially identified 20 morphology clusters (different from the final 21 M-types), color-coded. **c**, Feature set performance as described by the silhouette score and number of identified clusters (k) for each tested unique feature set (individual curves) at different dendrogram cut levels (3 clusters and 4 ‘deepsplit’ levels; cutreeHybrid function from Python and R dynamicTreeCut packages). The best-performing feature set is highlighted (green). **d**, The within-layer locations of several morphology types are biased to either the upper or lower half of the layer, thereby allocating them into L3a vs. L3b, or L5a vs. L5b morphology types. **e**, Z-scored morphology features (left), ranked by random forest classifier Gini-index (right, gray dashed line indicates equal feature contribution [1/29 features]). * Dendritic length located within a 15% local cortical thickness bin positioned relative to the soma, normalized to local cortical thickness. ^#^ Apical length within a 10% local cortical thickness bin positioned relative to the pia, normalized to total apical length. ^$^ Vertical distance between pia and closest apical dendrite, normalized to local cortical thickness. **f**, Representative morphologies for each cluster based on the PCA-transformed dataset (top row) or the best-performing iCCA unique feature set (bottom row) and cell-wise correspondence between the two analysis approaches (adjusted Rand index = 0.36). **g**, Confusion matrix for the performance of the random forest classifiers, displayed as the mean percentage of correctly-predicted cells per M-type relative to the corresponding true cluster size from iCCA. Purple luminance: percentage of neurons predicted by classifiers; numbers in colored boxes (left-most column and top row): M-type cluster number; numbers in purple boxes: n of cells (when n > 0). For (**d**), (**e**), and (**g**), colored boxes with numbers: M-types color-coded for k = 21 in Fig. 2a.

**Extended Data Fig. 6:**
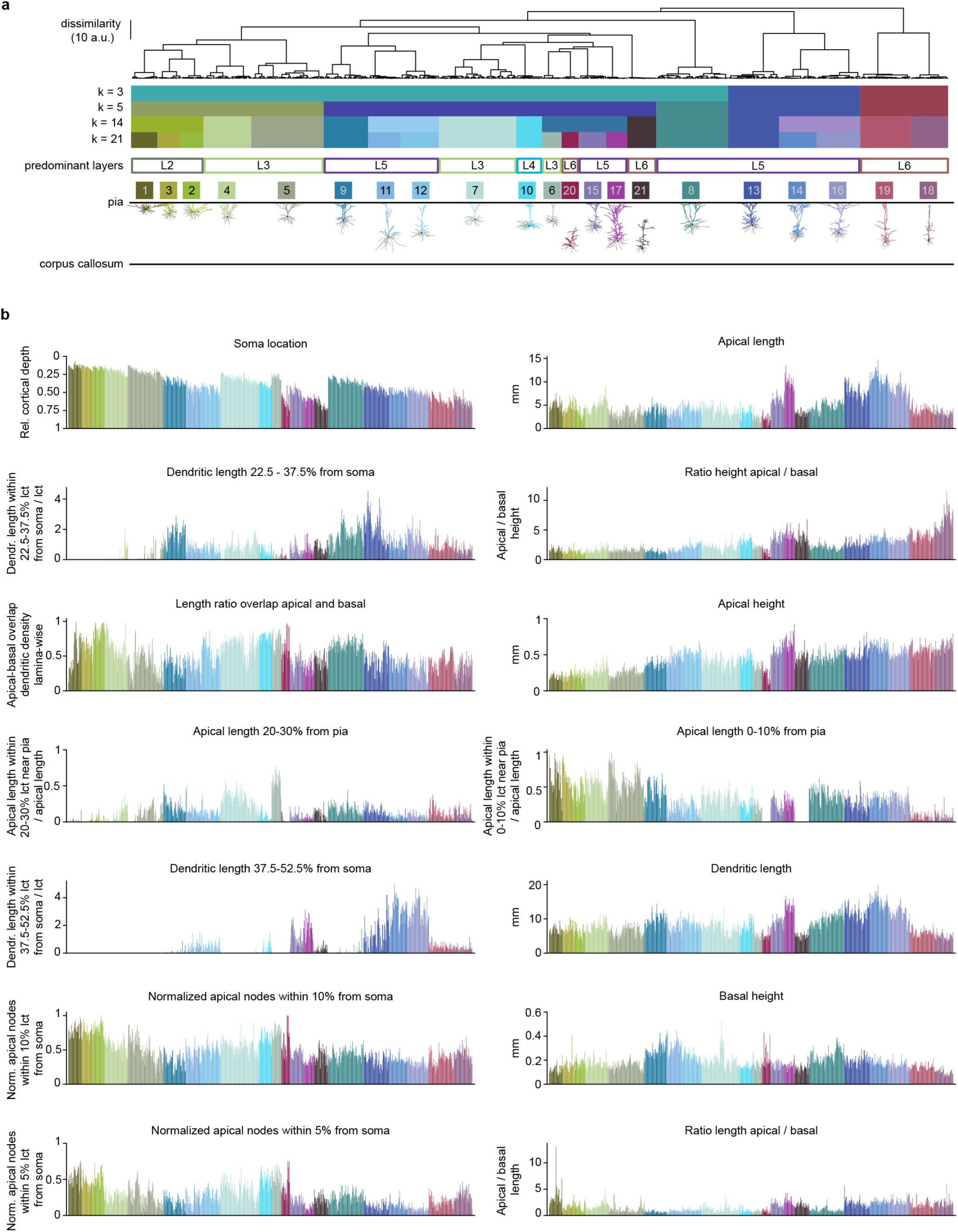
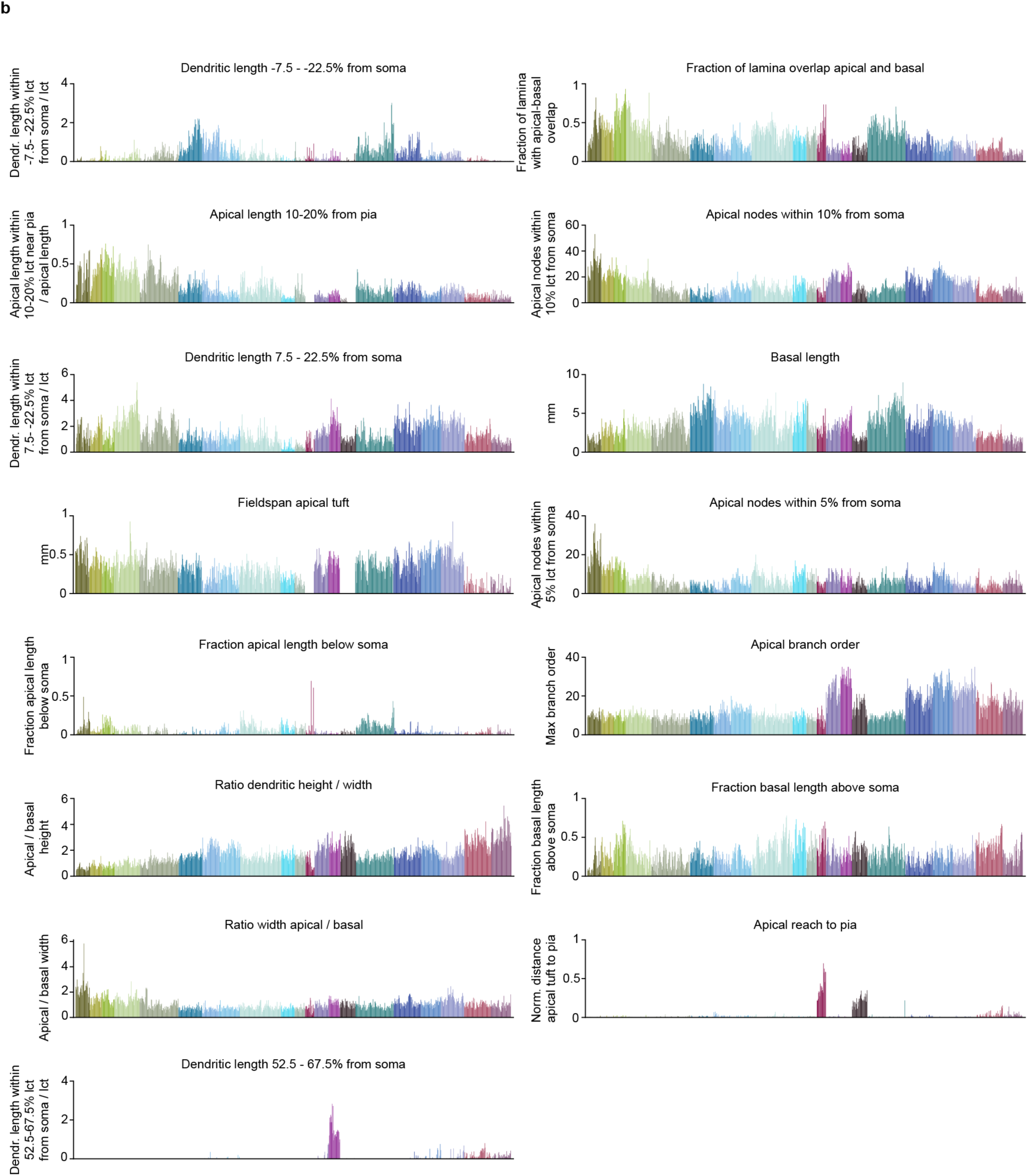
Distinct features of M-types. **a**, Consensus cluster dendrogram cut at different levels (k = 3, 5, 14, 21 clusters) indicated by color coding below the dendrogram to show how M-types relate hierarchically. The representative morphologies (bottom) were aligned to k = 21 clusters, same as Fig. 2e. Layer allocation was based on the predominant layer. **b**, All 29 features used for the iCCA, sorted by random forest classifier-derived Gini-index (Extended Data Fig. 5e). Histogram bars sorted in the same sequence as the dendrogram in (**a**), color-coded based on corresponding M-types.

**Extended Data Fig. 7:**
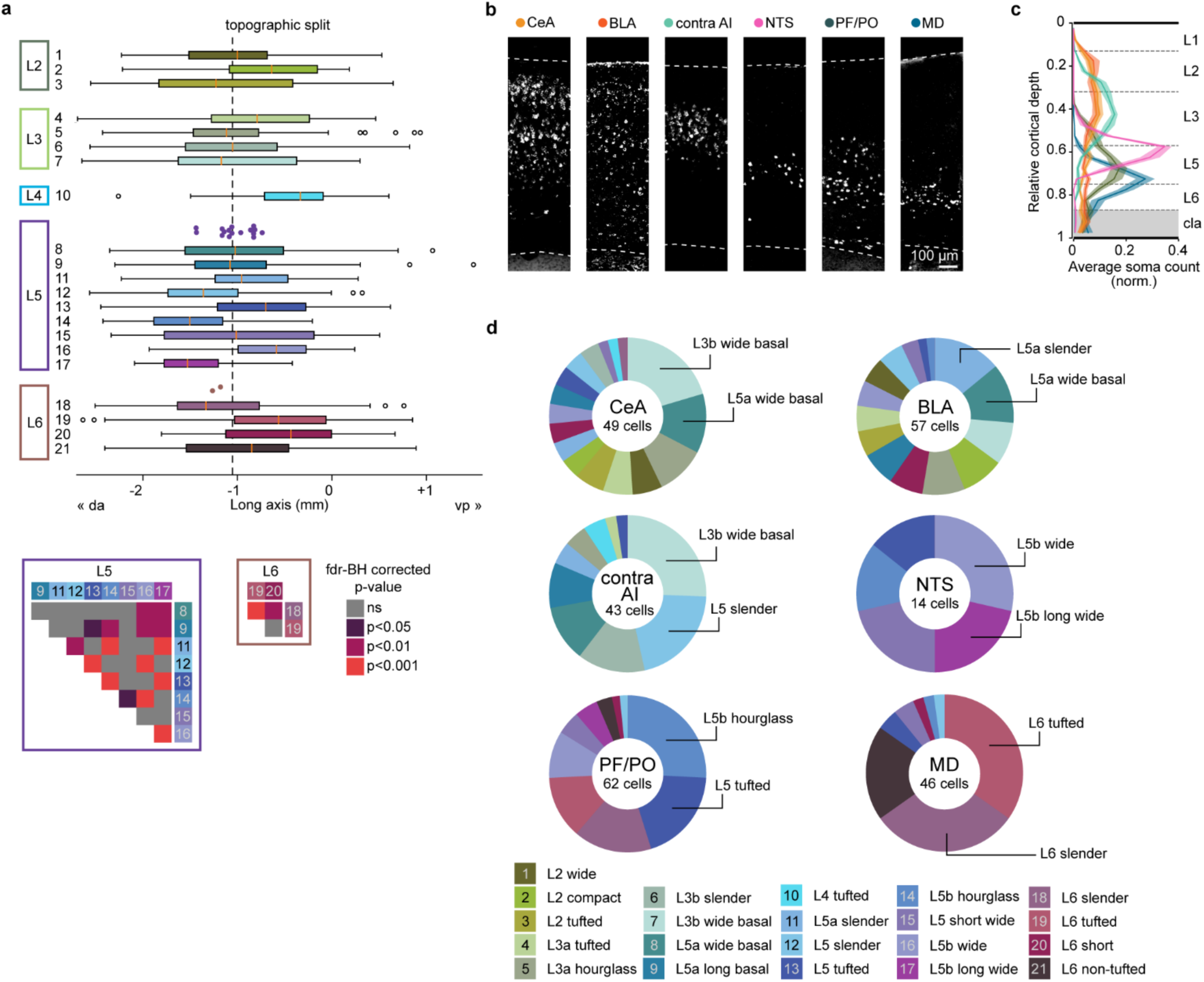
Topographic- and projection-based distributions of M-types. **a**, Topographic distribution of all M-types along the long axis of the insula (top), and the corresponding heatmaps of p-values from within-layer pair-wise comparisons in L5 and L6 (bottom, Mann-Whitney U test with false-discovery rate Benjamini-Hochberg (fdr-BH) correction). Note that all pair-wise comparisons within L2 and L3 were non-significant and, therefore, not shown here. Color-filled circles: segregation points from all significant within-layer comparisons; open circles: outliers; dashed line: average position of segregation points; orange ticks: median location for individual M-types. **b**, Representative photomicrographs of retrogradely-labeled neurons within the insular cortex that project to the CeA, BLA, contra AI, NTS, PF/PO, and MD, respectively. **c**, Quantification of retrogradely-labeled soma counts across the cortical depth, color-coded corresponding to (**b**). Shaded region: standard error. **d**, M-type composition of each projection target. The top two most-abundant morphology types are annotated. CeA: central amygdala; BLA: basolateral amygdala; contra AI: contralateral agranular insula; NTS: nucleus tractus solitarius; PF/PO: parafascicular/posterior thalamus; MD: mediodorsal thalamus.

**Extended Data Fig. 8:**
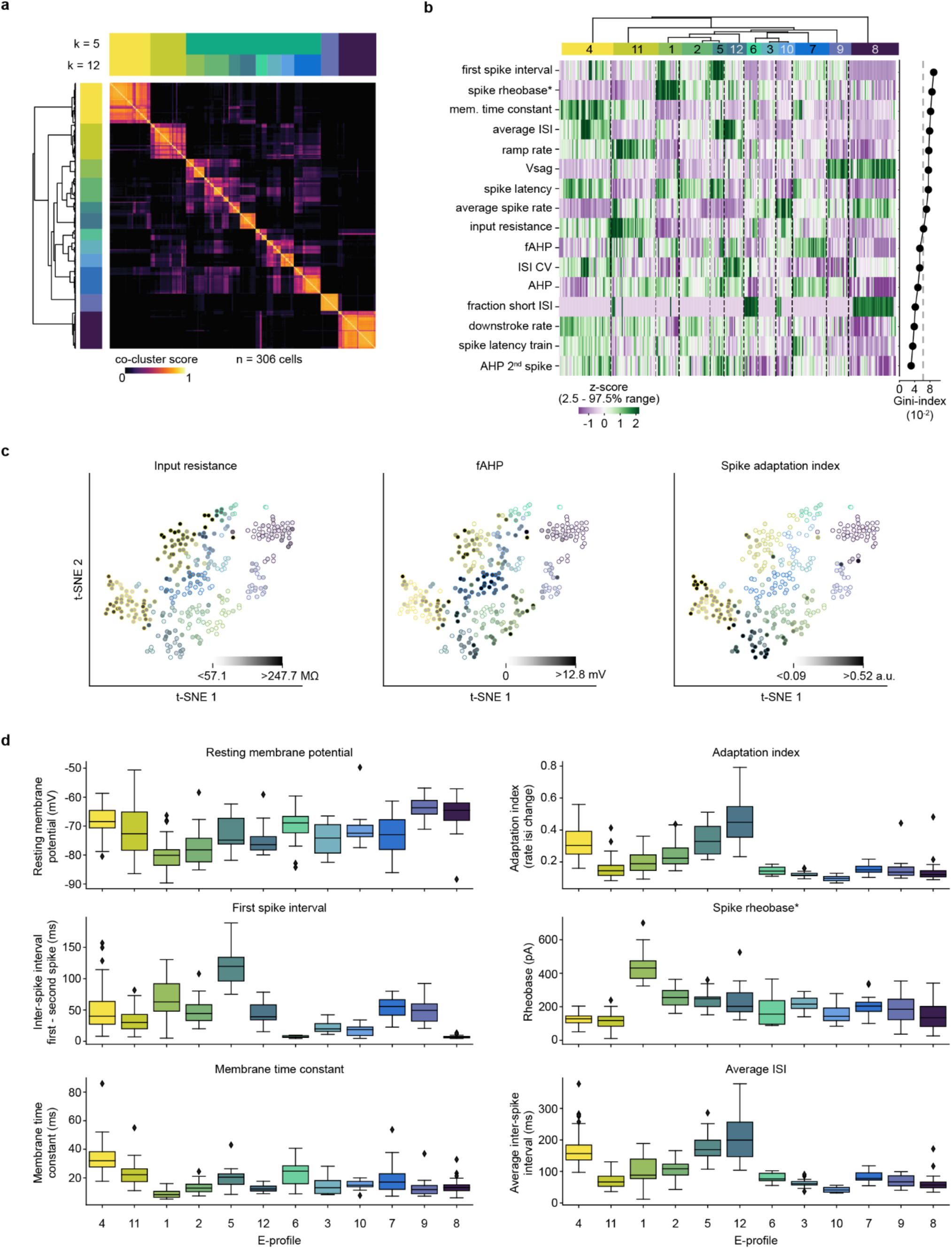
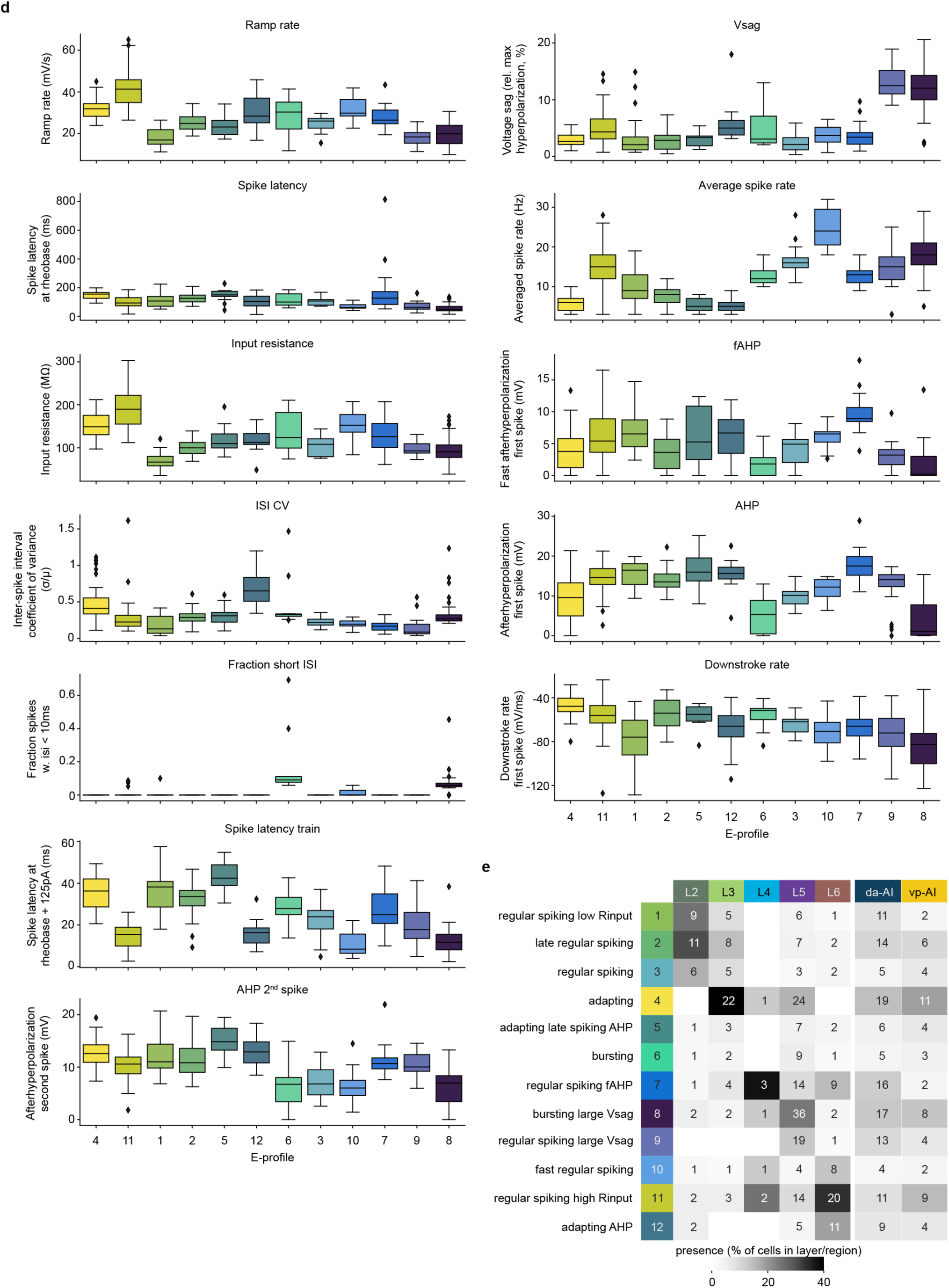
Distinct features of E-profiles. **a**, Hierarchical cluster analysis (HCA) on pair-wise co-cluster scores across 306 cells using the best performing electrical feature set. Dendrogram-based grouping is color-coded for 12 clusters (k = 12) based on optimal silhouette score, as well as for 5 clusters (k = 5). **b**, Z-scored electrical features (left), ranked by random forest classifier Gini-index (right, gray dashed line indicates equal feature contribution [1/16 features]). **c**, Representative electrical features contrasting E-profiles. The parameter amplitudes are shown in grayscale luminance of the filled dots, mapped onto the t-SNE plot in Fig. 4c. Edges of the filled dots are color-coded by E-profiles. **d**, All 16 electrical features used for the iCCA, as well as resting membrane potential and adaptation index. Boxplots are sorted in sequence as shown in the dendrogram from (**a**). **e**, Contingency table of cell allocations to either layer (left) or subregion (right) for each E-profile. The E-profiles are named based on their prominent electrical properties. Grayscale luminance: the percentage of cells in each E-profile allocated to a given layer or subregion. Numbers in gray boxes: n of observed cells (when n > 0).

**Extended Data Fig. 9:**
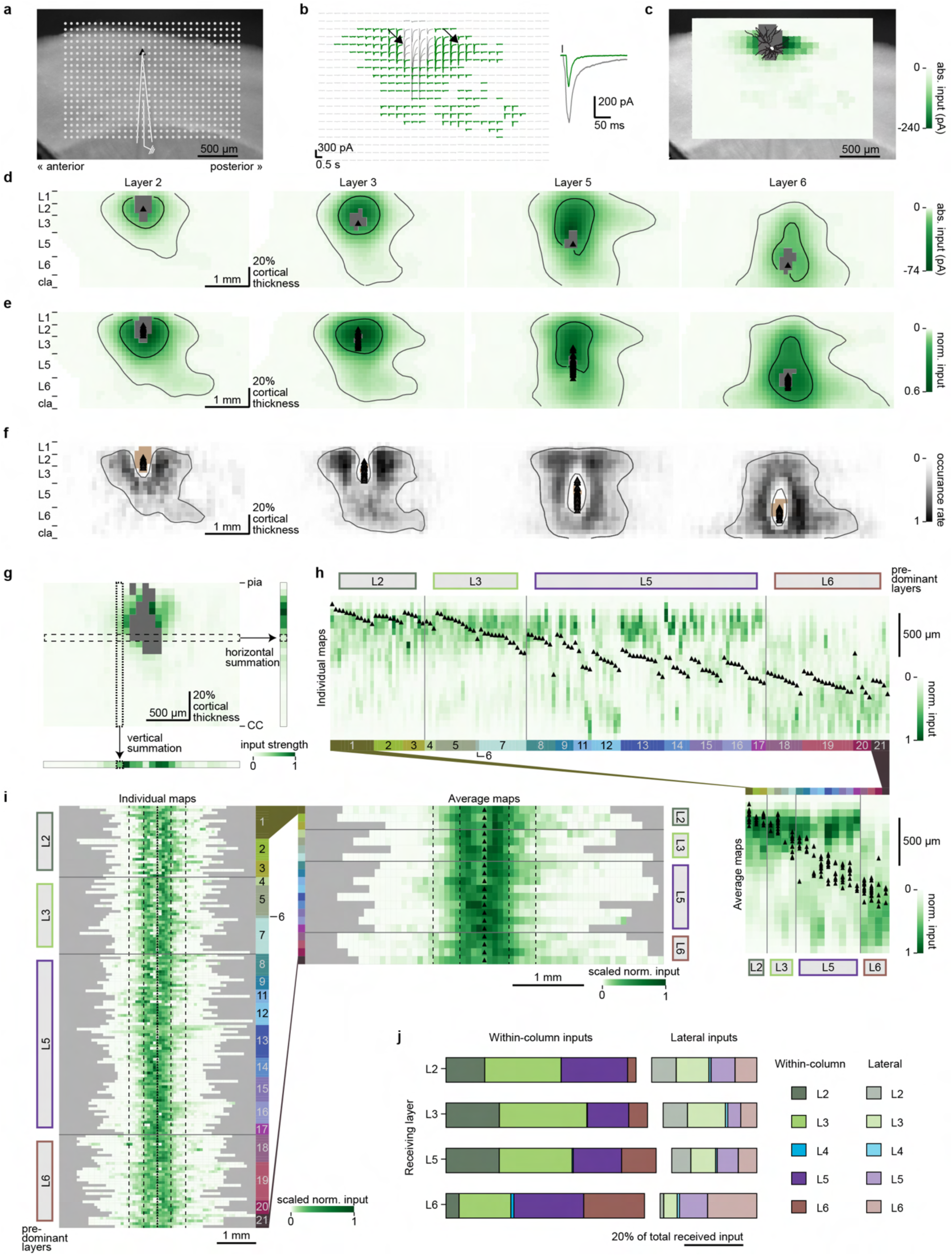
Laminar and intracolumnar analysis of LSPS maps. **a**, Example horizontal acute slice for LSPS recording. White dots: 75 × 75 µm stimulation grid; solid lines: the recording electrode; black triangle: soma location of the recorded cell. **b**, Trace map of current responses (left) corresponding to the stimulation sites in (**a**) and enlarged example traces of direct and synaptic responses (right, marked by arrowheads on the trace map). Sites with synaptic responses, direct responses, and no response are colored in green, dark gray, and light gray, respectively. Vertical tick: stimulation onset. **c**, Heatmap corresponding to the trace map in (**b**), overlaid with the acute slice shown in (**a**) and reconstructed dendritic morphology (black lines) of the recorded cell. Gray pixels: direct response sites. **d** and **e**, Average absolute input maps aligned by soma (**d**) and normalized input maps aligned by pia (**e**) of L2 (n = 18), L3 (n = 36), L5 (n = 64), and L6 (n = 35) neurons. Gray pixels: direct response sites; black triangles: soma locations. The average maps are Gaussian-filtered for visualization, shown in exponential color scale to highlight distal inputs, and contoured at 15^th^ and 85^th^ percentiles of all responses. **f**, Probability maps of input occurrence aligned by pia of the same neurons from (**d** and **e**). Probability maps are not Gaussian-filtered but contoured at 20% occurrence rate of their Gaussian-filtered counterparts to minimize noise. Brown pixels: direct response sites; black triangles: soma locations. **g**, 2D interpolated LSPS map of an example neuron and its vertically (bottom) and horizontally (right) collapsed 1D maps (Methods). Dotted column: example column on 2D map that sums up to the corresponding pixel (dotted pixel) on 1D map (bottom). Dashed row: example row on 2D map that sums up to the corresponding pixel (dashed pixel) on 1D map (right). All maps were normalized to their own maximum values and shown in linear color scale. **h**, Individual (top) and average (bottom right) horizontally collapsed maps grouped by M-types (n = 154). Individual maps are sorted within groups by soma cortical depth. Heatmaps are shown in linear scale. **i**, Individual (left) and averaged (upper right) vertically collapsed maps grouped by M-types (n = 154). Individual maps are sorted within groups by the number of non-zero pixels. Dashed lines indicate 95% dendritic density (inner) or maximum dendritic reach (outer). Heatmaps are shown in exponential scale to highlight distal inputs. For (**h** and **i**), colored boxes: M-types; trapezoids: connections of the same morphology types on the outer edges of individual and averaged maps; black triangles: soma locations. **j**, Normalized input strength arriving at each AI cortical layer. Within-column (saturated colors) and lateral (semitransparent colors) inputs are separated by gaps. cla, claustrum; CC: corpus callosum.

**Extended Data Fig. 10:**
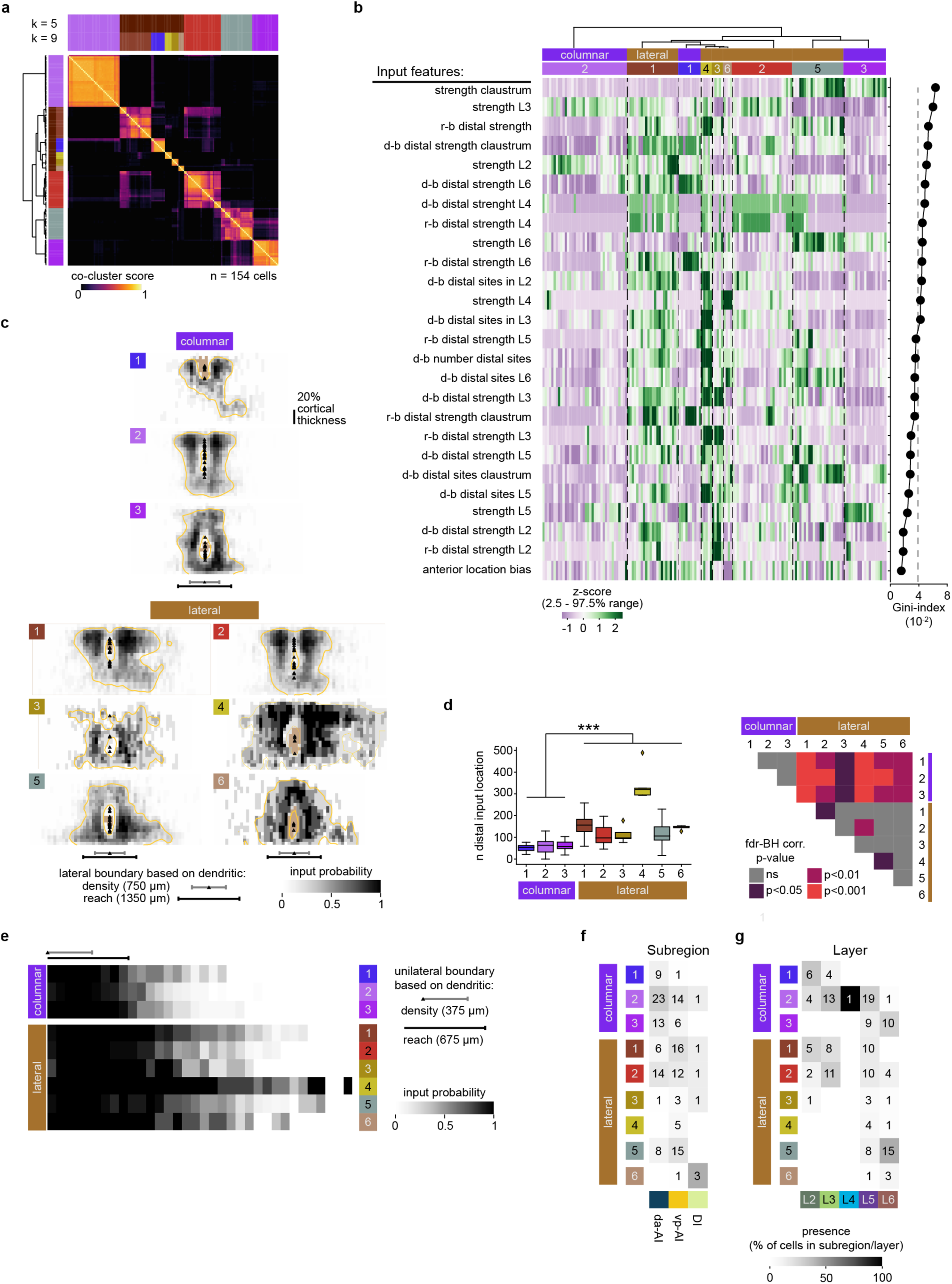
Local input features highlight lateral extent of input profiles. **a**, Hierarchical cluster analysis (HCA) on pair-wise co-cluster scores across 154 cells using the best performing input feature set. Dendrogram-based grouping is color-coded for 9 clusters (k = 9) based on optimal silhouette score, as well as for 5 clusters (k = 5). **b**, Z-scored input features (left), ranked by random forest classifier Gini-index (right, gray dashed line indicates equal feature contribution [1/26 features]). Input profile color-coding is consistent throughout this figure and Fig. 5. For lateral input features, the lateral border was defined by boundaries based on either 95% dendritic density (density-based, d-b) or maximum dendritic reach (reach-based, r-b). **c**, Probability maps of input occurrence aligned by pia for all I-profiles. Maps are interpolated vertically and normalized to the cortical thickness. Probability maps are not Gaussian-filtered but contoured at 20% occurrence rate of their Gaussian-filtered counterparts to minimize noise. Brown pixels: direct response sites; black triangles: soma locations; gray and black lateral whisker bars: lateral boundary thresholds used during feature extraction, based on dendritic density (750 µm) or reach (1350 µm), respectively. **d**, Number of distal input locations beyond the dendritic density-based borders (left), and the corresponding heatmap of p-values from pair-wise comparisons between I-profiles (right, Kruskal-Wallis H (8) = 80.77, p < 0.001, with post hoc Dunn’s comparisons with false-discovery rate Benjamini-Hochberg correction, fdr-BH corr.). Mann-Whitney U tests were used to further compare columnar and lateral input classes; U = 675.5, ***: p < 0.001. **e**, Input probability as a function of distance from soma. **f** and **g**, Contingency tables of I-profiles and -classes versus subregion (**f**) and layer (**g**). Gray luminance: the percentage of cells in each I-profile allocated to a given aforementioned modality. Numbers in gray boxes: n of observed cells (when n > 0).

**Extended Data Fig. 11:**
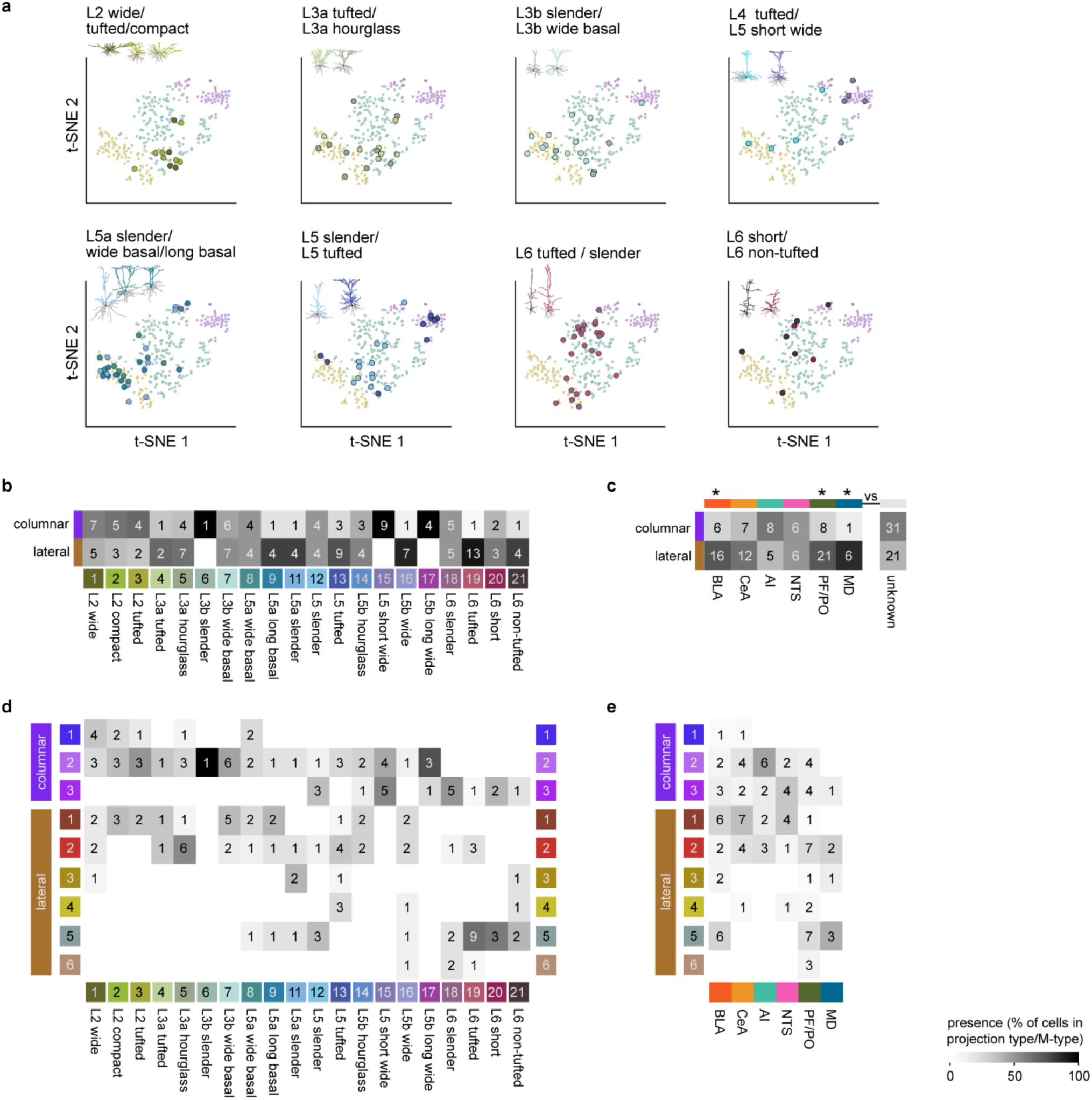
Electrical and input profiles are related to specific morphologies and projection targets. **a**, M-types mapped onto the tSNE plot in Fig. 4e. **b** and **c**, Contingency tables of the input classes versus M-types (**b**) and projection targets (**c**). **d** and **e**, Contingency tables of the input profiles versus M-types (**d**) and projection targets (**e**). For (**b**–**e**), grayscale luminance: the percentage of cells with either input profile/class within a given M-type or projection target type. Numbers in gray boxes: n of observed cells (when n > 0).

**Extended Data Fig. 12:**
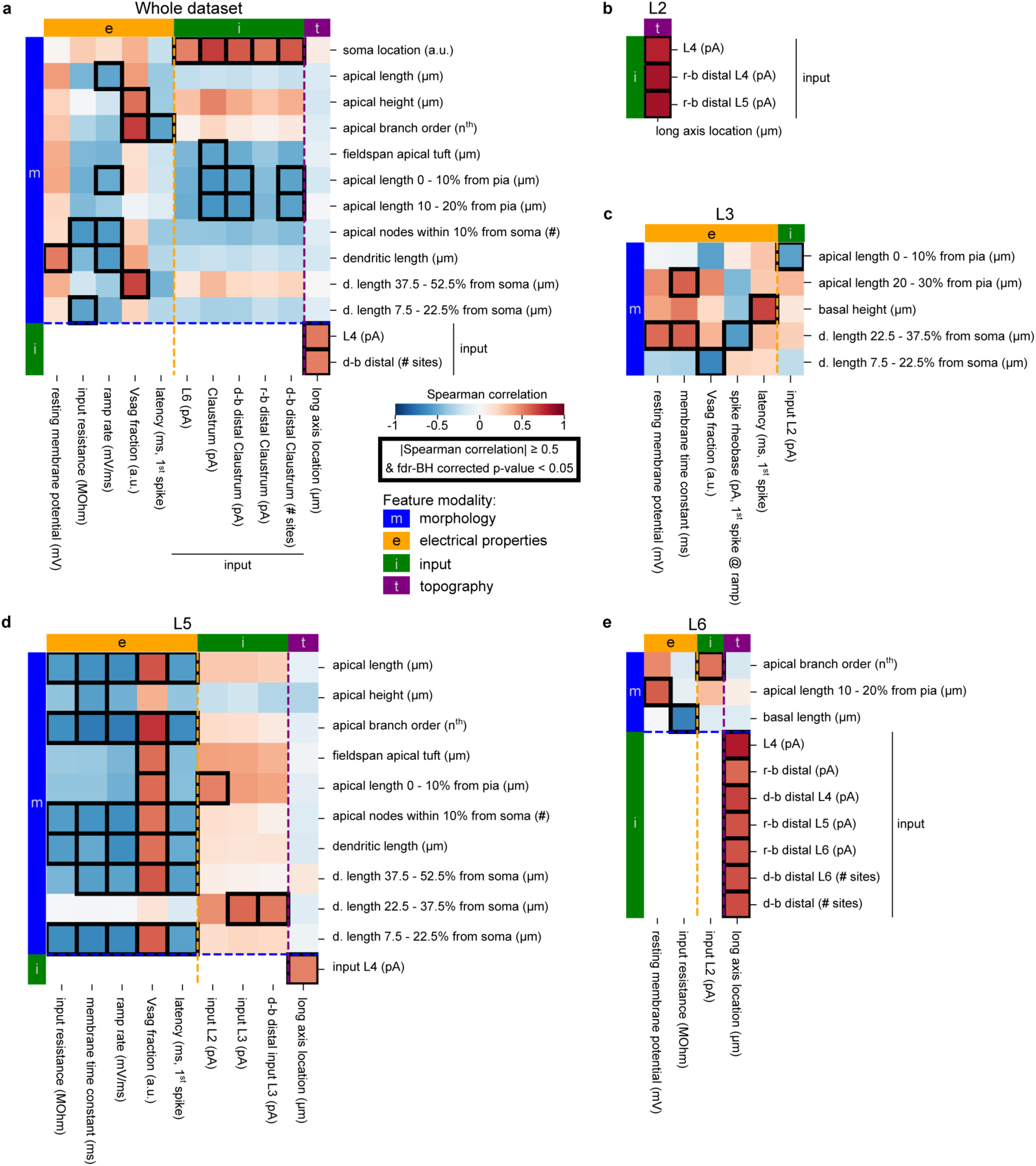
Cross-modal feature correlations. **a**–**e**, Feature pair-wise Spearman correlation between morphology (m, blue), electrical (e, yellow), input (i, green), and topographic (t, purple) features. Spearman correlations were computed based on the whole dataset (**a**), or subsets based on layer allocation of the individual cells (**b**–**e**). Note that topology (t, long axis location) correlated with input (i), but not morphology (m) or electrical (e), features in the whole dataset and layer subsets (L2, L5, and L6) (**a**, **b**, **d**, and **e**). Feature pairs with an absolute Spearman correlation ≥ 0.5 and false-discovery rate Benjamini-Hochberg (fdr-BH) corrected p-value < 0.05 are boxed. Only feature pairs yielding significant correlations between any two modalities are shown (Methods, whole dataset: n = 25, L2: n = 3, L3: n = 7, L5: n = 36, L6: n = 10). For lateral input features, the lateral border was defined by boundaries based on either 95% dendritic density (density-based, d-b) or maximum dendritic reach (reach-based, r-b).

**Extended Data Fig. 13:**
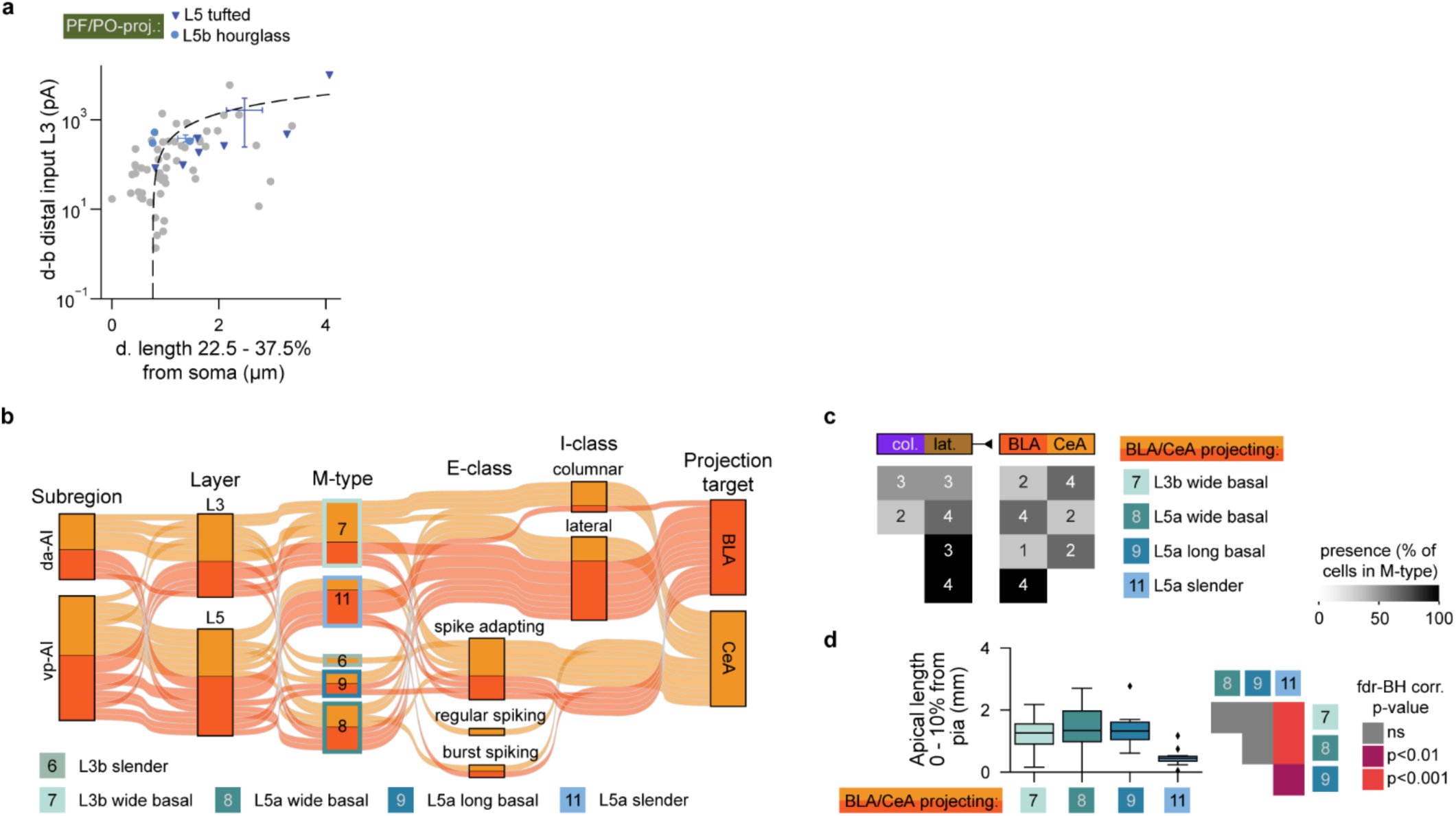
Informative features for cell type identification. **a**, For PF/PO-projecting *L5 tufted* (M-type 13) and *L5b hourglass* (M-type 14) neurons, distal L3 inputs are monotonically correlated with dendritic length, normalized to cortical thickness, within a 22.5 – 37.5% bin from soma towards pia (Spearman correlation = 0.52, semi-log scale). L3 distal input is defined as the summed L3 → L5 inputs beyond the dendritic density-based (d-b) boundaries. A linear fit (dashed line, 943.85 *x* – 704.55; r^2^ = 0.28) was applied. Color-coded whiskers: mean and standard error of plotted features for PF/PO-projecting *L5 tufted* (M-type 13) and *L5b hourglass* (M-type 14) neurons. **b**, Parallel categories diagram of L3b and L5a amygdala-projecting neurons and their allocations to subregions, M-types, E-classes, and I-classes. **c**, Contingency table of the same subset of M-types in (**b**) versus input classes or projection targets. Grayscale luminance: the percentage of cells within each M-type allocated to a given input class or projection target. Numbers in gray boxes: n of observed cells (when n > 0). **d**, Quantification of apical length within 10% cortical thickness bin from pia (left) and the corresponding heatmap of p-values from the pair-wise statistical comparisons between the same subset of M-types in (**b**) (right, Kruskal Wallis test H (3) = 14.08, p = 0.003, with post hoc Dunn’s comparisons with false-discovery rate Benjamini-Hochberg correction (fdr-BH corr.).

**Extended Data Fig. 14:**
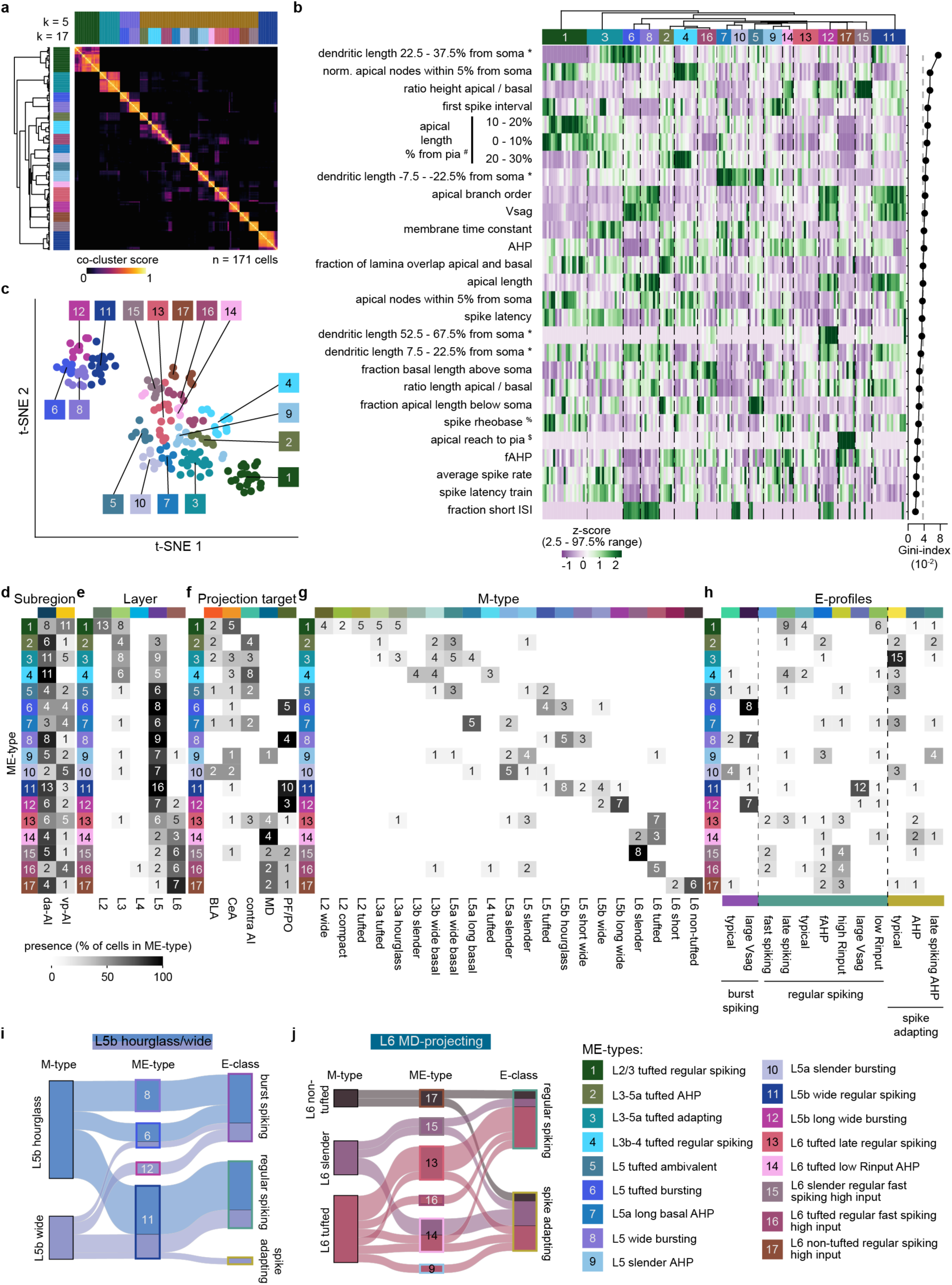
Analysis of combined morpho-electrical features and subsequent cell groups. **a**, Hierarchical cluster analysis (HCA) on pair-wise co-cluster scores across 171 cells using both morphological and electrical features. Dendrogram-based grouping is color-coded for 17 clusters (k = 17) based on optimal silhouette score, as well as for 5 clusters (k = 5). **b**, Z-scored electrical and morphological features (left), ranked by random forest classifier Gini-index (right, gray dashed line indicates equal feature contribution [1/27 features]). **c**, t-SNE distribution of morpho-electrical types (ME-types). For (**b** and **c**), colored boxes with numbers: ME-types color-coded for k = 17 in (**a**). **d**–**h**, Contingency table of cell allocations to subregion (**d**), layer (**e**), projection target (**f**), M-type (**g**), or E-profile (**h**). Note in (**f**), most ME-types only contained IT (e.g., BLA-, CeA-, or contra AI-projecting) or ET (e.g., MD- or PF/PO thalamus-projecting) neurons, suggesting IT and ET neurons (Fig. 6c) can be readily resolved using ME-types. Grayscale luminance: the percentage of cells in each ME-type allocated to a given modality. Numbers in gray boxes: n of observed cells (when n > 0). **i** and **j**, Parallel categories diagrams of *L5b hourglass* (M-type 14) and *L5b wide* (M-type 16) neurons (**i**; n = 23 cells) and L6 MD-projecting neurons (**j**; n = 14 cells) across M-types, ME-types, and E-classes. Several M-types and E-classes can be sub-specified into multiple ME-types. ME-type legend (bottom right) is shared across all panels.

**Extended Data Fig. 15:**
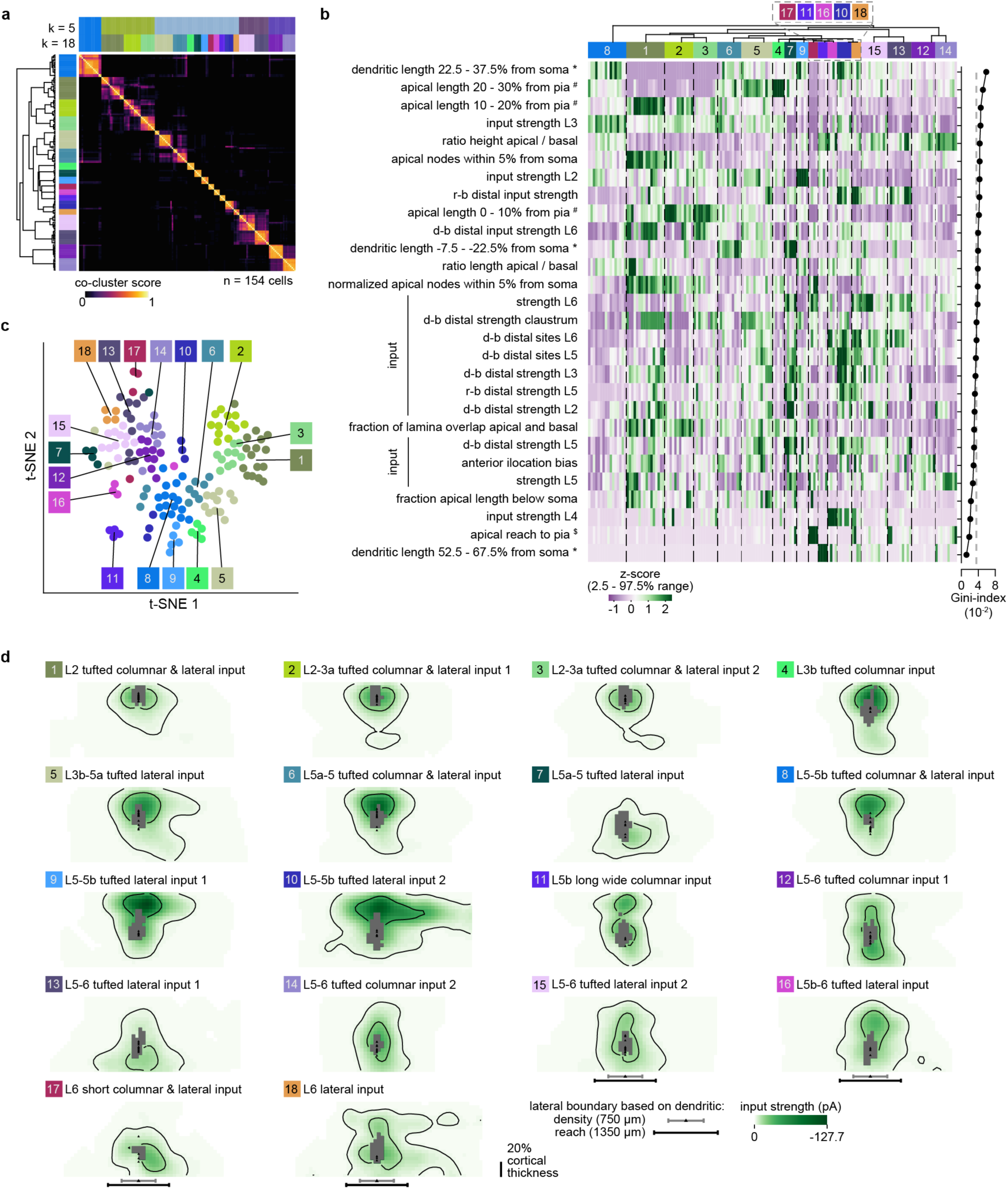
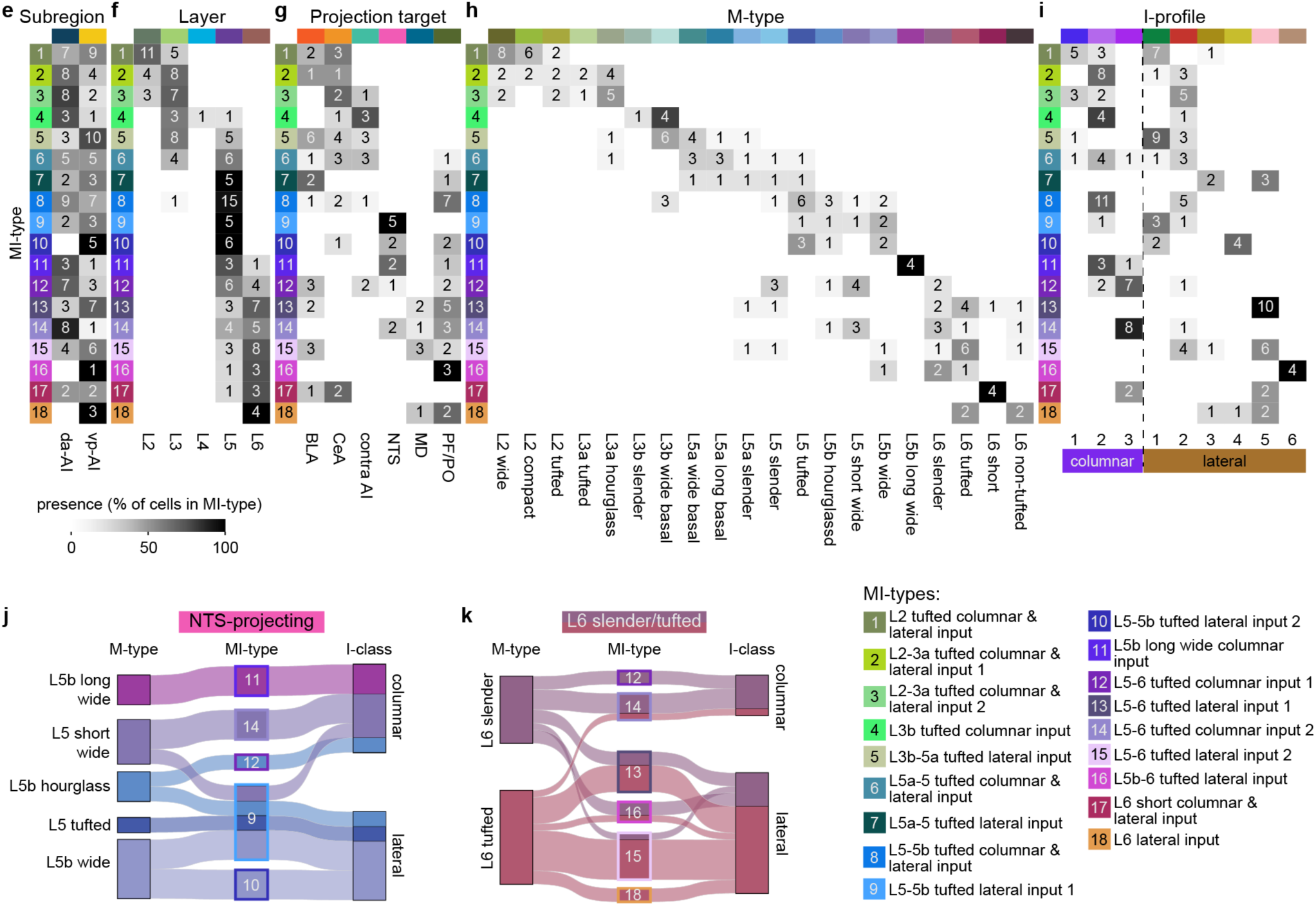
Analysis of combined morpho-input features and subsequent cell groups. **a**, Hierarchical cluster analysis (HCA) on pair-wise co-cluster scores across 154 cells using both morphological and input features. Dendrogram-based grouping is color-coded for 18 clusters (k = 18) based on optimal silhouette score, as well as for 5 clusters (k = 5). **b**, Z-scored morphological and input features (left), ranked by random forest classifier Gini-index (right, gray dashed line indicates equal feature contribution [1/28 features]). For distal input features, the border was defined by boundaries based on either 95% dendritic density (density-based, d-b) or maximum dendritic reach, (reach-based, r-b). * Dendritic length located within a 15% local cortical thickness bin positioned relative to the soma, normalized to local cortical thickness. ^#^ Apical length within a 10% local cortical thickness bin positioned relative to the pia, normalized to total apical length. ^$^ Vertical distance between pia and closest apical dendrite, normalized to local cortical thickness. **c**, t-SNE distribution of morpho-input types (MI-types). **d**, Average input maps grouped by MI-types, aligned by pia. Maps are vertically interpolated and normalized to the cortical thickness. Gray and black lateral whisker bars: lateral boundary thresholds, used during feature extraction, based on dendritic density (750 µm) or reach (1350 µm), respectively; gray pixels: direct response sites; black triangles: soma locations. For (**b**–**d**), colored boxes with numbers: MI-types color-coded for k = 18 in (**a**). **e**–**i**, Contingency tables of MI-types versus subregion (**e**), layer (**f**), projection target (**g**), M-type (**h**) and I-profile (**i**). Grayscale luminance: the percentage of cells in each MI-type allocated to a given aforementioned modality. Numbers in gray boxes: n of observed cells (when n > 0). **j**, Parallel categories diagrams of NTS-projecting neurons (n = 12). **k**, Parallel categories diagrams of *L6 slender* (M-type 18) and *L6 tufted* (M-type 19) neurons (n = 24). Note in (**i**-**k**), a majority of MI-types (12/18) retained predominance in either columnar or lateral input classes. MI-type legend (bottom right) is shared across all panels.

**Extended Data Fig. 16:**
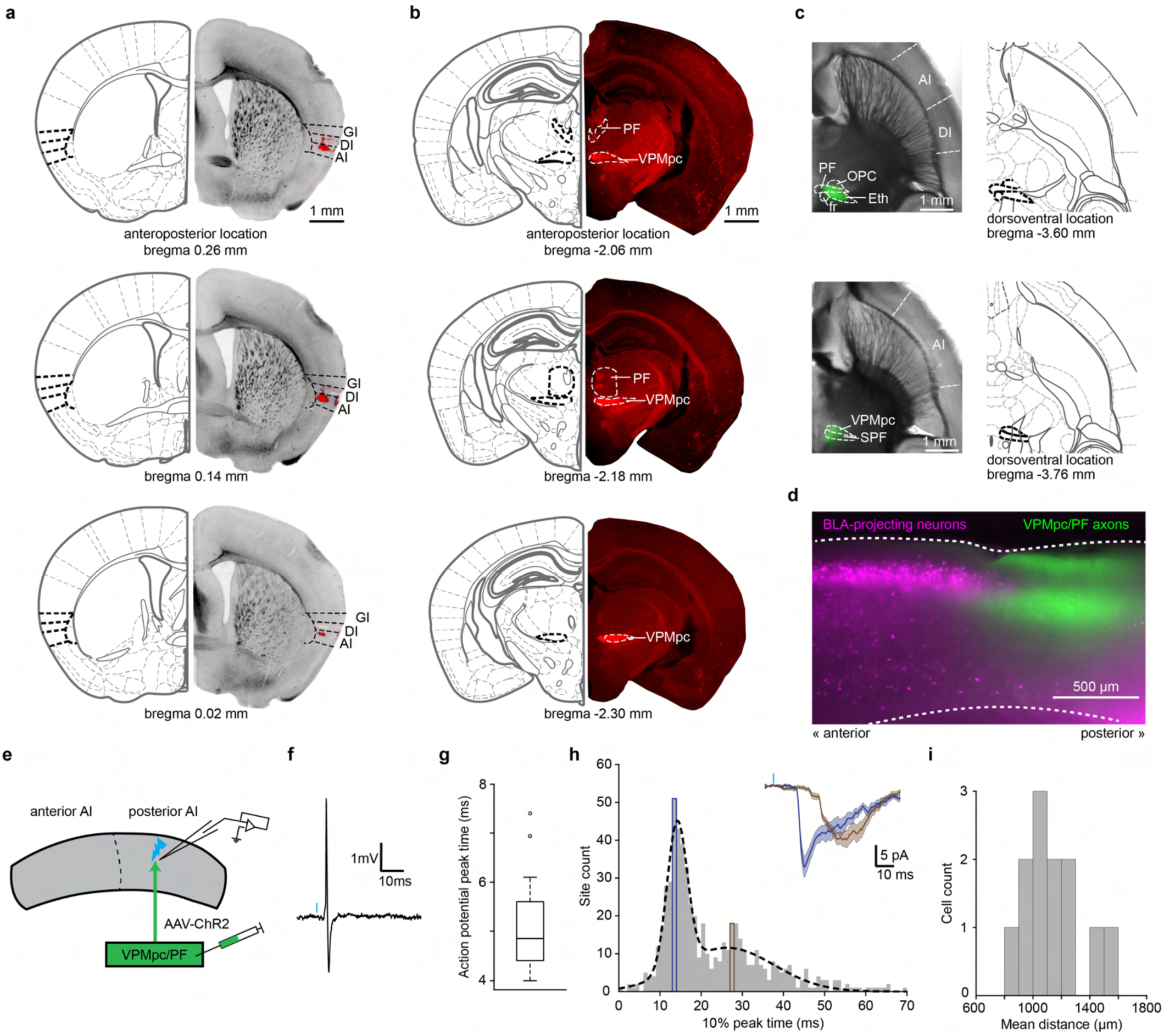
Anatomical analysis of the VPMpc/PF-AI circuit and kinetics of the synaptic responses. **a**, Example AI injection site of retrograde beads. Brightfield images (right) of all coronal sections (thickness: 50 µm) containing the injection site (red fluorescence) are shown and matched to equivalent sections from the Franklin and Paxinos atlas^45^ (left). **b**, Example epifluorescent images of bead locations in VPMpc and PF (right) and matched sections from the Franklin and Paxinos atlas (left), from the same animal shown in (**a**). Bead location images were linearly adjusted for brightness and contrast. **c**, Brightfield examples of acute horizontal brain slices (thickness: 350 µm) containing the thalamic ChR2 injection (left, green fluorescence) and matched sections from the Franklin and Paxinos atlas (right). All slices containing the injection are shown. For (**a**–**c**), positions relative to bregma are noted according to the best matched atlas sections. **d**, Overlay of retrogradely-labeled BLA-projecting neurons (magenta) and ChR2-positive VPMpc/PF axons (green). **e**, Schematic of ChR2 stimulation and loose-patch recording in the posterior AI. Blue flash: optogenetic stimulation. **f**, Example loose-patch trace acquired in an experiment illustrated in (**e**). Blue tick: stimulation onset. **g**, Time delay from stimulation onset to action potential peak (n = 14 responsive, out of 29 total recorded cells). **h**, Histogram of time delay from posterior stimulation onset to anterior response when it reaches 10% peak response and average response trace (inset) at the 14 – 15 ms (blue box) or 27 – 28 ms (brown box) bin. Blue tick: stimulation onset. Shaded area: standard error. Dotted line: bimodal Gaussian fit of the distribution. **i**, Histogram of mean anterior-to-posterior distance between recorded soma and response-yielding posterior stimulation sites (Fig. 7b). The closest distances from the soma to the stimulation columns containing each response-yielding site were averaged and plotted. PF, Parafascicular thalamus; OPC, Oval paracentral thalamus; Eth, Ethmoid thalamus; fr, fasciculus retroflexus; VPMpc, Ventroposteromedial thalamus, parvicellular part; SPF, Subparafascicular thalamus.

**Extended Data Table 1:** Iterative consensus cluster analysis (iCCA) parameters and performance across analyzed modalities. The optimal cluster number, silhouette score, and number of features in the feature set are based on the clustering results of the best-performing feature set (Methods). Five independent instances of random forest classifiers were trained on 75% of the dataset and the F1-score was calculated based on the classifiers’ predictions of the 25% withheld dataset.

|  |  | iCCA part 1 |  | iCCA part 2 |  |  |  |  |  |
| --- | --- | --- | --- | --- | --- | --- | --- | --- | --- |
| Modality | Features tested | Principal components (median, ± min. – max.) | Cluster number PCA-based | Number of informative features | Unique feature sets tested | Optimal cluster number | Silhouette score | Number of features in set | Random forest classifier F1 score (% median, ± min. – max.) |
| M-types | 67 | 20 ± 19 – 23 | 20 | 54 | 22 | 21 | 0.43 | 29 | 84.8 ± 82.9 – 85.3 |
| E-profiles | 29 | 10 ± 9 – 11 | 10 | 27 | 20 | 12 | 0.5 | 16 | 82.5 ± 78.4 – 85.3 |
| I-profiles | 45 | 15 ± 14 – 16 | 7 | 45 | 33 | 9 | 0.6 | 26 | 81.3 ± 75.0 – 91.2 |
| ME-types | 45 | 15 ± 14 – 16 | 7 | 44 | 21 | 17 | 0.58 | 27 | 71.8 ± 56.7 – 78.5 |
| MI-types | 55 | 20 ± 19 – 22 | 9 | 54 | 17 | 18 | 0.55 | 28 | 65.0 ± 61.9 – 72.1 |

## Notes

### Competing Interest Statement

The authors have declared no competing interest.

