## Supplementary Information for "Neuronal architecture of the mouse insular cortex underlying its diverse functions"

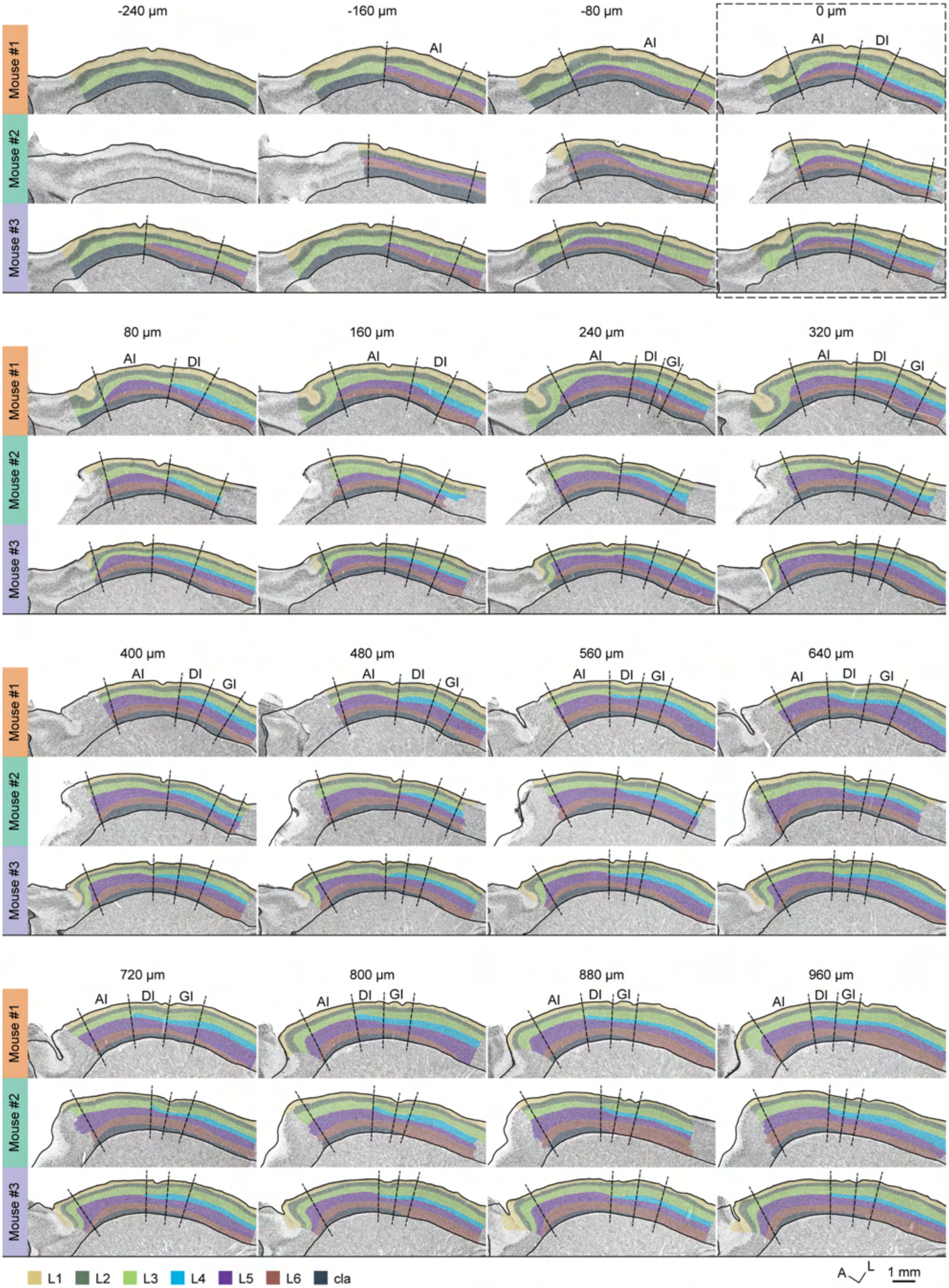
-240 µm
-160 µm
-80 µm
0 µm
80 µm
160 µm
240 µm
320 µm
400 µm
480 µm
560 µm
640 µm
720 µm
800 µm
880 µm
960 µm
Mouse #1
Mouse #2
Mouse #3
AI
DI
GI
L1
L2
L3
L4
L5
L6
cla
A
L
1 mm

1506

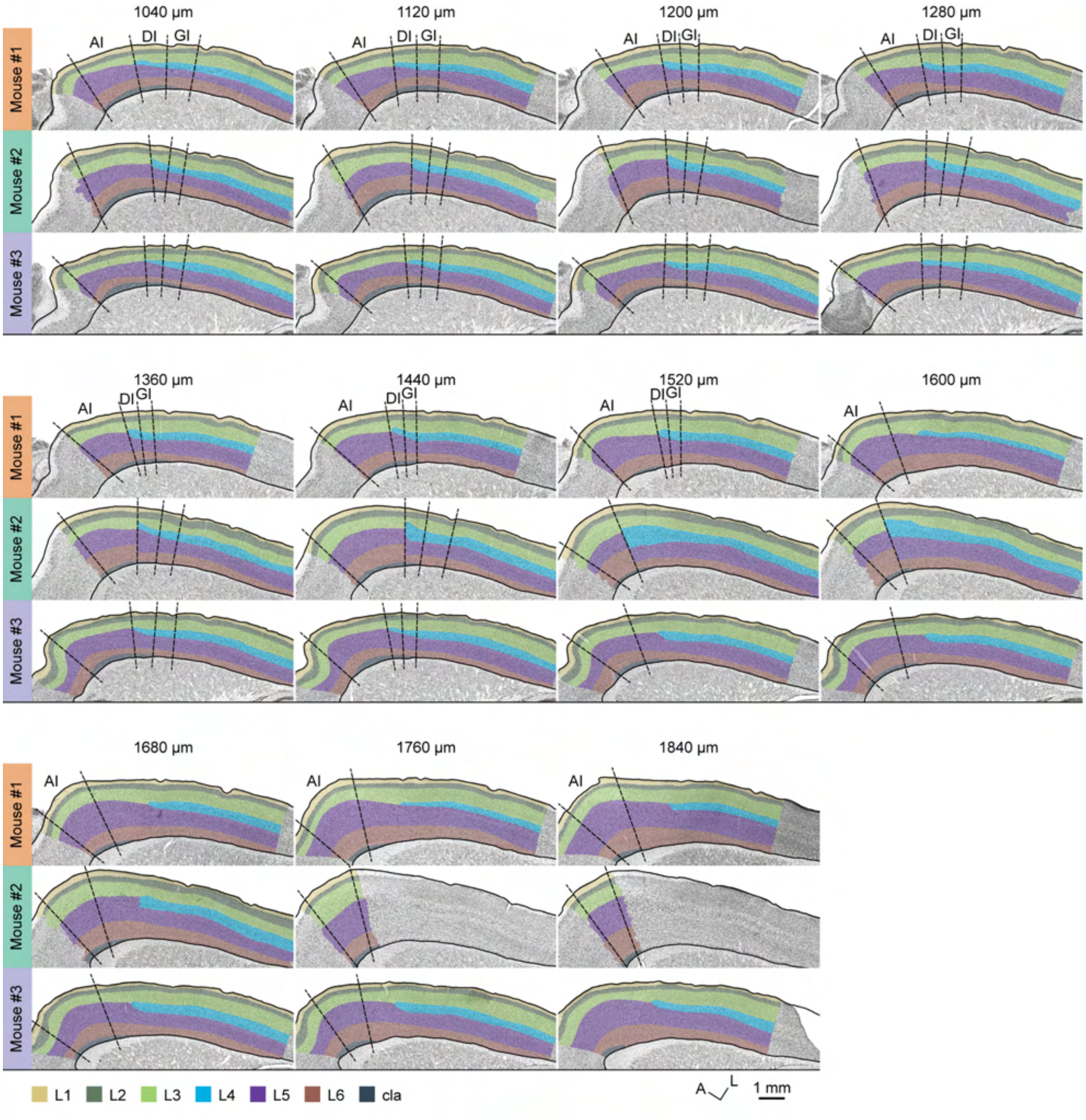

**Supplementary Fig. 1: Complete sets of consensus manual annotations of the insular cortex.** An alternating (every other) series of 40 µm-thick horizontal sections collected from three *PV-Cre*$^{+/-}$;*Ai9*$^{+/-}$ mice processed for Nissl histology. Overlay shows the average consensus from two independent expert annotators demarcating the locations of all cortical layers within and adjacent to the insular cortex, as well as the boundaries of its subregions. Sections from each animal are arranged in rows along the ventral-to-dorsal cutting axis (top-left to bottom-right) and aligned across animals based on the ventral-most section in which layer 4 (L4) is conspicuous (dashed box, top row, 4th column). The position of each section relative to this L4 anchor along the ventral-to-dorsal cutting axis is indicated above each column in μm. A, anterior; L, lateral; AI, agranular insula; DI, dysgranular insula; GI, granular insula; cla, claustrum; L1-L6, cortical layers 1-6.

1518

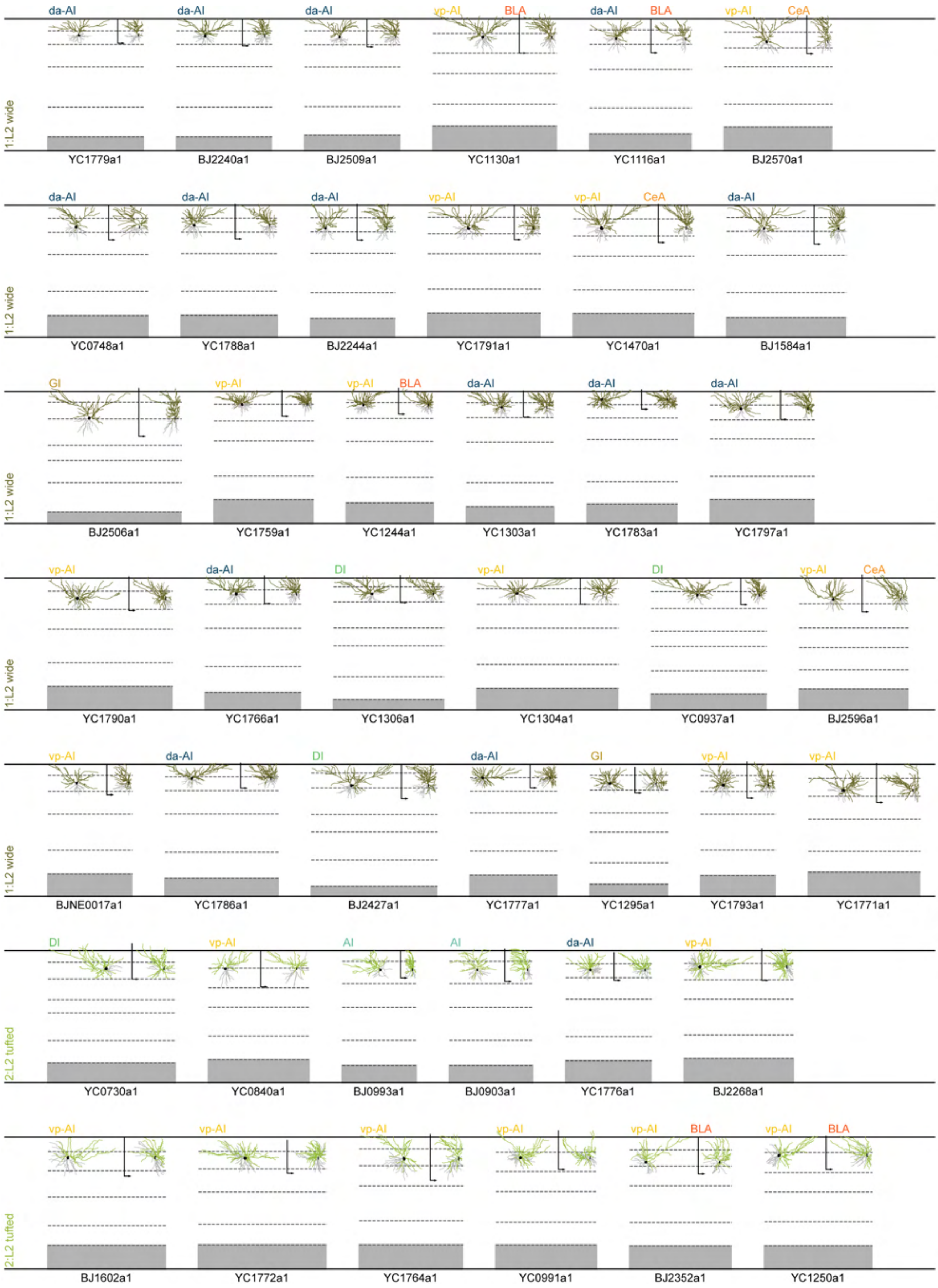
1:L2 wide
da-AI
da-AI
da-AI
vp-AI
BLA
da-AI
BLA
vp-AI
CeA
YC1779a1
BJ2240a1
BJ2509a1
YC1130a1
YC1116a1
BJ2570a1
1:L2 wide
da-AI
da-AI
da-AI
vp-AI
vp-AI
CeA
da-AI
YC0748a1
YC1788a1
BJ2244a1
YC1791a1
YC1470a1
BJ1584a1
1:L2 wide
GI
vp-AI
vp-AI
BLA
da-AI
da-AI
da-AI
BJ2506a1
YC1759a1
YC1244a1
YC1303a1
YC1783a1
YC1797a1
1:L2 wide
vp-AI
da-AI
DI
vp-AI
DI
vp-AI
CeA
YC1790a1
YC1766a1
YC1306a1
YC1304a1
YC0937a1
BJ2596a1
1:L2 wide
vp-AI
da-AI
DI
da-AI
GI
vp-AI
vp-AI
BJNE0017a1
YC1786a1
BJ2427a1
YC1777a1
YC1295a1
YC1793a1
YC1771a1
2:L2 tufted
DI
vp-AI
AI
AI
da-AI
vp-AI
YC0730a1
YC0840a1
BJ0993a1
BJ0903a1
YC1776a1
BJ2268a1
2:L2 tufted
vp-AI
vp-AI
vp-AI
vp-AI
vp-AI
BLA
vp-AI
BLA
BJ1602a1
YC1772a1
YC1764a1
YC0991a1
BJ2352a1
YC1250a1

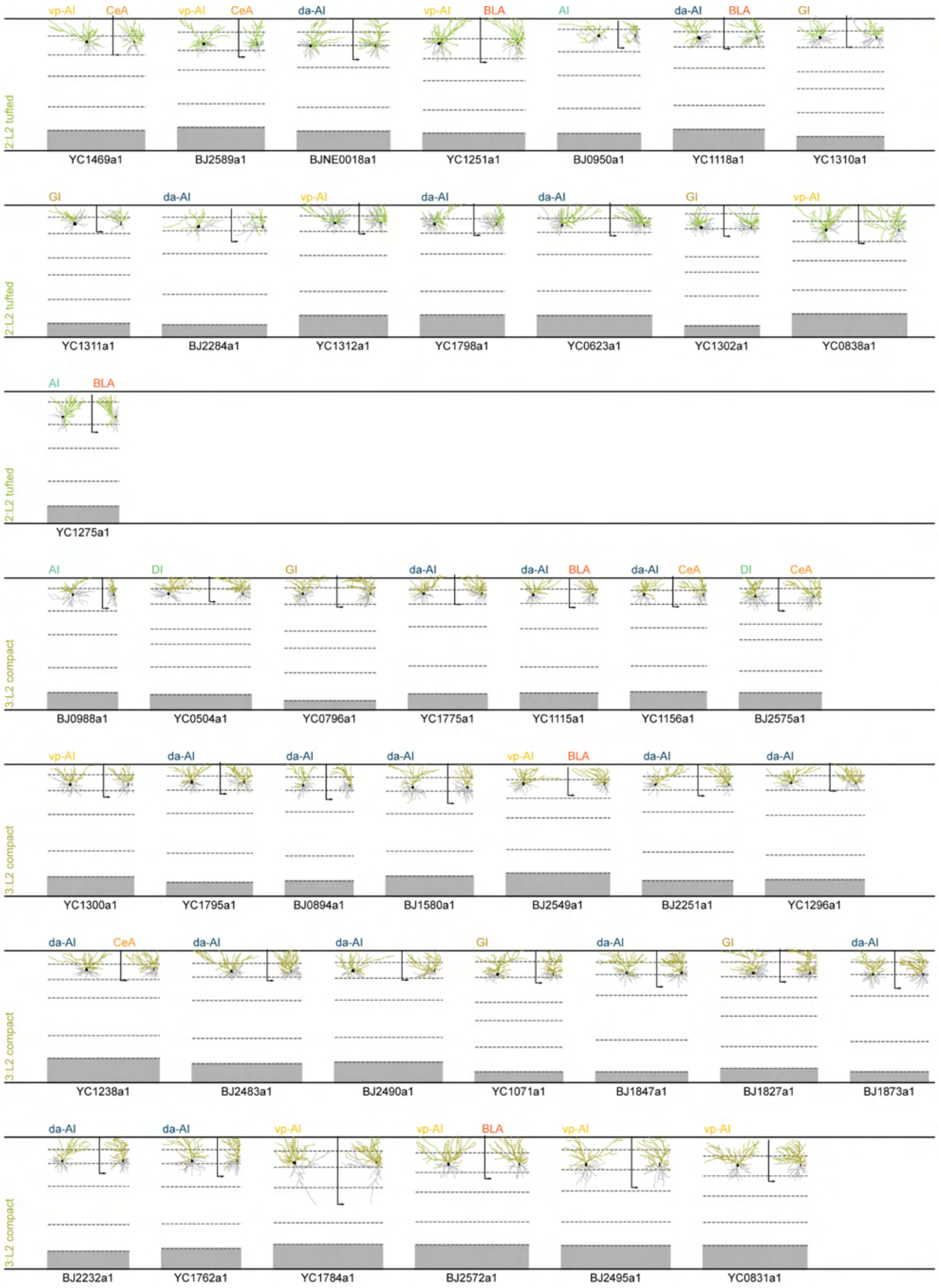
vp-AI CeA
YC1469a1
vp-AI CeA
BJ2589a1
da-AI
BJNE0018a1
vp-AI BLA
YC1251a1
AI
BJ0950a1
da-AI BLA
YC1118a1
GI
YC1310a1
2:L2 tufted
GI
YC1311a1
da-AI
BJ2284a1
vp-AI
YC1312a1
da-AI
YC1798a1
da-AI
YC0623a1
GI
YC1302a1
vp-AI
YC0838a1
2:L2 tufted
AI BLA
YC1275a1
2:L2 tufted
AI
BJ0988a1
DI
YC0504a1
GI
YC0796a1
da-AI
YC1775a1
da-AI BLA
YC1115a1
da-AI CeA
YC1156a1
DI CeA
BJ2575a1
3:L2 compact
vp-AI
YC1300a1
da-AI
YC1795a1
da-AI
BJ0894a1
da-AI
BJ1580a1
vp-AI BLA
BJ2549a1
da-AI
BJ2251a1
da-AI
YC1296a1
3:L2 compact
da-AI CeA
YC1238a1
da-AI
BJ2483a1
da-AI
BJ2490a1
GI
YC1071a1
da-AI
BJ1847a1
GI
BJ1827a1
da-AI
BJ1873a1
3:L2 compact
da-AI
BJ2232a1
da-AI
YC1762a1
vp-AI
YC1784a1
vp-AI BLA
BJ2572a1
vp-AI
BJ2495a1
vp-AI
YC0831a1
3:L2 compact

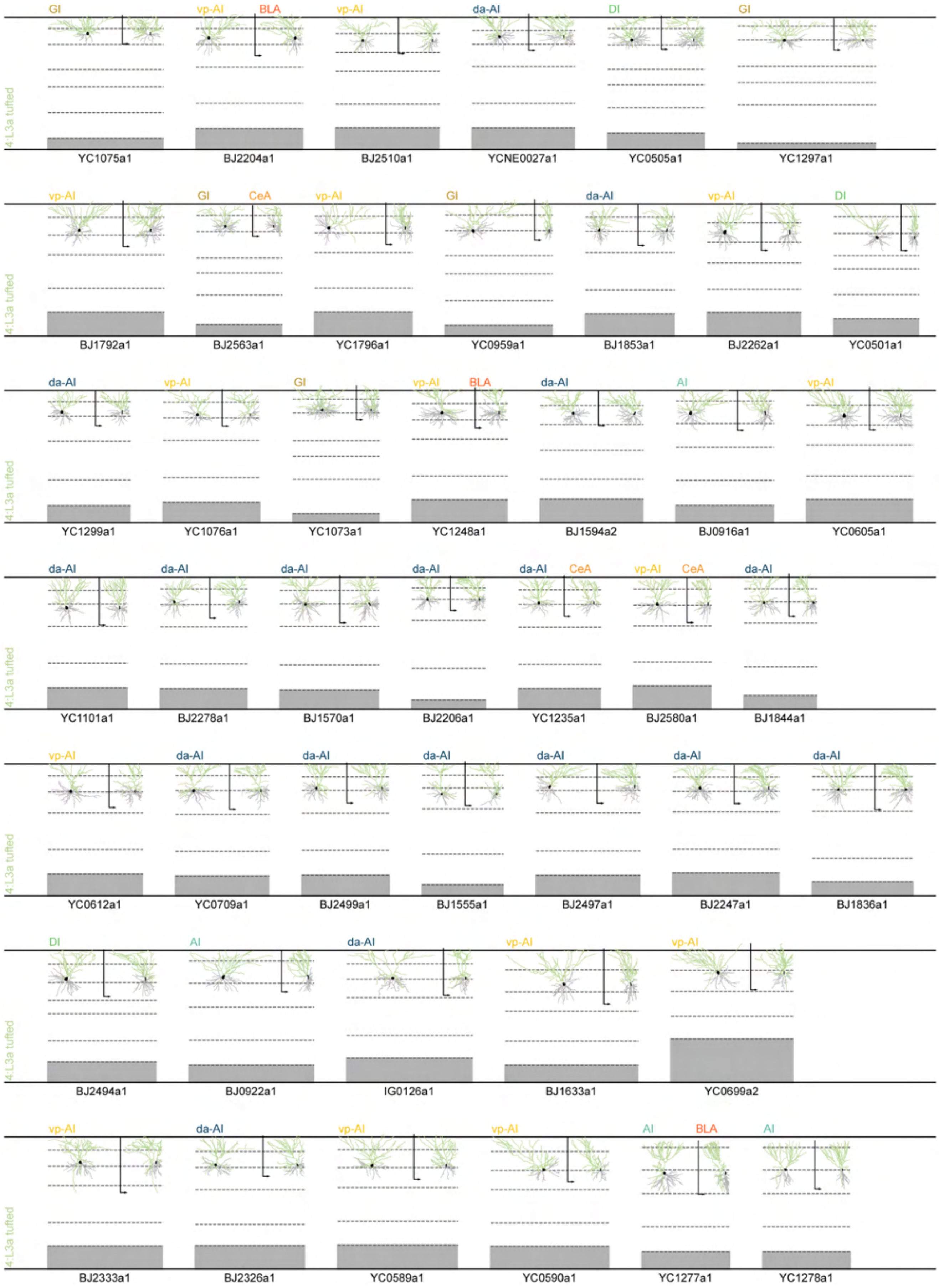
GI
vp-AI
BLA
vp-AI
da-AI
DI
GI
YC1075a1
BJ2204a1
BJ2510a1
YCNE0027a1
YC0505a1
YC1297a1
vp-AI
GI
CeA
vp-AI
GI
da-AI
vp-AI
DI
BJ1792a1
BJ2563a1
YC1796a1
YC0959a1
BJ1853a1
BJ2262a1
YC0501a1
da-AI
vp-AI
GI
vp-AI
BLA
da-AI
AI
vp-AI
YC1299a1
YC1076a1
YC1073a1
YC1248a1
BJ1594a2
BJ0916a1
YC0605a1
da-AI
da-AI
da-AI
da-AI
da-AI
CeA
vp-AI
CeA
da-AI
YC1101a1
BJ2278a1
BJ1570a1
BJ2206a1
YC1235a1
BJ2580a1
BJ1844a1
vp-AI
da-AI
da-AI
da-AI
da-AI
da-AI
da-AI
YC0612a1
YC0709a1
BJ2499a1
BJ1555a1
BJ2497a1
BJ2247a1
BJ1836a1
DI
AI
da-AI
vp-AI
vp-AI
BJ2494a1
BJ0922a1
IG0126a1
BJ1633a1
YC0699a2
vp-AI
da-AI
vp-AI
vp-AI
AI
BLA
AI
BJ2333a1
BJ2326a1
YC0589a1
YC0590a1
YC1277a1
YC1278a1

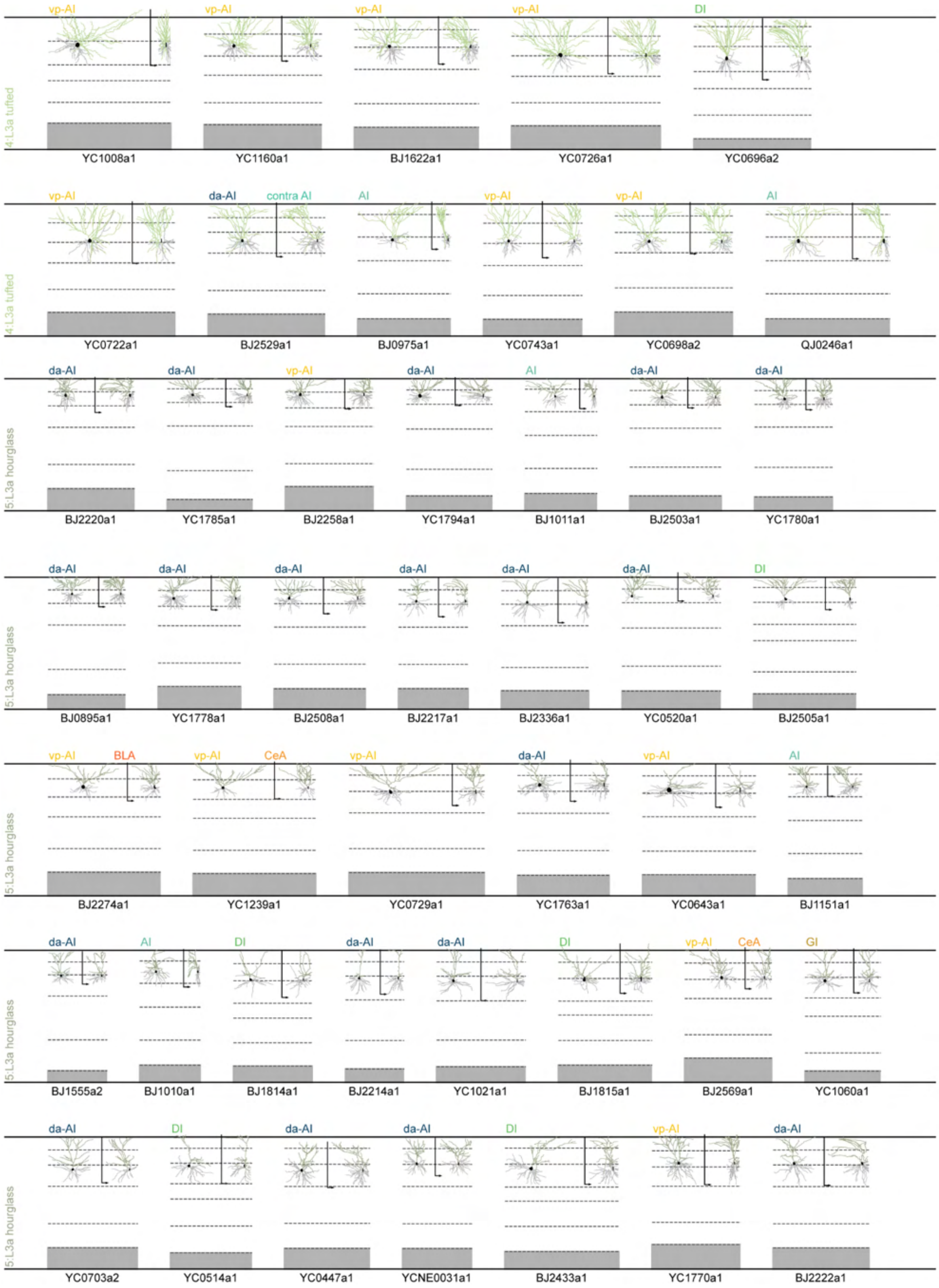
4:L3a tufted
vp-AI
YC1008a1
vp-AI
YC1160a1
vp-AI
BJ1622a1
vp-AI
YC0726a1
DI
YC0696a2
4:L3a tufted
vp-AI
YC0722a1
da-AI
contra AI
BJ2529a1
AI
BJ0975a1
vp-AI
YC0743a1
vp-AI
YC0698a2
AI
QJ0246a1
5:L3a hourglass
da-AI
BJ2220a1
da-AI
YC1785a1
vp-AI
BJ2258a1
da-AI
YC1794a1
AI
BJ1011a1
da-AI
BJ2503a1
da-AI
YC1780a1
5:L3a hourglass
da-AI
BJ0895a1
da-AI
YC1778a1
da-AI
BJ2508a1
da-AI
BJ2217a1
da-AI
BJ2336a1
da-AI
YC0520a1
DI
BJ2505a1
5:L3a hourglass
vp-AI
BLA
BJ2274a1
vp-AI
CeA
YC1239a1
vp-AI
YC0729a1
da-AI
YC1763a1
vp-AI
YC0643a1
AI
BJ1151a1
5:L3a hourglass
da-AI
BJ1555a2
AI
BJ1010a1
DI
BJ1814a1
da-AI
BJ2214a1
da-AI
YC1021a1
DI
BJ1815a1
vp-AI
CeA
BJ2569a1
GI
YC1060a1
5:L3a hourglass
da-AI
YC0703a2
DI
YC0514a1
da-AI
YC0447a1
da-AI
YCNE0031a1
DI
BJ2433a1
vp-AI
YC1770a1
da-AI
BJ2222a1

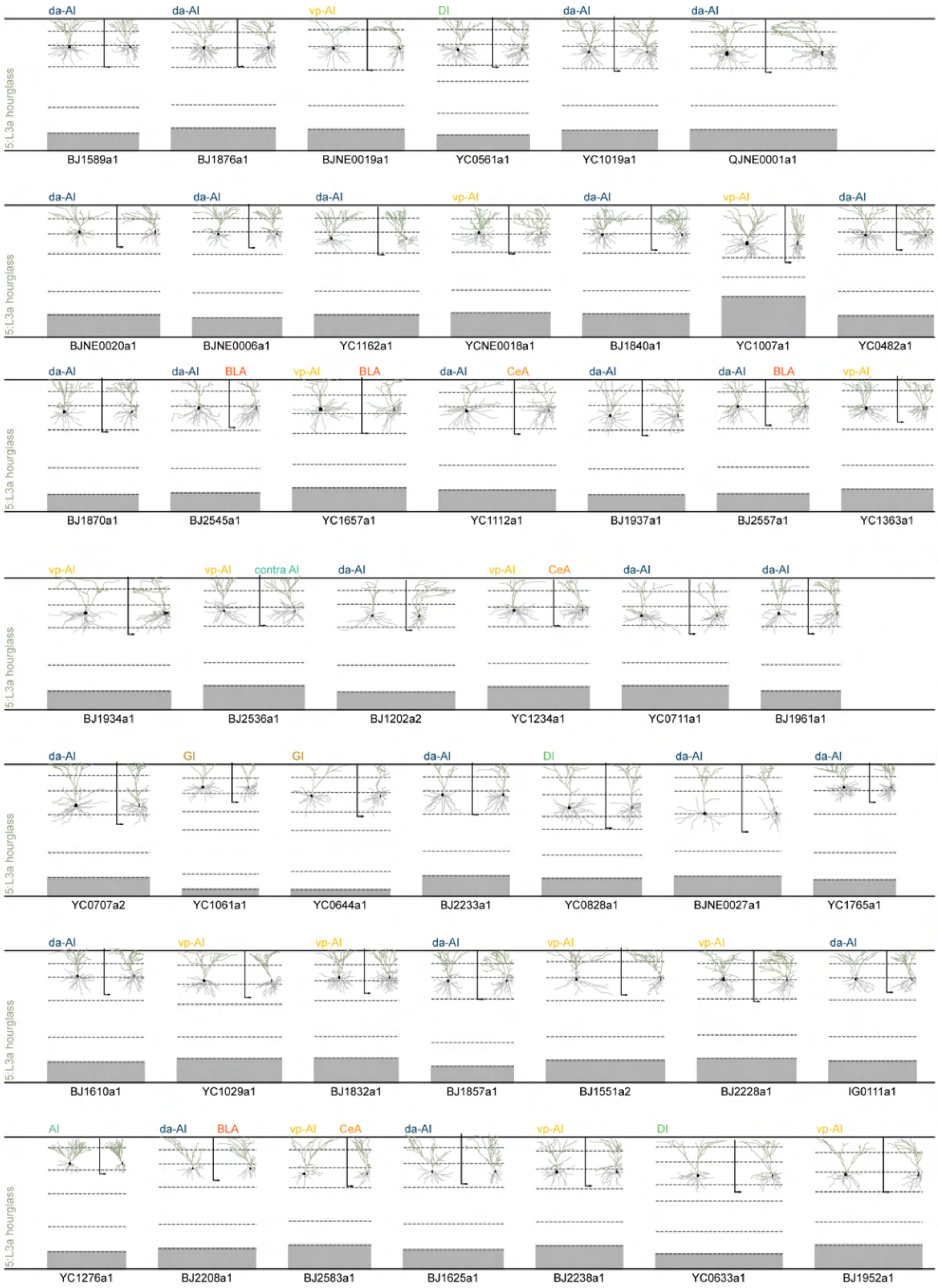
5:L3a hourglass
da-AI
da-AI
vp-AI
DI
da-AI
da-AI
BJ1589a1
BJ1876a1
BJNE0019a1
YC0561a1
YC1019a1
QJNE0001a1
da-AI
da-AI
da-AI
vp-AI
da-AI
vp-AI
da-AI
BJNE0020a1
BJNE0006a1
YC1162a1
YCNE0018a1
BJ1840a1
YC1007a1
YC0482a1
da-AI
da-AI BLA
vp-AI BLA
da-AI CeA
da-AI
da-AI BLA
vp-AI
BJ1870a1
BJ2545a1
YC1657a1
YC1112a1
BJ1937a1
BJ2557a1
YC1363a1
vp-AI
vp-AI contra AI
da-AI
vp-AI CeA
da-AI
da-AI
BJ1934a1
BJ2536a1
BJ1202a2
YC1234a1
YC0711a1
BJ1961a1
da-AI
GI
GI
da-AI
DI
da-AI
da-AI
YC0707a2
YC1061a1
YC0644a1
BJ2233a1
YC0828a1
BJNE0027a1
YC1765a1
da-AI
vp-AI
vp-AI
da-AI
vp-AI
vp-AI
da-AI
BJ1610a1
YC1029a1
BJ1832a1
BJ1857a1
BJ1551a2
BJ2228a1
IG0111a1
AI
da-AI BLA
vp-AI CeA
da-AI
vp-AI
DI
vp-AI
YC1276a1
BJ2208a1
BJ2583a1
BJ1625a1
BJ2238a1
YC0633a1
BJ1952a1

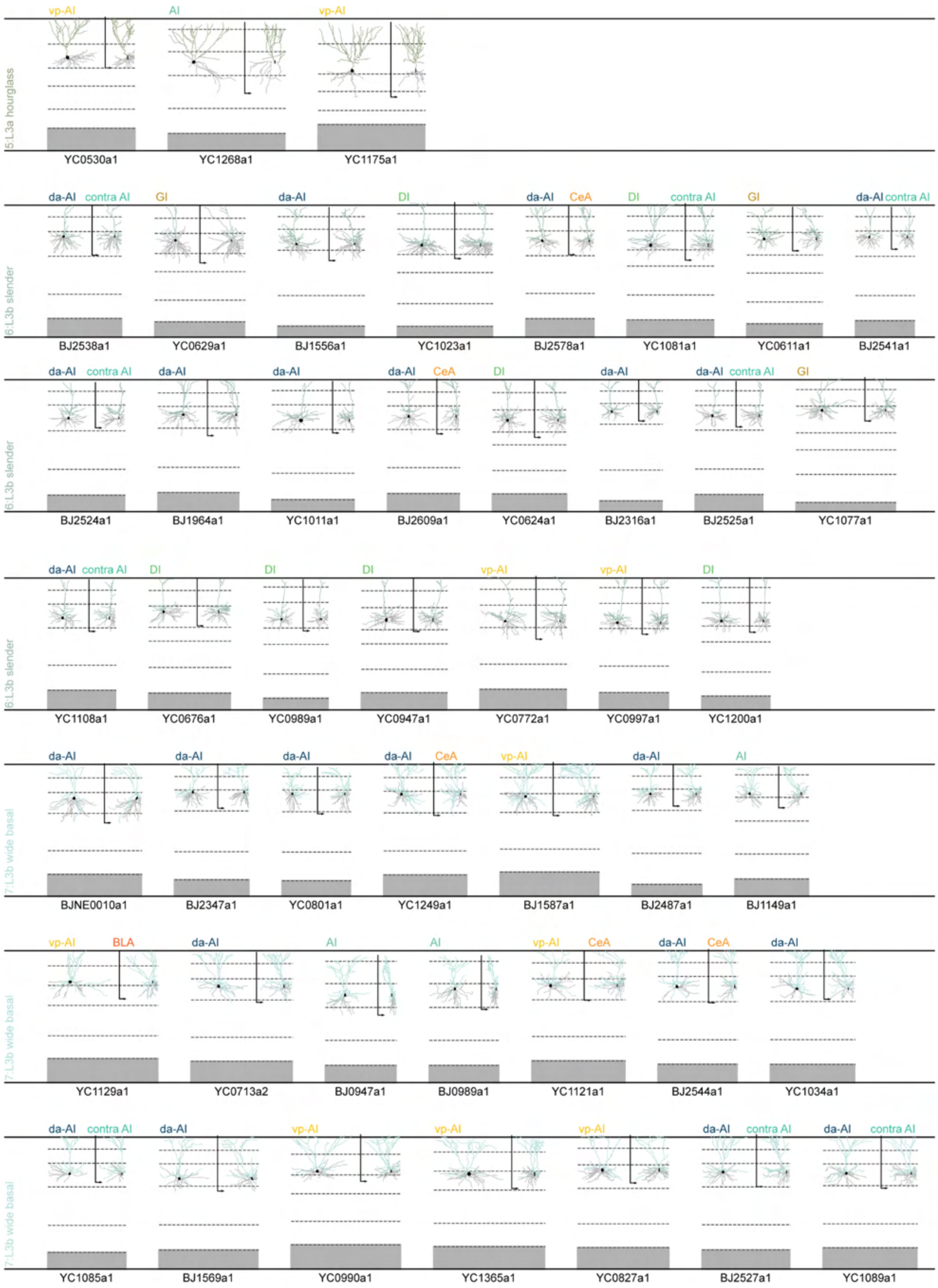
5:L3a hourglass
vp-AI
AI
vp-AI
YC0530a1
YC1268a1
YC1175a1
6:L3b slender
da-AI contra AI
GI
da-AI
DI
da-AI CeA
DI contra AI
GI
da-AI contra AI
BJ2538a1
YC0629a1
BJ1556a1
YC1023a1
BJ2578a1
YC1081a1
YC0611a1
BJ2541a1
6:L3b slender
da-AI contra AI
da-AI
da-AI
da-AI CeA
DI
da-AI
da-AI contra AI
GI
BJ2524a1
BJ1964a1
YC1011a1
BJ2609a1
YC0624a1
BJ2316a1
BJ2525a1
YC1077a1
6:L3b slender
da-AI contra AI
DI
DI
DI
vp-AI
vp-AI
DI
YC1108a1
YC0676a1
YC0989a1
YC0947a1
YC0772a1
YC0997a1
YC1200a1
7:L3b wide basal
da-AI
da-AI
da-AI
da-AI CeA
vp-AI
da-AI
AI
BJNE0010a1
BJ2347a1
YC0801a1
YC1249a1
BJ1587a1
BJ2487a1
BJ1149a1
7:L3b wide basal
vp-AI BLA
da-AI
AI
AI
vp-AI CeA
da-AI CeA
da-AI
YC1129a1
YC0713a2
BJ0947a1
BJ0989a1
YC1121a1
BJ2544a1
YC1034a1
7:L3b wide basal
da-AI contra AI
da-AI
vp-AI
vp-AI
vp-AI
da-AI contra AI
da-AI contra AI
YC1085a1
BJ1569a1
YC0990a1
YC1365a1
YC0827a1
BJ2527a1
YC1089a1

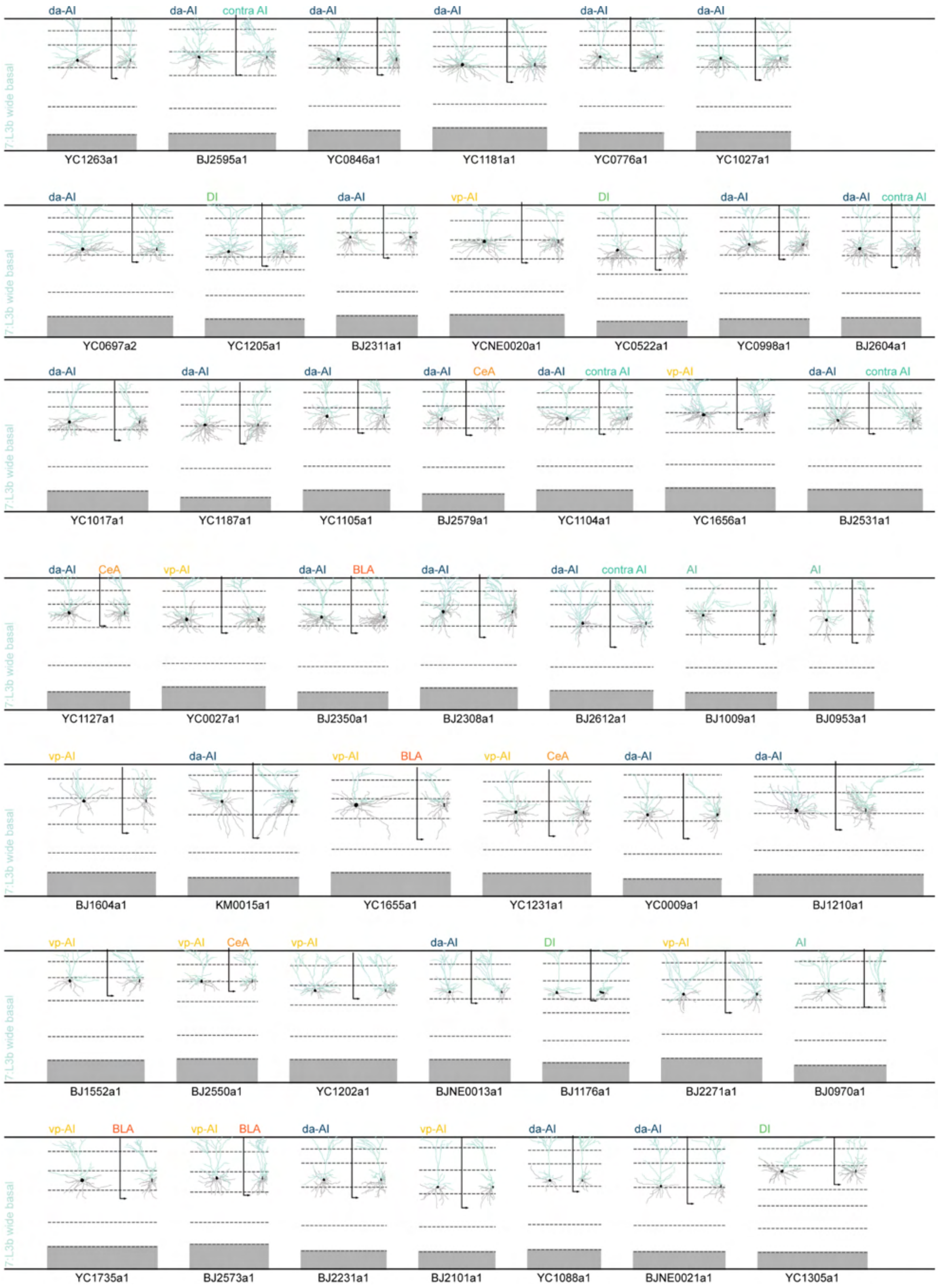
7:L3b wide basal
da-AI
YC1263a1
da-AI contra AI
BJ2595a1
da-AI
YC0846a1
da-AI
YC1181a1
da-AI
YC0776a1
da-AI
YC1027a1
7:L3b wide basal
da-AI
YC0697a2
DI
YC1205a1
da-AI
BJ2311a1
vp-AI
YCNE0020a1
DI
YC0522a1
da-AI
YC0998a1
da-AI contra AI
BJ2604a1
7:L3b wide basal
da-AI
YC1017a1
da-AI
YC1187a1
da-AI
YC1105a1
da-AI CeA
BJ2579a1
da-AI contra AI
YC1104a1
vp-AI
YC1656a1
da-AI contra AI
BJ2531a1
7:L3b wide basal
da-AI CeA
YC1127a1
vp-AI
YC0027a1
da-AI BLA
BJ2350a1
da-AI
BJ2308a1
da-AI contra AI
BJ2612a1
AI
BJ1009a1
AI
BJ0953a1
7:L3b wide basal
vp-AI
BJ1604a1
da-AI
KM0015a1
vp-AI BLA
YC1655a1
vp-AI CeA
YC1231a1
da-AI
YC0009a1
da-AI
BJ1210a1
7:L3b wide basal
vp-AI
BJ1552a1
vp-AI CeA
BJ2550a1
vp-AI
YC1202a1
da-AI
BJNE0013a1
DI
BJ1176a1
vp-AI
BJ2271a1
AI
BJ0970a1
7:L3b wide basal
vp-AI BLA
YC1735a1
vp-AI BLA
BJ2573a1
da-AI
BJ2231a1
vp-AI
BJ2101a1
da-AI
YC1088a1
da-AI
BJNE0021a1
DI
YC1305a1

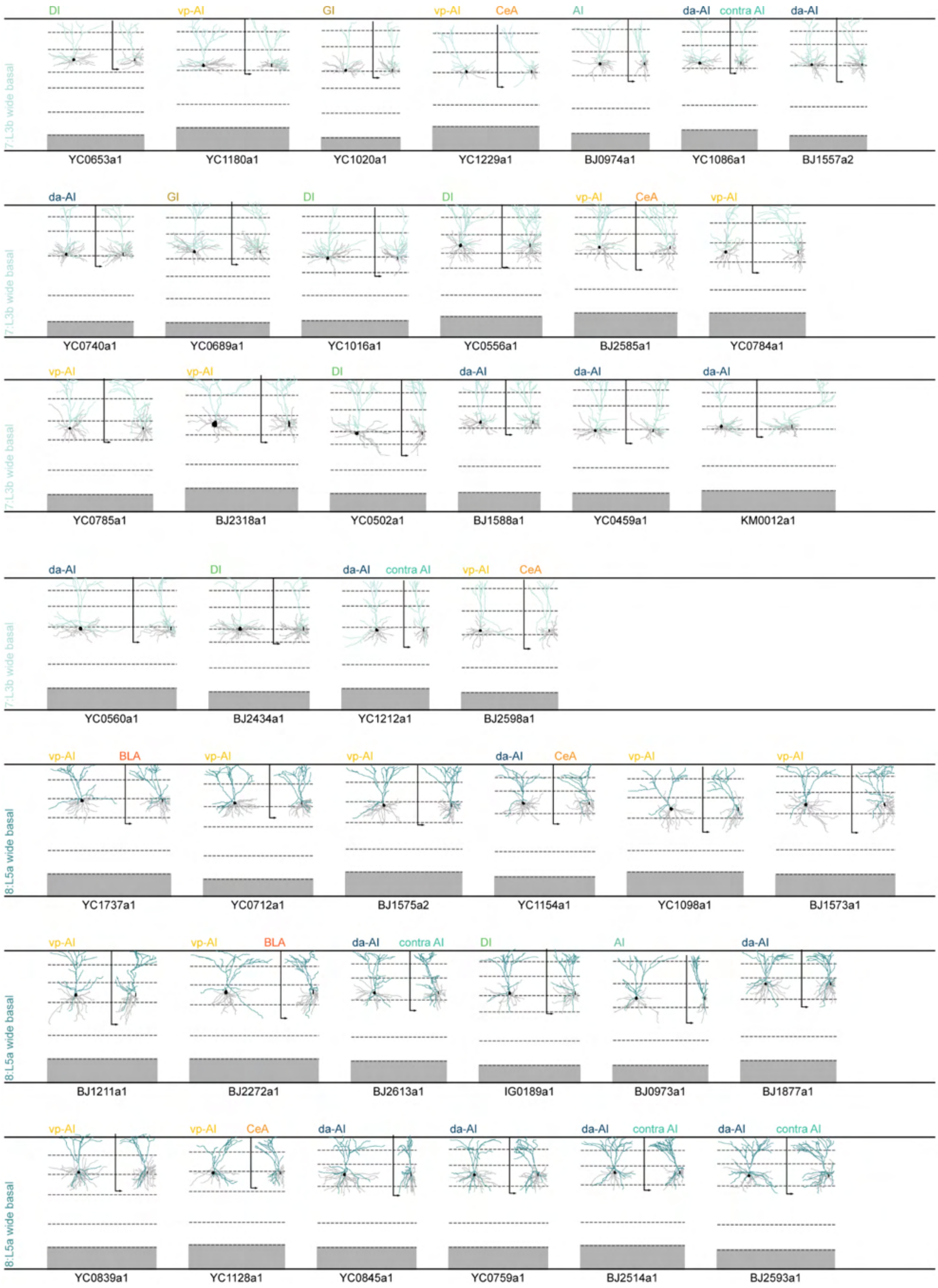

7:L3b wide basal
DI
YC0653a1
vp-AI
YC1180a1
GI
YC1020a1
vp-AI CeA
YC1229a1
AI
BJ0974a1
da-AI contra AI
YC1086a1
da-AI
BJ1557a2
7:L3b wide basal
da-AI
YC0740a1
GI
YC0689a1
DI
YC1016a1
DI
YC0556a1
vp-AI CeA
BJ2585a1
vp-AI
YC0784a1
7:L3b wide basal
vp-AI
YC0785a1
vp-AI
BJ2318a1
DI
YC0502a1
da-AI
BJ1588a1
da-AI
YC0459a1
da-AI
KM0012a1
7:L3b wide basal
da-AI
YC0560a1
DI
BJ2434a1
da-AI contra AI
YC1212a1
vp-AI CeA
BJ2598a1
8:L5a wide basal
vp-AI BLA
YC1737a1
vp-AI
YC0712a1
vp-AI
BJ1575a2
da-AI CeA
YC1154a1
vp-AI
YC1098a1
vp-AI
BJ1573a1
8:L5a wide basal
vp-AI
BJ1211a1
vp-AI BLA
BJ2272a1
da-AI contra AI
BJ2613a1
DI
IG0189a1
AI
BJ0973a1
da-AI
BJ1877a1
8:L5a wide basal
vp-AI
YC0839a1
vp-AI CeA
YC1128a1
da-AI
YC0845a1
da-AI
YC0759a1
da-AI contra AI
BJ2514a1
da-AI contra AI
BJ2593a1

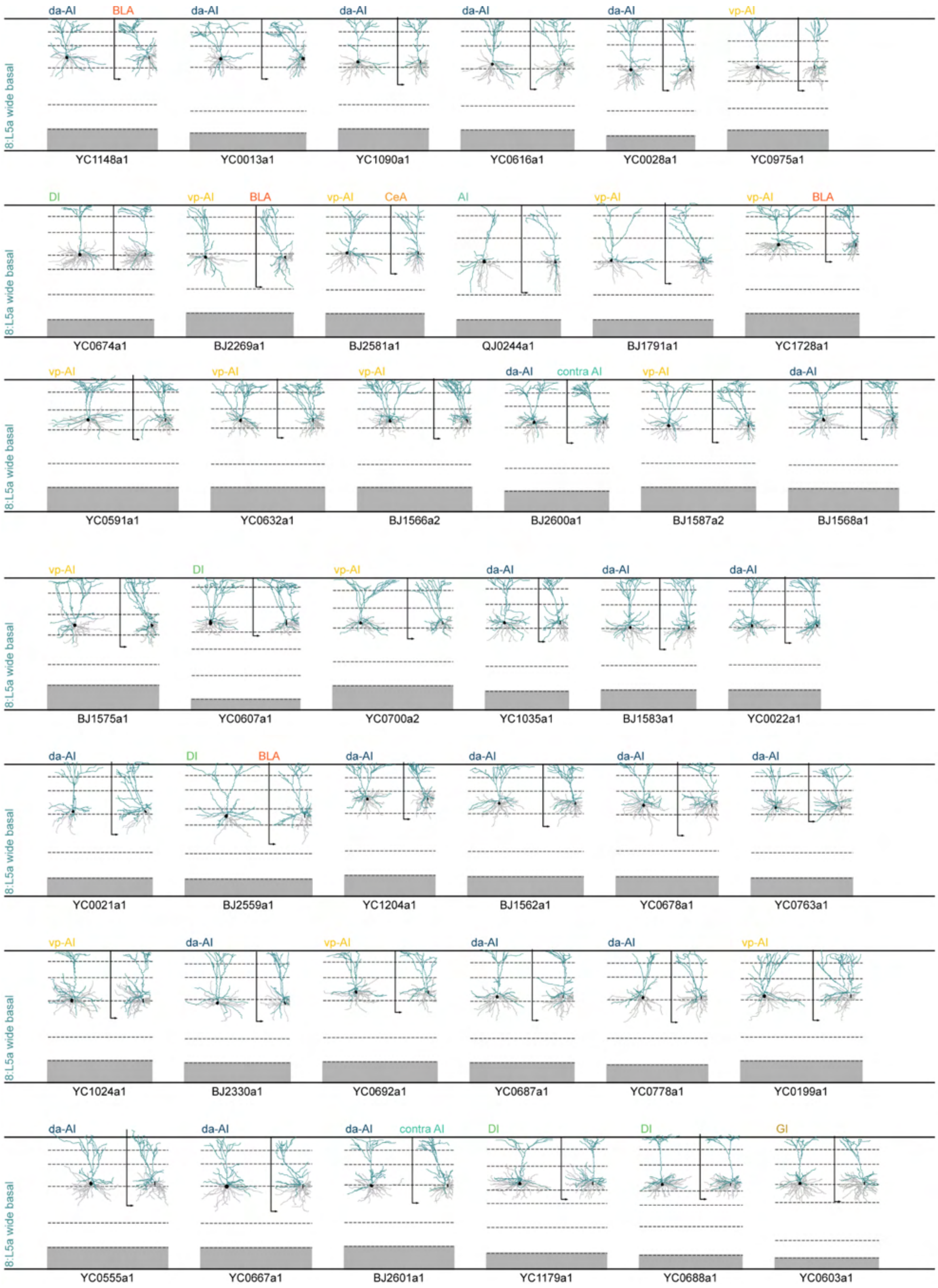
da-AI
BLA
da-AI
da-AI
da-AI
da-AI
vp-AI
8:L5a wide basal
YC1148a1
YC0013a1
YC1090a1
YC0616a1
YC0028a1
YC0975a1
DI
vp-AI
BLA
vp-AI
CeA
AI
vp-AI
vp-AI
BLA
8:L5a wide basal
YC0674a1
BJ2269a1
BJ2581a1
QJ0244a1
BJ1791a1
YC1728a1
vp-AI
vp-AI
vp-AI
da-AI
contra AI
vp-AI
da-AI
8:L5a wide basal
YC0591a1
YC0632a1
BJ1566a2
BJ2600a1
BJ1587a2
BJ1568a1
vp-AI
DI
vp-AI
da-AI
da-AI
da-AI
8:L5a wide basal
BJ1575a1
YC0607a1
YC0700a2
YC1035a1
BJ1583a1
YC0022a1
da-AI
DI
BLA
da-AI
da-AI
da-AI
da-AI
8:L5a wide basal
YC0021a1
BJ2559a1
YC1204a1
BJ1562a1
YC0678a1
YC0763a1
vp-AI
da-AI
vp-AI
da-AI
da-AI
vp-AI
8:L5a wide basal
YC1024a1
BJ2330a1
YC0692a1
YC0687a1
YC0778a1
YC0199a1
da-AI
da-AI
da-AI
contra AI
DI
DI
GI
8:L5a wide basal
YC0555a1
YC0667a1
BJ2601a1
YC1179a1
YC0688a1
YC0603a1

1526

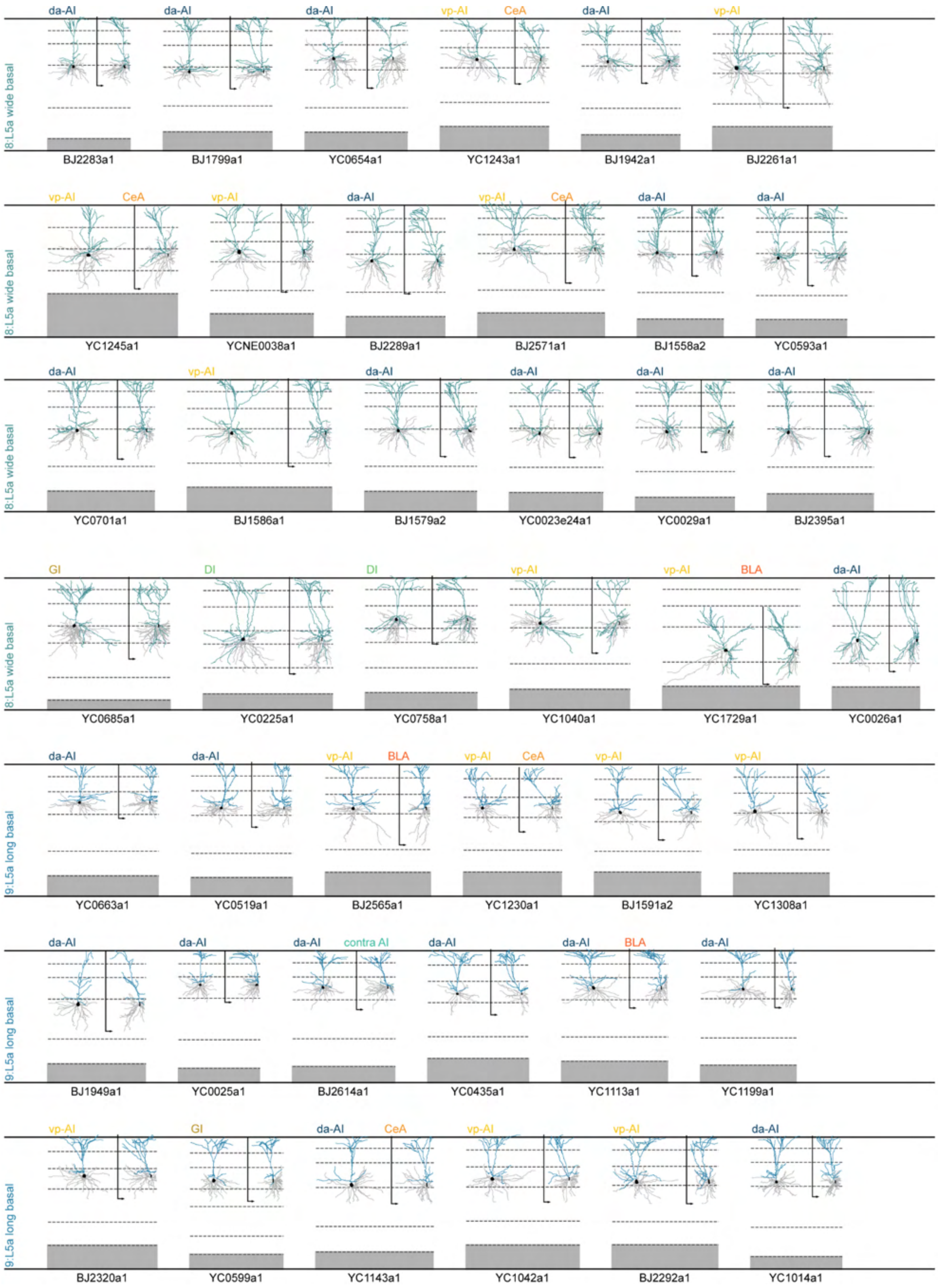
8:L5a wide basal
da-AI
BJ2283a1
da-AI
BJ1799a1
da-AI
YC0654a1
vp-AI
CeA
YC1243a1
da-AI
BJ1942a1
vp-AI
BJ2261a1
vp-AI
CeA
YC1245a1
vp-AI
YCNE0038a1
da-AI
BJ2289a1
vp-AI
CeA
BJ2571a1
da-AI
BJ1558a2
da-AI
YC0593a1
da-AI
YC0701a1
vp-AI
BJ1586a1
da-AI
BJ1579a2
da-AI
YC0023e24a1
da-AI
YC0029a1
da-AI
BJ2395a1
GI
YC0685a1
DI
YC0225a1
DI
YC0758a1
vp-AI
YC1040a1
vp-AI
BLA
YC1729a1
da-AI
YC0026a1
9:L5a long basal
da-AI
YC0663a1
da-AI
YC0519a1
vp-AI
BLA
BJ2565a1
vp-AI
CeA
YC1230a1
vp-AI
BJ1591a2
vp-AI
YC1308a1
da-AI
BJ1949a1
da-AI
YC0025a1
da-AI
contra AI
BJ2614a1
da-AI
YC0435a1
da-AI
BLA
YC1113a1
da-AI
YC1199a1
vp-AI
BJ2320a1
GI
YC0599a1
da-AI
CeA
YC1143a1
vp-AI
YC1042a1
vp-AI
BJ2292a1
da-AI
YC1014a1

1527

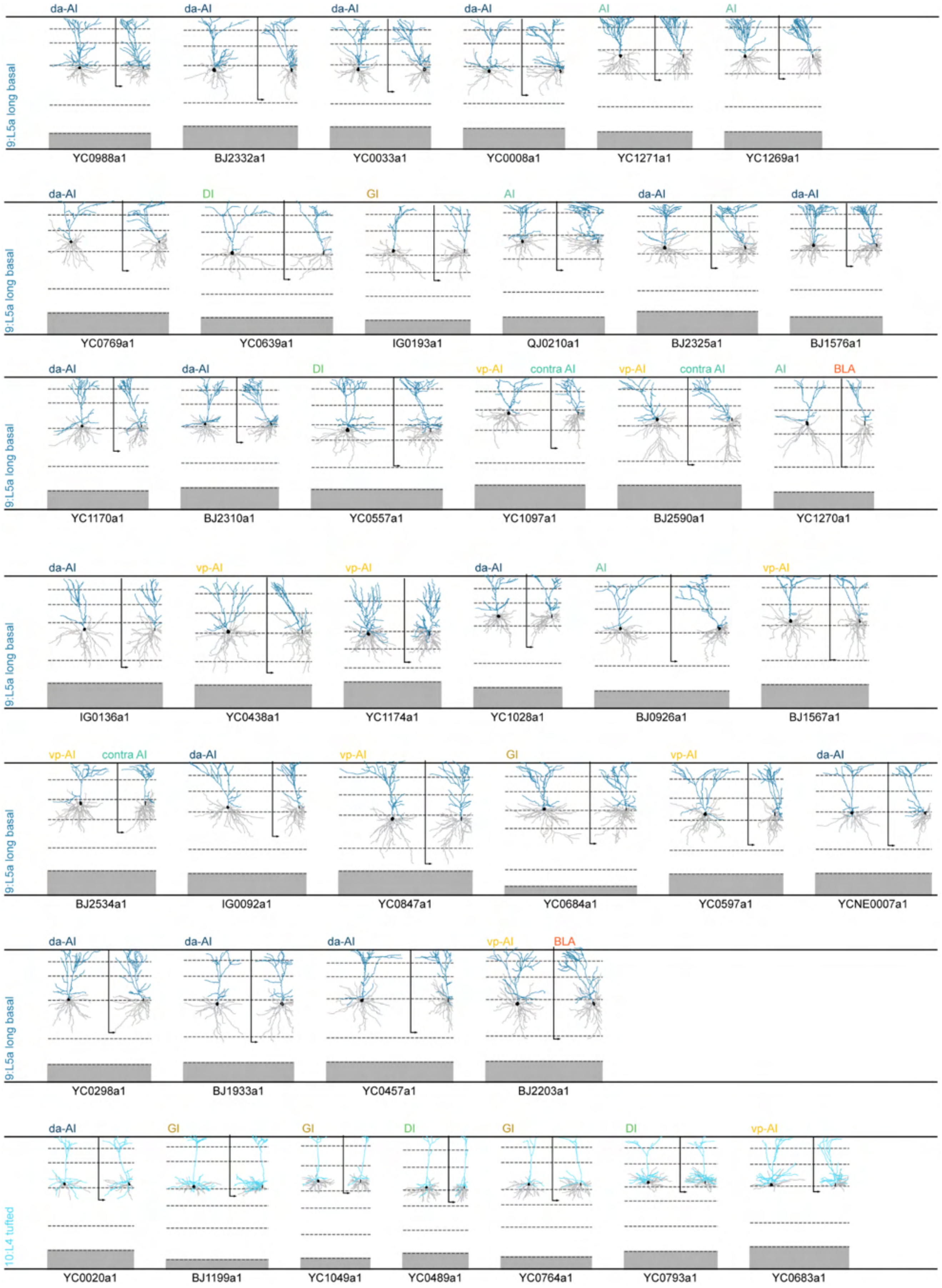
9:L5a long basal
da-AI
da-AI
da-AI
da-AI
AI
AI
YC0988a1
BJ2332a1
YC0033a1
YC0008a1
YC1271a1
YC1269a1
9:L5a long basal
da-AI
DI
GI
AI
da-AI
da-AI
YC0769a1
YC0639a1
IG0193a1
QJ0210a1
BJ2325a1
BJ1576a1
9:L5a long basal
da-AI
da-AI
DI
vp-AI
contra AI
vp-AI
contra AI
AI
BLA
YC1170a1
BJ2310a1
YC0557a1
YC1097a1
BJ2590a1
YC1270a1
9:L5a long basal
da-AI
vp-AI
vp-AI
da-AI
AI
vp-AI
IG0136a1
YC0438a1
YC1174a1
YC1028a1
BJ0926a1
BJ1567a1
9:L5a long basal
vp-AI
contra AI
da-AI
vp-AI
GI
vp-AI
da-AI
BJ2534a1
IG0092a1
YC0847a1
YC0684a1
YC0597a1
YCNE0007a1
9:L5a long basal
da-AI
da-AI
da-AI
vp-AI
BLA
YC0298a1
BJ1933a1
YC0457a1
BJ2203a1
10:L4 tufted
da-AI
GI
GI
DI
GI
DI
vp-AI
YC0020a1
BJ1199a1
YC1049a1
YC0489a1
YC0764a1
YC0793a1
YC0683a1

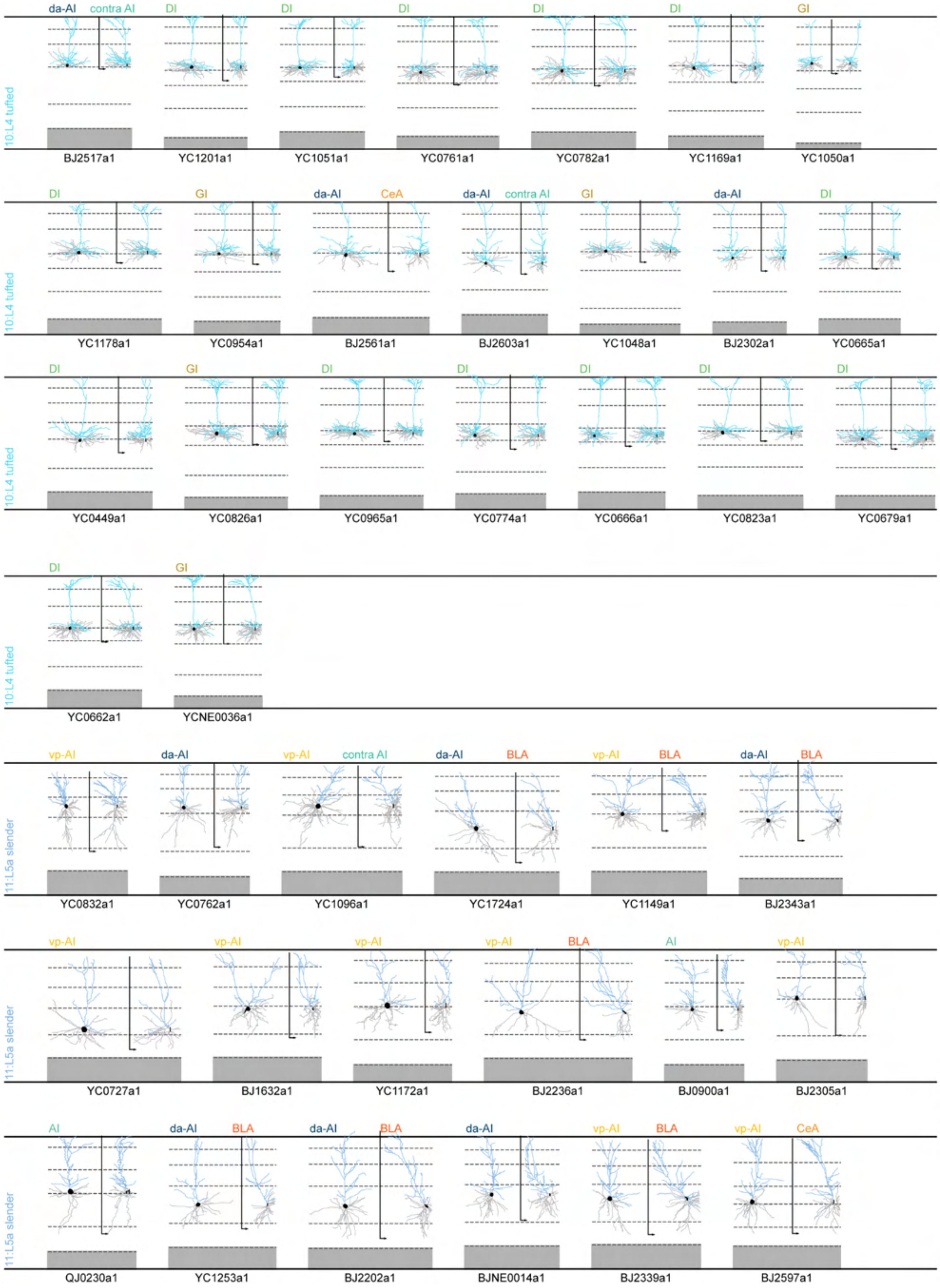

da-AI
contra AI
DI
DI
DI
DI
DI
GI
10:L4 tufted
BJ2517a1
YC1201a1
YC1051a1
YC0761a1
YC0782a1
YC1169a1
YC1050a1
DI
GI
da-AI
CeA
da-AI
contra AI
GI
da-AI
DI
10:L4 tufted
YC1178a1
YC0954a1
BJ2561a1
BJ2603a1
YC1048a1
BJ2302a1
YC0665a1
DI
GI
DI
DI
DI
DI
DI
10:L4 tufted
YC0449a1
YC0826a1
YC0965a1
YC0774a1
YC0666a1
YC0823a1
YC0679a1
DI
GI
10:L4 tufted
YC0662a1
YCNE0036a1
vp-AI
da-AI
vp-AI
contra AI
da-AI
BLA
vp-AI
BLA
da-AI
BLA
11:L5a slender
YC0832a1
YC0762a1
YC1096a1
YC1724a1
YC1149a1
BJ2343a1
vp-AI
vp-AI
vp-AI
vp-AI
BLA
AI
vp-AI
11:L5a slender
YC0727a1
BJ1632a1
YC1172a1
BJ2236a1
BJ0900a1
BJ2305a1
AI
da-AI
BLA
da-AI
BLA
da-AI
vp-AI
BLA
vp-AI
CeA
11:L5a slender
QJ0230a1
YC1253a1
BJ2202a1
BJNE0014a1
BJ2339a1
BJ2597a1

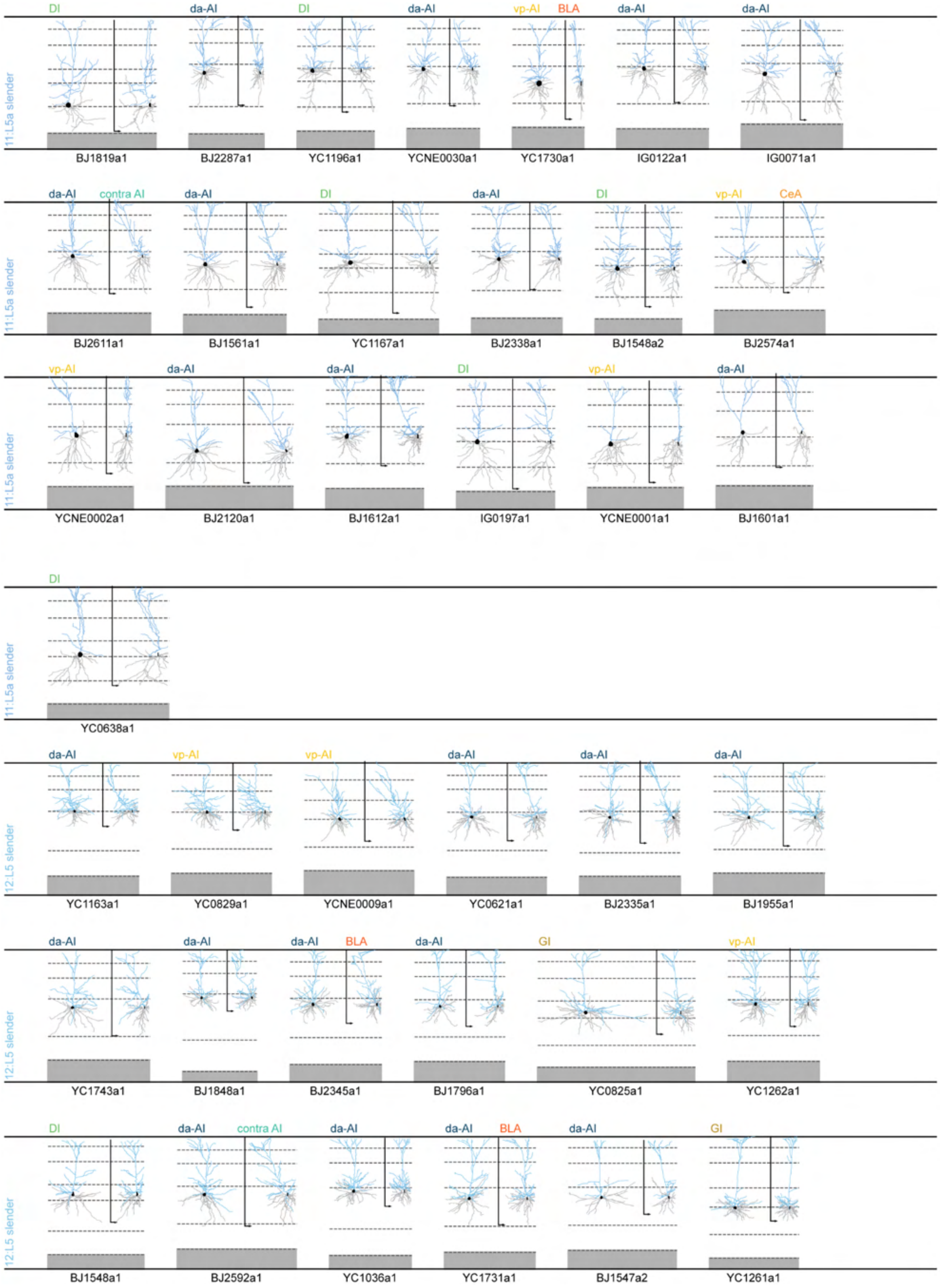
DI
da-AI
DI
da-AI
vp-AI
BLA
da-AI
da-AI
11:L5a slender
BJ1819a1
BJ2287a1
YC1196a1
YCNE0030a1
YC1730a1
IG0122a1
IG0071a1
da-AI
contra AI
da-AI
DI
da-AI
DI
vp-AI
CeA
11:L5a slender
BJ2611a1
BJ1561a1
YC1167a1
BJ2338a1
BJ1548a2
BJ2574a1
vp-AI
da-AI
da-AI
DI
vp-AI
da-AI
11:L5a slender
YCNE0002a1
BJ2120a1
BJ1612a1
IG0197a1
YCNE0001a1
BJ1601a1
DI
11:L5a slender
YC0638a1
da-AI
vp-AI
vp-AI
da-AI
da-AI
da-AI
12:L5 slender
YC1163a1
YC0829a1
YCNE0009a1
YC0621a1
BJ2335a1
BJ1955a1
da-AI
da-AI
da-AI
BLA
da-AI
GI
vp-AI
12:L5 slender
YC1743a1
BJ1848a1
BJ2345a1
BJ1796a1
YC0825a1
YC1262a1
DI
da-AI
contra AI
da-AI
da-AI
BLA
da-AI
GI
12:L5 slender
BJ1548a1
BJ2592a1
YC1036a1
YC1731a1
BJ1547a2
YC1261a1

1531 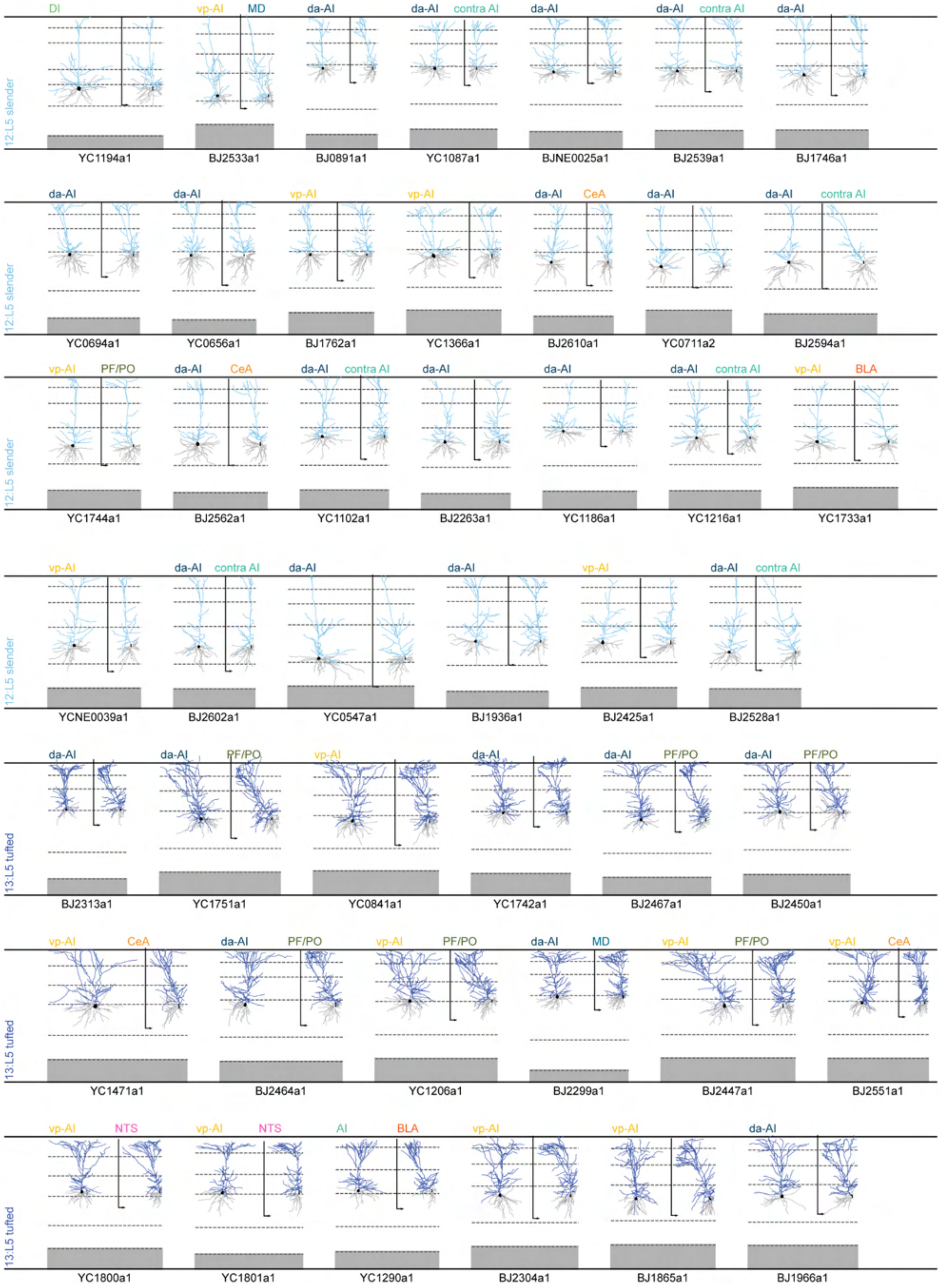
12:L5 slender
DI
YC1194a1
vp-AI
MD
BJ2533a1
da-AI
BJ0891a1
da-AI
contra AI
YC1087a1
da-AI
BJNE0025a1
da-AI
contra AI
BJ2539a1
da-AI
BJ1746a1
da-AI
YC0694a1
da-AI
YC0656a1
vp-AI
BJ1762a1
vp-AI
YC1366a1
da-AI
CeA
BJ2610a1
da-AI
YC0711a2
da-AI
contra AI
BJ2594a1
vp-AI
PF/PO
YC1744a1
da-AI
CeA
BJ2562a1
da-AI
contra AI
YC1102a1
da-AI
BJ2263a1
da-AI
YC1186a1
da-AI
contra AI
YC1216a1
vp-AI
BLA
YC1733a1
vp-AI
YCNE0039a1
da-AI
contra AI
BJ2602a1
da-AI
YC0547a1
da-AI
BJ1936a1
vp-AI
BJ2425a1
da-AI
contra AI
BJ2528a1
13:L5 tufted
da-AI
BJ2313a1
da-AI
PF/PO
YC1751a1
vp-AI
YC0841a1
da-AI
YC1742a1
da-AI
PF/PO
BJ2467a1
da-AI
PF/PO
BJ2450a1
vp-AI
CeA
YC1471a1
da-AI
PF/PO
BJ2464a1
vp-AI
PF/PO
YC1206a1
da-AI
MD
BJ2299a1
vp-AI
PF/PO
BJ2447a1
vp-AI
CeA
BJ2551a1
vp-AI
NTS
YC1800a1
vp-AI
NTS
YC1801a1
AI
BLA
YC1290a1
vp-AI
BJ2304a1
vp-AI
BJ1865a1
da-AI
BJ1966a1

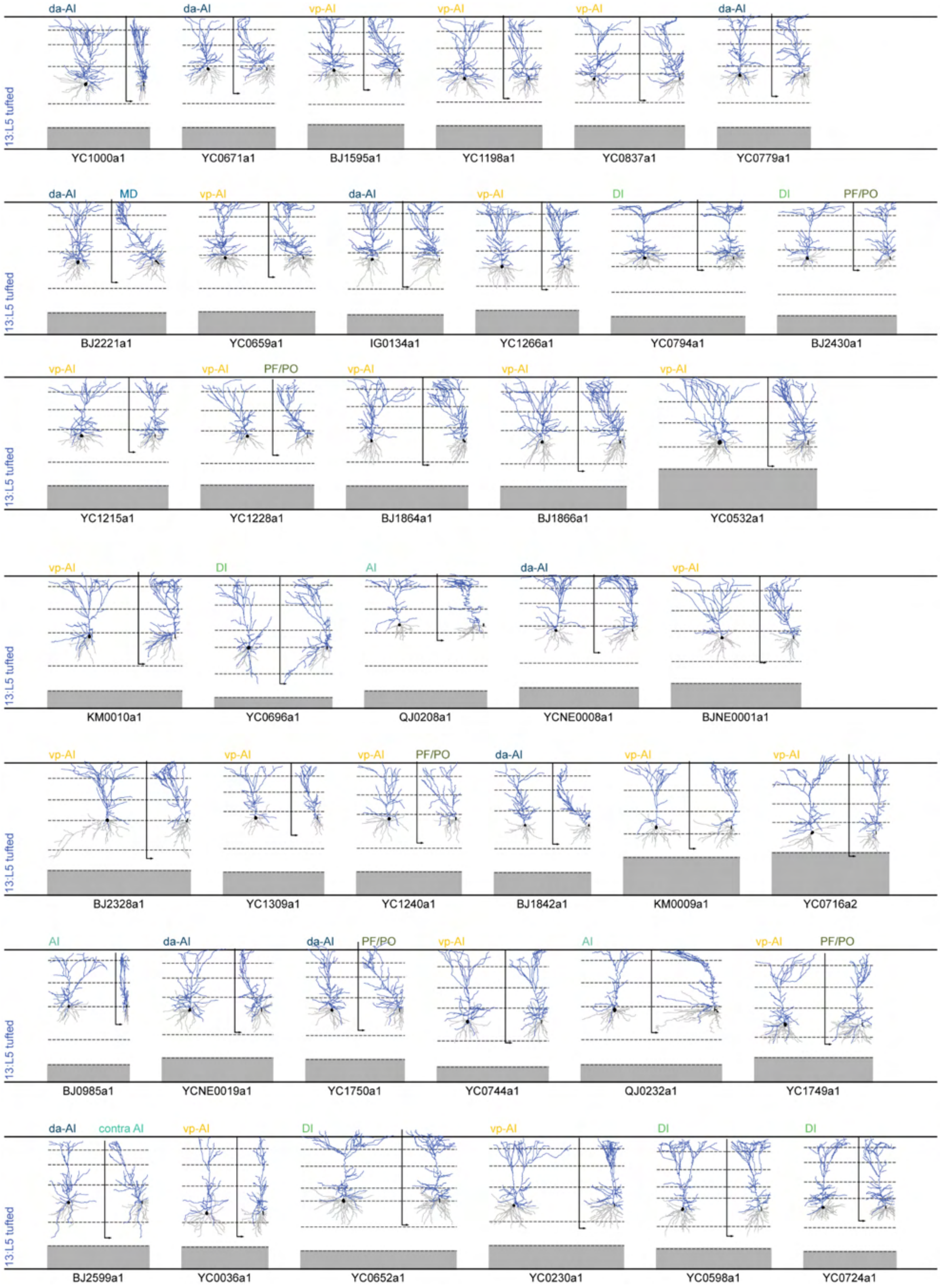
da-AI
da-AI
vp-AI
vp-AI
vp-AI
da-AI
13:L5 tufted
YC1000a1
YC0671a1
BJ1595a1
YC1198a1
YC0837a1
YC0779a1
da-AI
MD
vp-AI
da-AI
vp-AI
DI
DI
PF/PO
13:L5 tufted
BJ2221a1
YC0659a1
IG0134a1
YC1266a1
YC0794a1
BJ2430a1
vp-AI
vp-AI
PF/PO
vp-AI
vp-AI
vp-AI
13:L5 tufted
YC1215a1
YC1228a1
BJ1864a1
BJ1866a1
YC0532a1
vp-AI
DI
AI
da-AI
vp-AI
13:L5 tufted
KM0010a1
YC0696a1
QJ0208a1
YCNE0008a1
BJNE0001a1
vp-AI
vp-AI
vp-AI
PF/PO
da-AI
vp-AI
vp-AI
13:L5 tufted
BJ2328a1
YC1309a1
YC1240a1
BJ1842a1
KM0009a1
YC0716a2
AI
da-AI
da-AI
PF/PO
vp-AI
AI
vp-AI
PF/PO
13:L5 tufted
BJ0985a1
YCNE0019a1
YC1750a1
YC0744a1
QJ0232a1
YC1749a1
da-AI
contra AI
vp-AI
DI
vp-AI
DI
DI
13:L5 tufted
BJ2599a1
YC0036a1
YC0652a1
YC0230a1
YC0598a1
YC0724a1

1532

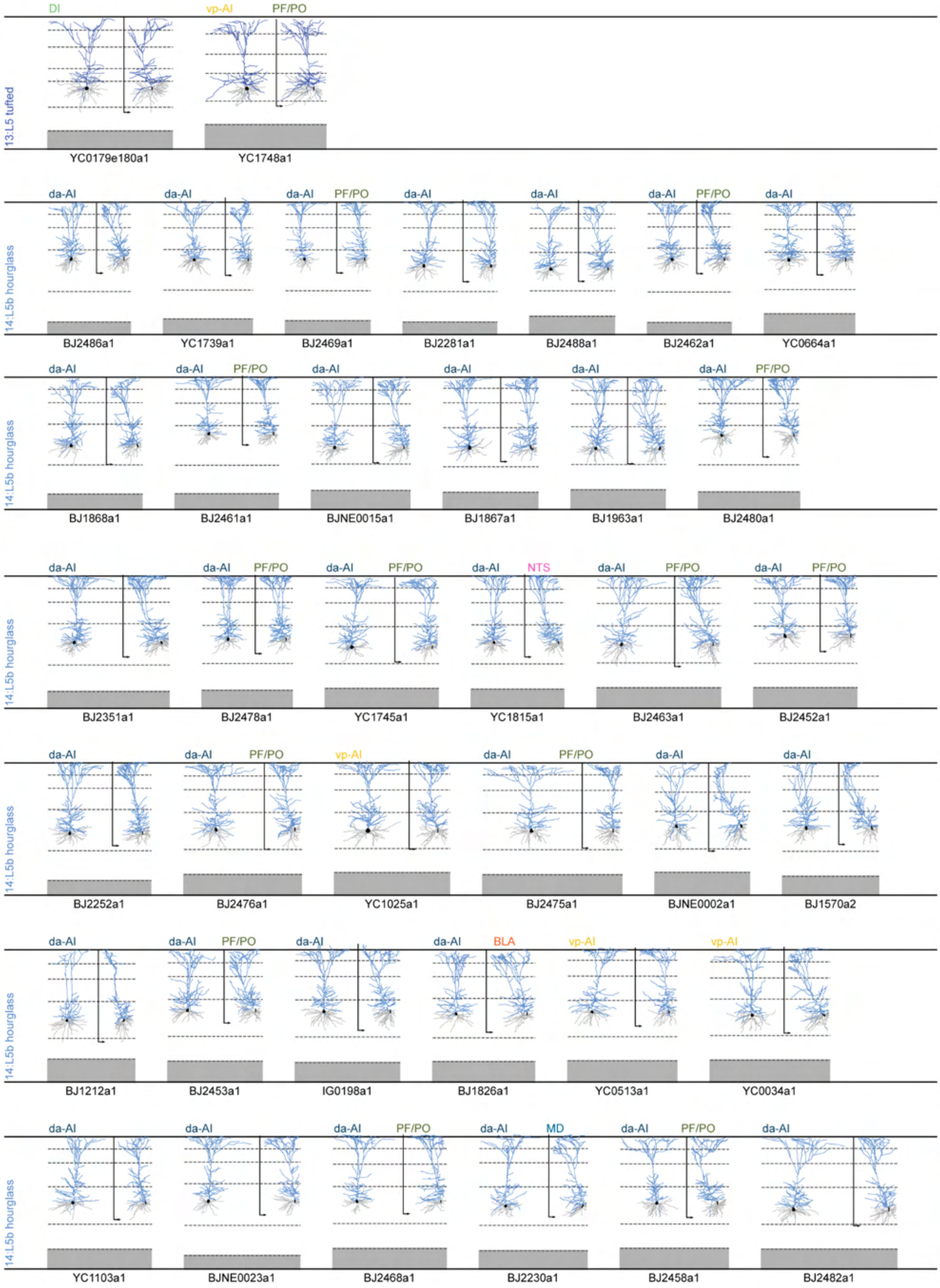
DI
vp-AI
PF/PO
13:L5 tufted
YC0179e180a1
YC1748a1
14:L5b hourglass
da-AI
PF/PO
BJ2486a1
YC1739a1
BJ2469a1
BJ2281a1
BJ2488a1
BJ2462a1
YC0664a1
BJ1868a1
BJ2461a1
BJNE0015a1
BJ1867a1
BJ1963a1
BJ2480a1
NTS
BJ2351a1
BJ2478a1
YC1745a1
YC1815a1
BJ2463a1
BJ2452a1
BJ2252a1
BJ2476a1
YC1025a1
BJ2475a1
BJNE0002a1
BJ1570a2
BLA
BJ1212a1
BJ2453a1
IG0198a1
BJ1826a1
YC0513a1
YC0034a1
MD
YC1103a1
BJNE0023a1
BJ2468a1
BJ2230a1
BJ2458a1
BJ2482a1

1533

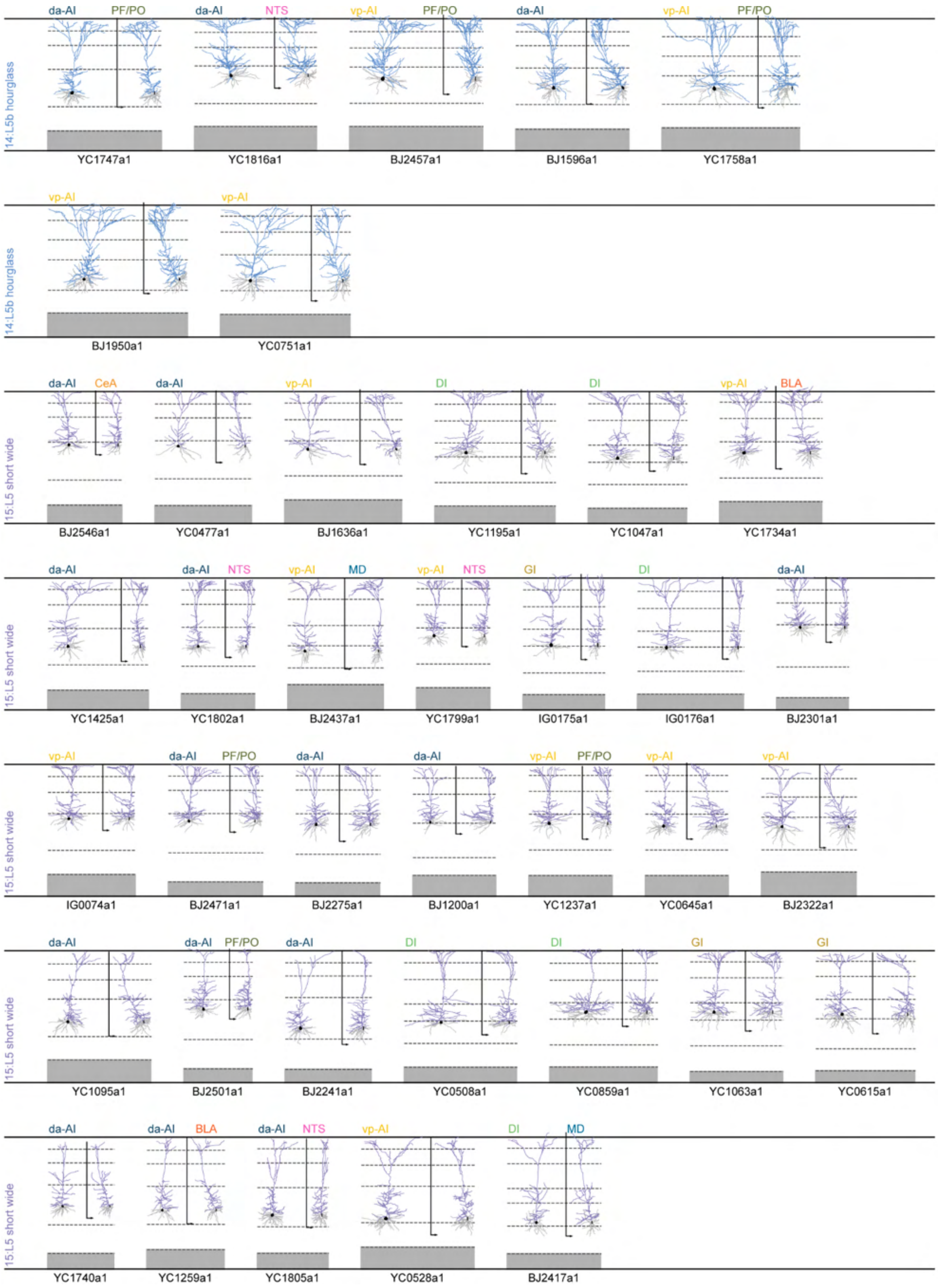
14:L5b hourglass
da-AI PF/PO
YC1747a1
da-AI NTS
YC1816a1
vp-AI PF/PO
BJ2457a1
da-AI
BJ1596a1
vp-AI PF/PO
YC1758a1
14:L5b hourglass
vp-AI
BJ1950a1
vp-AI
YC0751a1
15:L5 short wide
da-AI CeA
BJ2546a1
da-AI
YC0477a1
vp-AI
BJ1636a1
DI
YC1195a1
DI
YC1047a1
vp-AI BLA
YC1734a1
15:L5 short wide
da-AI
YC1425a1
da-AI NTS
YC1802a1
vp-AI MD
BJ2437a1
vp-AI NTS
YC1799a1
GI
IG0175a1
DI
IG0176a1
da-AI
BJ2301a1
15:L5 short wide
vp-AI
IG0074a1
da-AI PF/PO
BJ2471a1
da-AI
BJ2275a1
da-AI
BJ1200a1
vp-AI PF/PO
YC1237a1
vp-AI
YC0645a1
vp-AI
BJ2322a1
15:L5 short wide
da-AI
YC1095a1
da-AI PF/PO
BJ2501a1
da-AI
BJ2241a1
DI
YC0508a1
DI
YC0859a1
GI
YC1063a1
GI
YC0615a1
15:L5 short wide
da-AI
YC1740a1
da-AI BLA
YC1259a1
da-AI NTS
YC1805a1
vp-AI
YC0528a1
DI MD
BJ2417a1

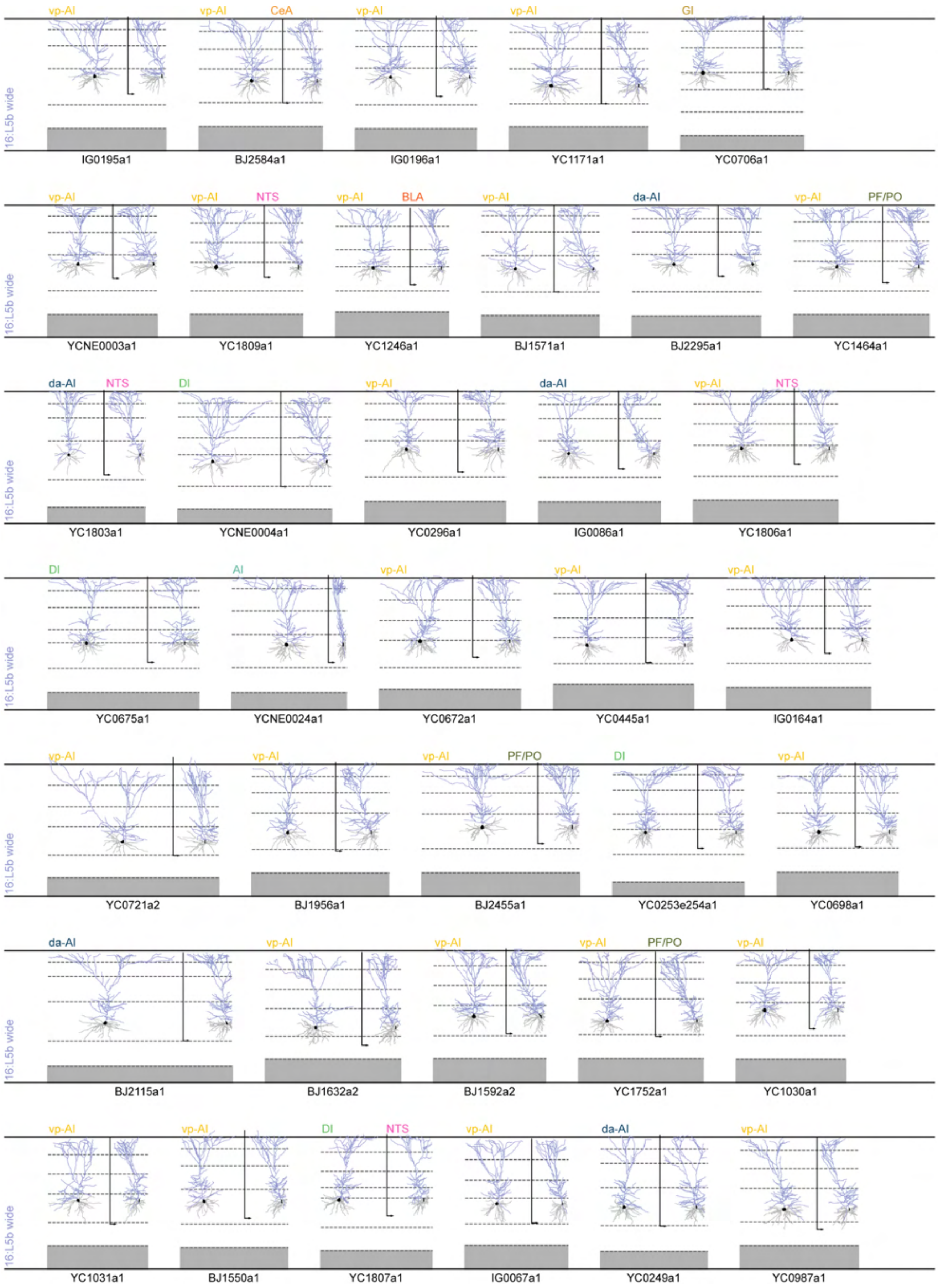
16:L5b wide
vp-AI
IG0195a1
vp-AI
CeA
BJ2584a1
vp-AI
IG0196a1
vp-AI
YC1171a1
GI
YC0706a1
16:L5b wide
vp-AI
YCNE0003a1
vp-AI
NTS
YC1809a1
vp-AI
BLA
YC1246a1
vp-AI
BJ1571a1
da-AI
BJ2295a1
vp-AI
PF/PO
YC1464a1
16:L5b wide
da-AI
NTS
YC1803a1
DI
YCNE0004a1
vp-AI
YC0296a1
da-AI
IG0086a1
vp-AI
NTS
YC1806a1
16:L5b wide
DI
YC0675a1
AI
YCNE0024a1
vp-AI
YC0672a1
vp-AI
YC0445a1
vp-AI
IG0164a1
16:L5b wide
vp-AI
YC0721a2
vp-AI
BJ1956a1
vp-AI
PF/PO
BJ2455a1
DI
YC0253e254a1
vp-AI
YC0698a1
16:L5b wide
da-AI
BJ2115a1
vp-AI
BJ1632a2
vp-AI
BJ1592a2
vp-AI
PF/PO
YC1752a1
vp-AI
YC1030a1
16:L5b wide
vp-AI
YC1031a1
vp-AI
BJ1550a1
DI
NTS
YC1807a1
vp-AI
IG0067a1
da-AI
YC0249a1
vp-AI
YC0987a1

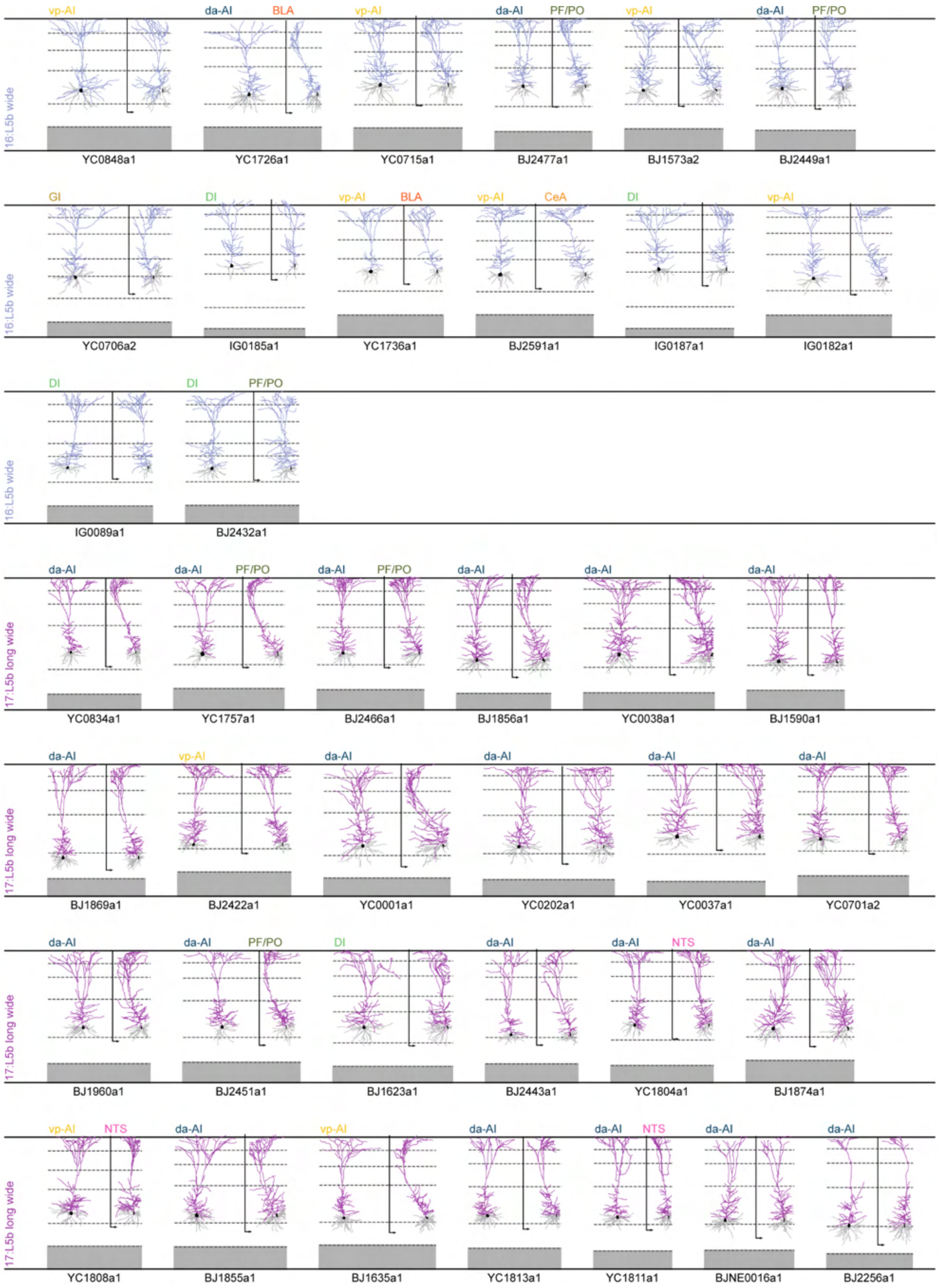
16:L5b wide
vp-AI
YC0848a1
da-AI
BLA
YC1726a1
vp-AI
YC0715a1
da-AI
PF/PO
BJ2477a1
vp-AI
BJ1573a2
da-AI
PF/PO
BJ2449a1
16:L5b wide
GI
YC0706a2
DI
IG0185a1
vp-AI
BLA
YC1736a1
vp-AI
CeA
BJ2591a1
DI
IG0187a1
vp-AI
IG0182a1
16:L5b wide
DI
IG0089a1
DI
PF/PO
BJ2432a1
17:L5b long wide
da-AI
YC0834a1
da-AI
PF/PO
YC1757a1
da-AI
PF/PO
BJ2466a1
da-AI
BJ1856a1
da-AI
YC0038a1
da-AI
BJ1590a1
17:L5b long wide
da-AI
BJ1869a1
vp-AI
BJ2422a1
da-AI
YC0001a1
da-AI
YC0202a1
da-AI
YC0037a1
da-AI
YC0701a2
17:L5b long wide
da-AI
BJ1960a1
da-AI
PF/PO
BJ2451a1
DI
BJ1623a1
da-AI
BJ2443a1
da-AI
NTS
YC1804a1
da-AI
BJ1874a1
17:L5b long wide
vp-AI
NTS
YC1808a1
da-AI
BJ1855a1
vp-AI
BJ1635a1
da-AI
YC1813a1
da-AI
NTS
YC1811a1
da-AI
BJNE0016a1
da-AI
BJ2256a1

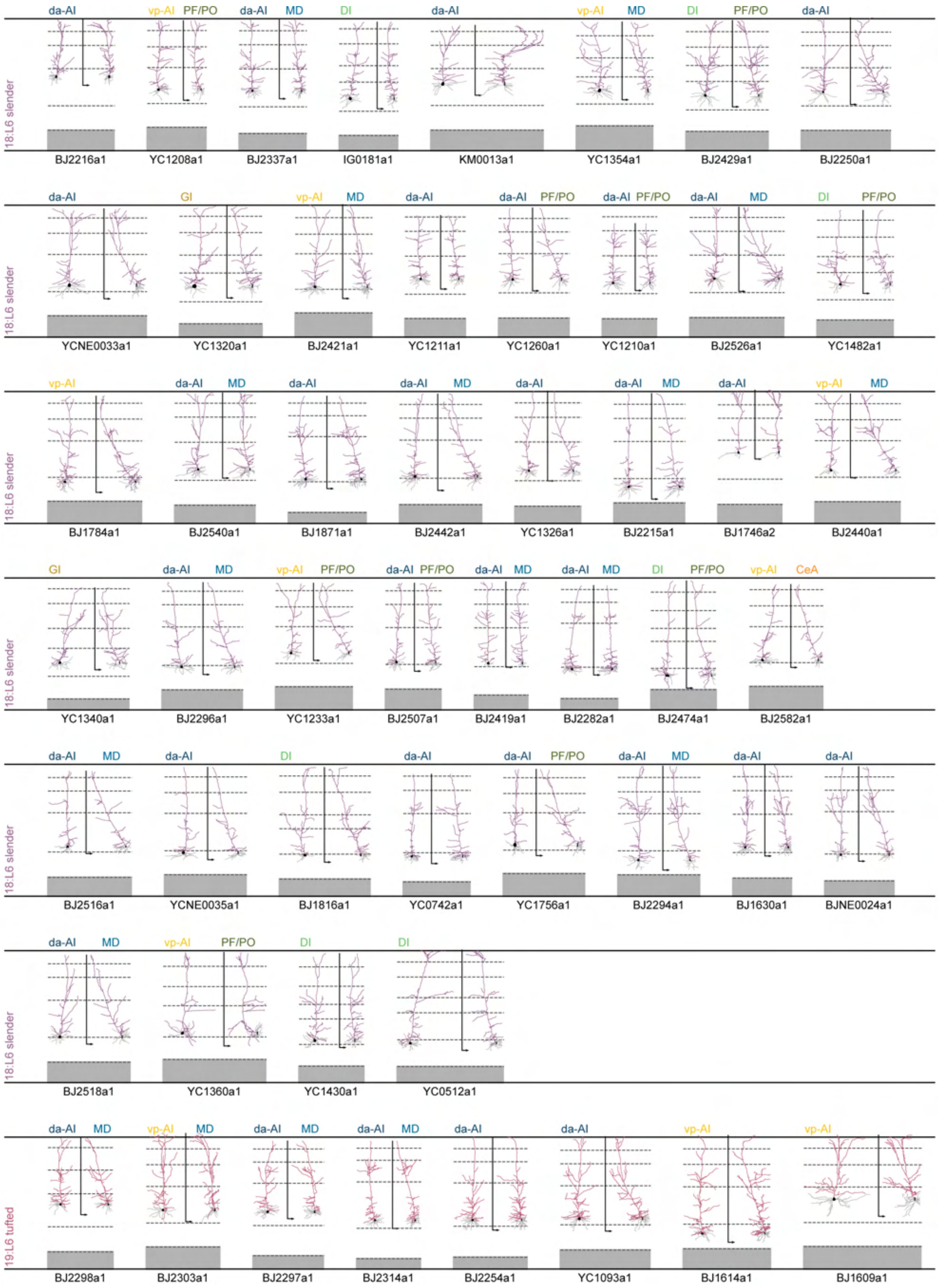
18:L6 slender
da-AI
BJ2216a1
vp-AI PF/PO
YC1208a1
da-AI MD
BJ2337a1
DI
IG0181a1
da-AI
KM0013a1
vp-AI MD
YC1354a1
DI PF/PO
BJ2429a1
da-AI
BJ2250a1
18:L6 slender
da-AI
YCNE0033a1
GI
YC1320a1
vp-AI MD
BJ2421a1
da-AI
YC1211a1
da-AI PF/PO
YC1260a1
da-AI PF/PO
YC1210a1
da-AI MD
BJ2526a1
DI PF/PO
YC1482a1
18:L6 slender
vp-AI
BJ1784a1
da-AI MD
BJ2540a1
da-AI
BJ1871a1
da-AI MD
BJ2442a1
da-AI
YC1326a1
da-AI MD
BJ2215a1
da-AI
BJ1746a2
vp-AI MD
BJ2440a1
18:L6 slender
GI
YC1340a1
da-AI MD
BJ2296a1
vp-AI PF/PO
YC1233a1
da-AI PF/PO
BJ2507a1
da-AI MD
BJ2419a1
da-AI MD
BJ2282a1
DI PF/PO
BJ2474a1
vp-AI CeA
BJ2582a1
18:L6 slender
da-AI MD
BJ2516a1
da-AI
YCNE0035a1
DI
BJ1816a1
da-AI
YC0742a1
da-AI PF/PO
YC1756a1
da-AI MD
BJ2294a1
da-AI
BJ1630a1
da-AI
BJNE0024a1
18:L6 slender
da-AI MD
BJ2518a1
vp-AI PF/PO
YC1360a1
DI
YC1430a1
DI
YC0512a1
19:L6 tufted
da-AI MD
BJ2298a1
vp-AI MD
BJ2303a1
da-AI MD
BJ2297a1
da-AI MD
BJ2314a1
da-AI
BJ2254a1
da-AI
YC1093a1
vp-AI
BJ1614a1
vp-AI
BJ1609a1

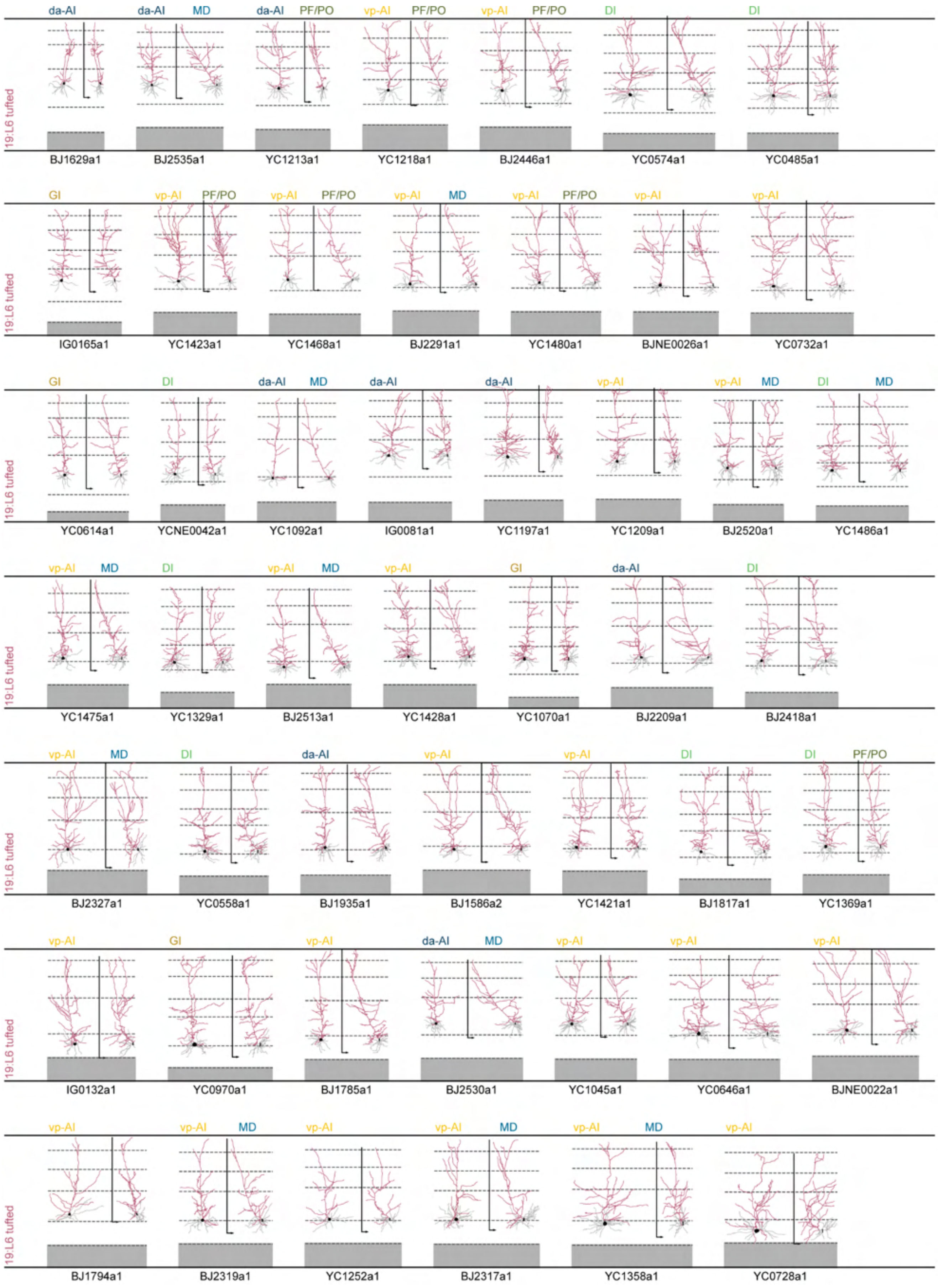
19:L6 tufted
da-AI
da-AI MD
da-AI PF/PO
vp-AI PF/PO
vp-AI PF/PO
DI
DI
BJ1629a1
BJ2535a1
YC1213a1
YC1218a1
BJ2446a1
YC0574a1
YC0485a1
GI
vp-AI PF/PO
vp-AI PF/PO
vp-AI MD
vp-AI PF/PO
vp-AI
vp-AI
IG0165a1
YC1423a1
YC1468a1
BJ2291a1
YC1480a1
BJNE0026a1
YC0732a1
GI
DI
da-AI MD
da-AI
da-AI
vp-AI
vp-AI MD
DI MD
YC0614a1
YCNE0042a1
YC1092a1
IG0081a1
YC1197a1
YC1209a1
BJ2520a1
YC1486a1
vp-AI MD
DI
vp-AI MD
vp-AI
GI
da-AI
DI
YC1475a1
YC1329a1
BJ2513a1
YC1428a1
YC1070a1
BJ2209a1
BJ2418a1
vp-AI MD
DI
da-AI
vp-AI
vp-AI
DI
DI PF/PO
BJ2327a1
YC0558a1
BJ1935a1
BJ1586a2
YC1421a1
BJ1817a1
YC1369a1
vp-AI
GI
vp-AI
da-AI MD
vp-AI
vp-AI
vp-AI
IG0132a1
YC0970a1
BJ1785a1
BJ2530a1
YC1045a1
YC0646a1
BJNE0022a1
vp-AI
vp-AI MD
vp-AI
vp-AI MD
vp-AI MD
vp-AI
BJ1794a1
BJ2319a1
YC1252a1
BJ2317a1
YC1358a1
YC0728a1

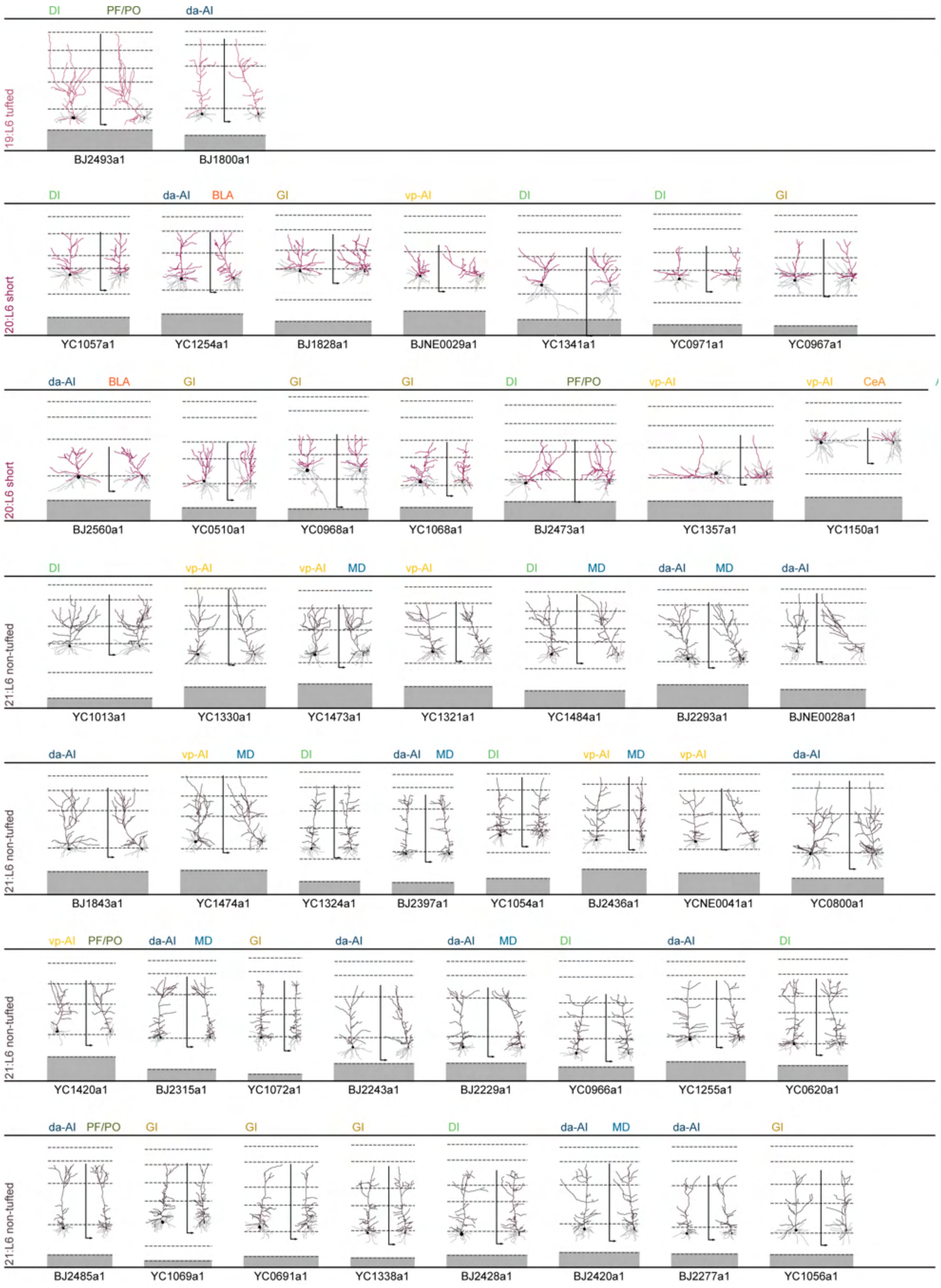

DI
PF/PO
da-AI
19:L6 tufted
BJ2493a1
BJ1800a1
DI
da-AI
BLA
GI
vp-AI
DI
DI
GI
20:L6 short
YC1057a1
YC1254a1
BJ1828a1
BJNE0029a1
YC1341a1
YC0971a1
YC0967a1
da-AI
BLA
GI
GI
GI
DI
PF/PO
vp-AI
vp-AI
CeA
20:L6 short
BJ2560a1
YC0510a1
YC0968a1
YC1068a1
BJ2473a1
YC1357a1
YC1150a1
DI
vp-AI
vp-AI
MD
vp-AI
DI
MD
da-AI
MD
da-AI
21:L6 non-tufted
YC1013a1
YC1330a1
YC1473a1
YC1321a1
YC1484a1
BJ2293a1
BJNE0028a1
da-AI
vp-AI
MD
DI
da-AI
MD
DI
vp-AI
MD
vp-AI
da-AI
21:L6 non-tufted
BJ1843a1
YC1474a1
YC1324a1
BJ2397a1
YC1054a1
BJ2436a1
YCNE0041a1
YC0800a1
vp-AI
PF/PO
da-AI
MD
GI
da-AI
da-AI
MD
DI
da-AI
DI
21:L6 non-tufted
YC1420a1
BJ2315a1
YC1072a1
BJ2243a1
BJ2229a1
YC0966a1
YC1255a1
YC0620a1
da-AI
PF/PO
GI
GI
GI
DI
da-AI
MD
da-AI
GI
21:L6 non-tufted
BJ2485a1
YC1069a1
YC0691a1
YC1338a1
BJ2428a1
BJ2420a1
BJ2277a1
YC1056a1

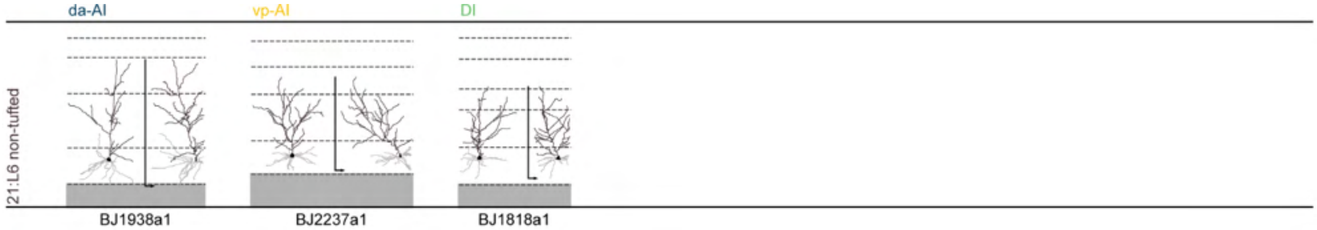

**Supplementary Fig. 2: Overview of 958 reconstructed dendritic morphologies.** Apical (colored) and basal (gray) dendrites shown in the *en face* (left) and side-view (right) planes for each reconstructed morphology that passed quality control (Methods). Dimensions are scaled to the local cortical thickness. Dashed lines demarcate layer boundaries between pia (top) and corpus callosum (bottom) for L1, L2, L3, L4[#], L5, L6, and claustrum (gray zone), as determined by mean local layer thickness following registration to the Nissl atlas (Extended Data Figs. 3 and 4). Corresponding insular subregion allocations (top-left) and projection targets (top-right, if determined) are shown as text. Individual morphologies are sorted within each M-type based on cluster analysis dendrogram. [#]L4 is only shown when located in the (dys)granular insula.

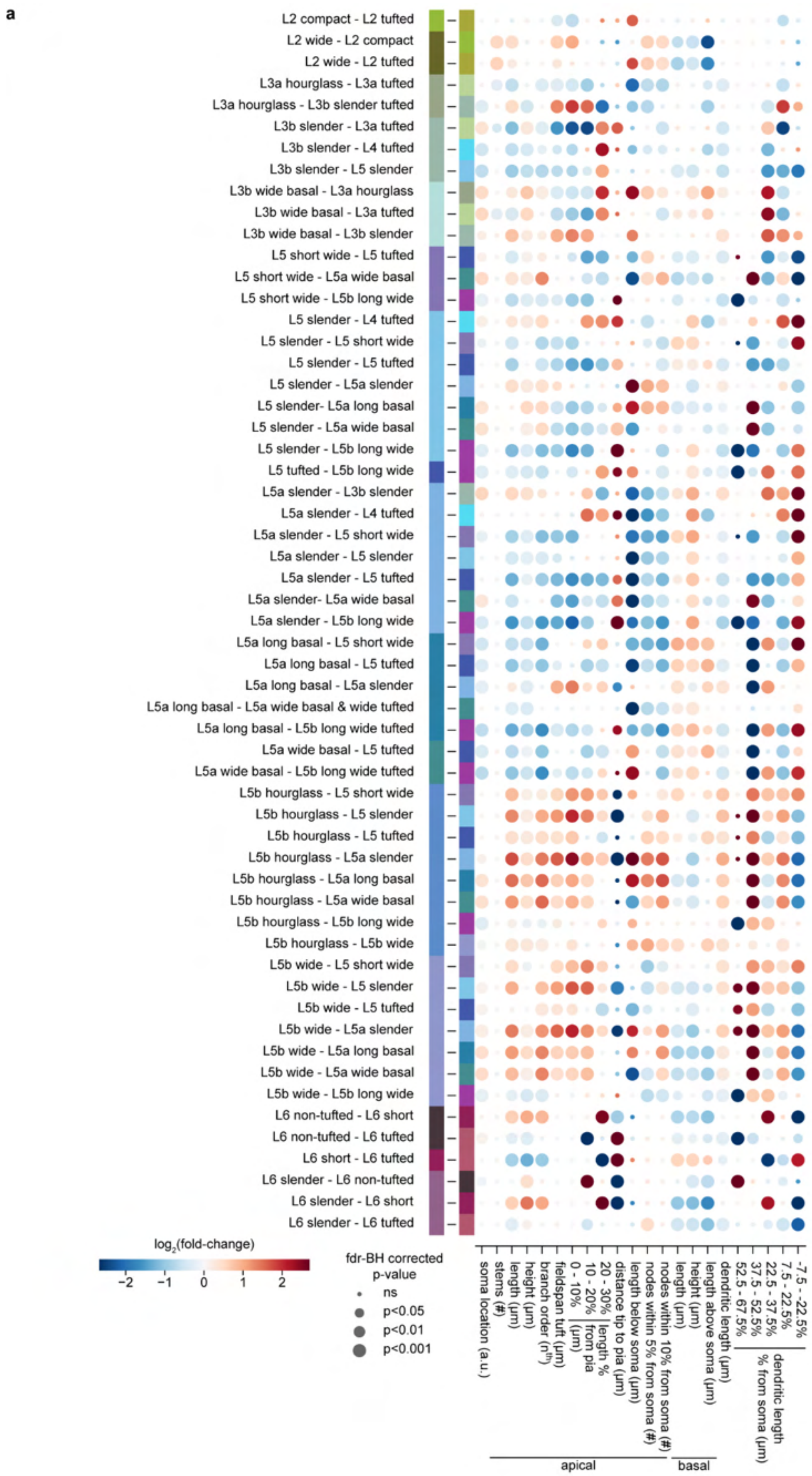
a
L2 compact - L2 tufted
L2 wide - L2 compact
L2 wide - L2 tufted
L3a hourglass - L3a tufted
L3a hourglass - L3b slender tufted
L3b slender - L3a tufted
L3b slender - L4 tufted
L3b slender - L5 slender
L3b wide basal - L3a hourglass
L3b wide basal - L3a tufted
L3b wide basal - L3b slender
L5 short wide - L5 tufted
L5 short wide - L5a wide basal
L5 short wide - L5b long wide
L5 slender - L4 tufted
L5 slender - L5 short wide
L5 slender - L5 tufted
L5 slender - L5a slender
L5 slender- L5a long basal
L5 slender - L5a wide basal
L5 slender - L5b long wide
L5 tufted - L5b long wide
L5a slender - L3b slender
L5a slender - L4 tufted
L5a slender - L5 short wide
L5a slender - L5 slender
L5a slender - L5 tufted
L5a slender- L5a wide basal
L5a slender - L5b long wide
L5a long basal - L5 short wide
L5a long basal - L5 tufted
L5a long basal - L5a slender
L5a long basal - L5a wide basal & wide tufted
L5a long basal - L5b long wide tufted
L5a wide basal - L5 tufted
L5a wide basal - L5b long wide tufted
L5b hourglass - L5 short wide
L5b hourglass - L5 slender
L5b hourglass - L5 tufted
L5b hourglass - L5a slender
L5b hourglass - L5a long basal
L5b hourglass - L5a wide basal
L5b hourglass - L5b long wide
L5b hourglass - L5b wide
L5b wide - L5 short wide
L5b wide - L5 slender
L5b wide - L5 tufted
L5b wide - L5a slender
L5b wide - L5a long basal
L5b wide - L5a wide basal
L5b wide - L5b long wide
L6 non-tufted - L6 short
L6 non-tufted - L6 tufted
L6 short - L6 tufted
L6 slender - L6 non-tufted
L6 slender - L6 short
L6 slender - L6 tufted
log2(fold-change)
−2
−1
0
1
2
fdr-BH corrected
p-value
ns
p<0.05
p<0.01
p<0.001
soma location (a.u.)
stems (#)
length (μm)
height (μm)
branch order (n^th)
fieldspan tuft (μm)
0 - 10%
10 - 20%
20 - 30%
from pia
length %
distance tip to pia (μm)
length below soma (μm)
nodes within 5% from soma (#)
nodes within 10% from soma (#)
length (μm)
height (μm)
length above soma (μm)
dendritic length (μm)
52.5 - 67.5%
37.5 - 52.5%
22.5 - 37.5%
7.5 - 22.5%
-7.5 - -22.5%
% from soma (μm)
dendritic length
apical
basal

1550

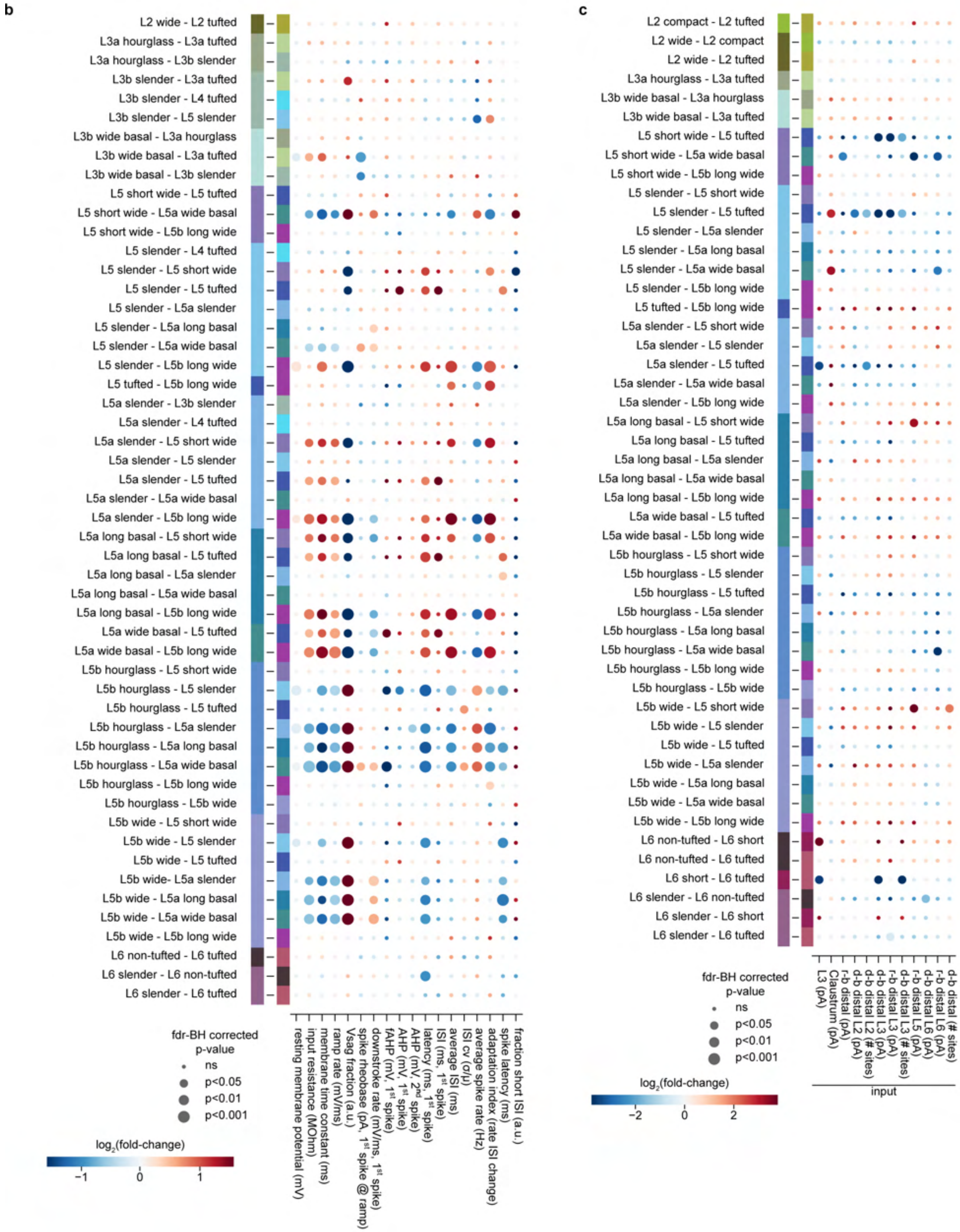
b
L2 wide - L2 tufted
L3a hourglass - L3a tufted
L3a hourglass - L3b slender
L3b slender - L3a tufted
L3b slender - L4 tufted
L3b slender - L5 slender
L3b wide basal - L3a hourglass
L3b wide basal - L3a tufted
L3b wide basal - L3b slender
L5 short wide - L5 tufted
L5 short wide - L5a wide basal
L5 short wide - L5b long wide
L5 slender - L4 tufted
L5 slender - L5 short wide
L5 slender - L5 tufted
L5 slender - L5a slender
L5 slender - L5a long basal
L5 slender - L5a wide basal
L5 slender - L5b long wide
L5 tufted - L5b long wide
L5a slender - L3b slender
L5a slender - L4 tufted
L5a slender - L5 short wide
L5a slender - L5 slender
L5a slender - L5 tufted
L5a slender - L5a wide basal
L5a slender - L5b long wide
L5a long basal - L5 short wide
L5a long basal - L5 tufted
L5a long basal - L5a slender
L5a long basal - L5a wide basal
L5a long basal - L5b long wide
L5a wide basal - L5 tufted
L5a wide basal - L5b long wide
L5b hourglass - L5 short wide
L5b hourglass - L5 slender
L5b hourglass - L5 tufted
L5b hourglass - L5a slender
L5b hourglass - L5a long basal
L5b hourglass - L5a wide basal
L5b hourglass - L5b long wide
L5b hourglass - L5b wide
L5b wide - L5 short wide
L5b wide - L5 slender
L5b wide - L5 tufted
L5b wide- L5a slender
L5b wide - L5a long basal
L5b wide - L5a wide basal
L5b wide - L5b long wide
L6 non-tufted - L6 tufted
L6 slender - L6 non-tufted
L6 slender - L6 tufted
fdr-BH corrected p-value
ns
p<0.05
p<0.01
p<0.001
log2(fold-change)
−1
0
1
resting membrane potential (mV)
input resistance (MOhm)
membrane time constant (ms)
ramp rate (mV/ms)
Vsag fraction (a.u.)
spike rheobase (pA, 1st spike @ ramp)
downstroke rate (mV/ms, 1st spike)
fAHP (mV, 1st spike)
AHP (mV, 1st spike)
AHP (mV, 2nd spike)
latency (ms, 1st spike)
ISI (ms, 1st spike)
average ISI (ms)
ISI cv (σ/μ)
average spike rate (Hz)
adaptation index (rate ISI change)
spike latency (ms)
fraction short ISI (a.u.)
c
L2 compact - L2 tufted
L2 wide - L2 compact
L2 wide - L2 tufted
L3a hourglass - L3a tufted
L3b wide basal - L3a hourglass
L3b wide basal - L3a tufted
L5 short wide - L5 tufted
L5 short wide - L5a wide basal
L5 short wide - L5b long wide
L5 slender - L5 short wide
L5 slender - L5 tufted
L5 slender - L5a slender
L5 slender - L5a long basal
L5 slender - L5a wide basal
L5 slender - L5b long wide
L5 tufted - L5b long wide
L5a slender - L5 short wide
L5a slender - L5 slender
L5a slender - L5 tufted
L5a slender - L5a wide basal
L5a slender - L5b long wide
L5a long basal - L5 short wide
L5a long basal - L5 tufted
L5a long basal - L5a slender
L5a long basal - L5a wide basal
L5a long basal - L5b long wide
L5a wide basal - L5 tufted
L5a wide basal - L5b long wide
L5b hourglass - L5 short wide
L5b hourglass - L5 slender
L5b hourglass - L5 tufted
L5b hourglass - L5a slender
L5b hourglass - L5a long basal
L5b hourglass - L5a wide basal
L5b hourglass - L5b long wide
L5b hourglass - L5b wide
L5b wide - L5 short wide
L5b wide - L5 slender
L5b wide - L5 tufted
L5b wide - L5a slender
L5b wide - L5a long basal
L5b wide - L5a wide basal
L5b wide - L5b long wide
L6 non-tufted - L6 short
L6 non-tufted - L6 tufted
L6 short - L6 tufted
L6 slender - L6 non-tufted
L6 slender - L6 short
L6 slender - L6 tufted
fdr-BH corrected p-value
ns
p<0.05
p<0.01
p<0.001
log2(fold-change)
−2
0
2
L3 (pA)
Claustrum (pA)
r-b distal (pA)
d-b distal L2 (pA)
d-b distal L2 (# sites)
d-b distal L3 (pA)
r-b distal L3 (pA)
d-b distal L3 (# sites)
r-b distal L5 (pA)
d-b distal L6 (pA)
r-b distal L6 (pA)
d-b distal (# sites)
input

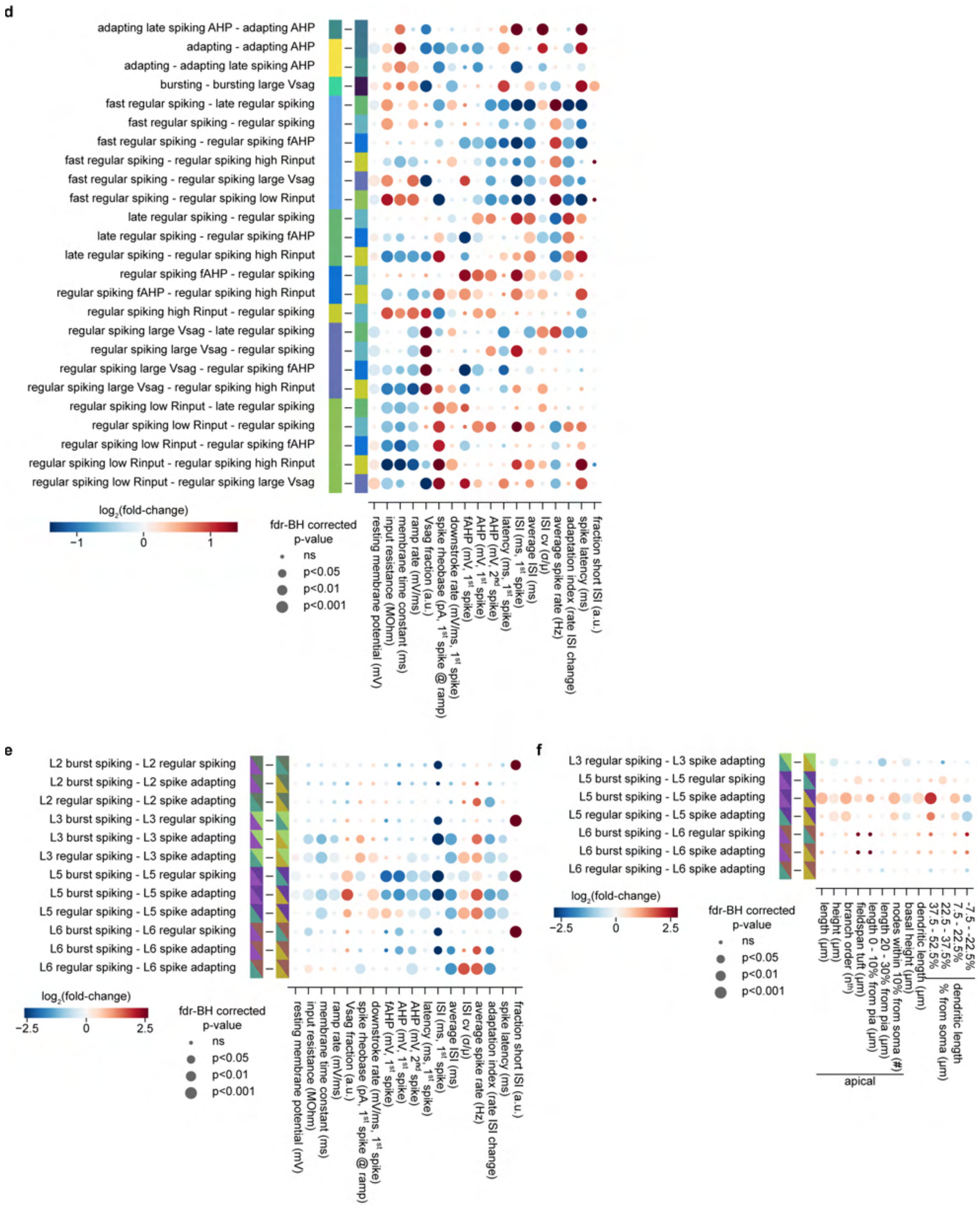
d
adapting late spiking AHP - adapting AHP
adapting - adapting AHP
adapting - adapting late spiking AHP
bursting - bursting large Vsag
fast regular spiking - late regular spiking
fast regular spiking - regular spiking
fast regular spiking - regular spiking fAHP
fast regular spiking - regular spiking high Rinput
fast regular spiking - regular spiking large Vsag
fast regular spiking - regular spiking low Rinput
late regular spiking - regular spiking
late regular spiking - regular spiking fAHP
late regular spiking - regular spiking high Rinput
regular spiking fAHP - regular spiking
regular spiking fAHP - regular spiking high Rinput
regular spiking high Rinput - regular spiking
regular spiking large Vsag - late regular spiking
regular spiking large Vsag - regular spiking
regular spiking large Vsag - regular spiking fAHP
regular spiking large Vsag - regular spiking high Rinput
regular spiking low Rinput - late regular spiking
regular spiking low Rinput - regular spiking
regular spiking low Rinput - regular spiking fAHP
regular spiking low Rinput - regular spiking high Rinput
regular spiking low Rinput - regular spiking large Vsag
log2(fold-change)
-1
0
1
fdr-BH corrected p-value
ns
p<0.05
p<0.01
p<0.001
resting membrane potential (mV)
input resistance (MOhm)
membrane time constant (ms)
ramp rate (mV/ms)
Vsag fraction (a.u.)
spike rheobase (pA, 1st spike @ ramp)
downstroke rate (mV/ms, 1st spike)
fAHP (mV, 1st spike)
AHP (mV, 1st spike)
AHP (mV, 2nd spike)
latency (ms, 1st spike)
ISI (ms, 1st spike)
average ISI (ms)
ISI cv (σ/μ)
average spike rate (Hz)
adaptation index (rate ISI change)
spike latency (ms)
fraction short ISI (a.u.)
e
L2 burst spiking - L2 regular spiking
L2 burst spiking - L2 spike adapting
L2 regular spiking - L2 spike adapting
L3 burst spiking - L3 regular spiking
L3 burst spiking - L3 spike adapting
L3 regular spiking - L3 spike adapting
L5 burst spiking - L5 regular spiking
L5 burst spiking - L5 spike adapting
L5 regular spiking - L5 spike adapting
L6 burst spiking - L6 regular spiking
L6 burst spiking - L6 spike adapting
L6 regular spiking - L6 spike adapting
log2(fold-change)
-2.5
0
2.5
fdr-BH corrected p-value
ns
p<0.05
p<0.01
p<0.001
resting membrane potential (mV)
input resistance (MOhm)
membrane time constant (ms)
ramp rate (mV/ms)
Vsag fraction (a.u.)
spike rheobase (pA, 1st spike @ ramp)
downstroke rate (mV/ms, 1st spike)
fAHP (mV, 1st spike)
AHP (mV, 1st spike)
AHP (mV, 2nd spike)
latency (ms, 1st spike)
ISI (ms, 1st spike)
average ISI (ms)
ISI cv (σ/μ)
average spike rate (Hz)
adaptation index (rate ISI change)
spike latency (ms)
fraction short ISI (a.u.)
f
L3 regular spiking - L3 spike adapting
L5 burst spiking - L5 regular spiking
L5 burst spiking - L5 spike adapting
L5 regular spiking - L5 spike adapting
L6 burst spiking - L6 regular spiking
L6 burst spiking - L6 spike adapting
L6 regular spiking - L6 spike adapting
log2(fold-change)
-2.5
0
2.5
fdr-BH corrected p-value
ns
p<0.05
p<0.01
p<0.001
length (μm)
height (μm)
branch order (nth)
fieldspan tuft (μm)
length 0 - 10% from pia (μm)
length 20 - 30% from pia (μm)
nodes within 10% from soma (#)
basal height (μm)
dendritic length (μm)
37.5 - 52.5%
22.5 - 37.5%
7.5 - 22.5%
-7.5 - -22.5%
% from soma (μm)
dendritic length
apical

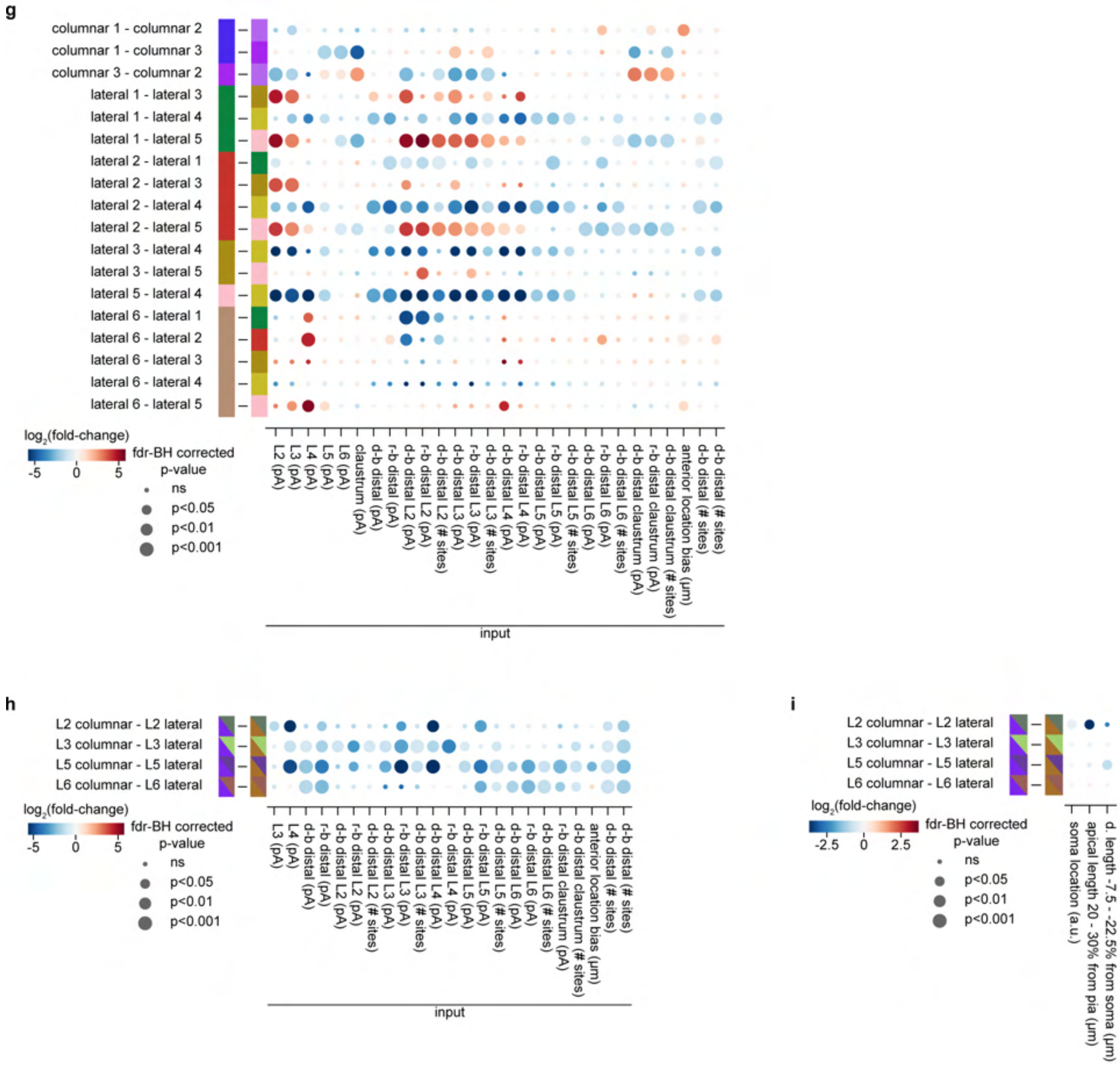
g
columnar 1 - columnar 2
columnar 1 - columnar 3
columnar 3 - columnar 2
lateral 1 - lateral 3
lateral 1 - lateral 4
lateral 1 - lateral 5
lateral 2 - lateral 1
lateral 2 - lateral 3
lateral 2 - lateral 4
lateral 2 - lateral 5
lateral 3 - lateral 4
lateral 3 - lateral 5
lateral 5 - lateral 4
lateral 6 - lateral 1
lateral 6 - lateral 2
lateral 6 - lateral 3
lateral 6 - lateral 4
lateral 6 - lateral 5
log2(fold-change)
-5
0
5
fdr-BH corrected p-value
ns
p<0.05
p<0.01
p<0.001
L2 (pA)
L3 (pA)
L4 (pA)
L5 (pA)
L6 (pA)
claustrum (pA)
d-b distal (pA)
r-b distal (pA)
d-b distal L2 (pA)
r-b distal L2 (pA)
d-b distal L2 (# sites)
d-b distal L3 (pA)
r-b distal L3 (pA)
d-b distal L3 (# sites)
d-b distal L4 (pA)
r-b distal L4 (pA)
d-b distal L5 (pA)
r-b distal L5 (pA)
d-b distal L5 (# sites)
d-b distal L6 (pA)
r-b distal L6 (pA)
d-b distal L6 (# sites)
d-b distal claustrum (pA)
r-b distal claustrum (pA)
d-b distal claustrum (# sites)
anterior location bias (µm)
d-b distal (# sites)
d-b distal (# sites)
input
h
L2 columnar - L2 lateral
L3 columnar - L3 lateral
L5 columnar - L5 lateral
L6 columnar - L6 lateral
log2(fold-change)
-5
0
5
fdr-BH corrected p-value
ns
p<0.05
p<0.01
p<0.001
L3 (pA)
L4 (pA)
d-b distal (pA)
r-b distal (pA)
d-b distal L2 (pA)
r-b distal L2 (pA)
d-b distal L2 (# sites)
d-b distal L3 (pA)
r-b distal L3 (pA)
d-b distal L3 (# sites)
d-b distal L4 (pA)
r-b distal L4 (pA)
d-b distal L5 (pA)
r-b distal L5 (pA)
d-b distal L5 (# sites)
d-b distal L6 (pA)
r-b distal L6 (pA)
d-b distal L6 (# sites)
r-b distal claustrum (pA)
d-b distal claustrum (# sites)
anterior location bias (µm)
d-b distal (# sites)
d-b distal (# sites)
input
i
L2 columnar - L2 lateral
L3 columnar - L3 lateral
L5 columnar - L5 lateral
L6 columnar - L6 lateral
log2(fold-change)
-2.5
0
2.5
fdr-BH corrected p-value
ns
p<0.05
p<0.01
p<0.001
soma location (a.u.)
apical length 20 - 30% from pia (µm)
d. length -7.5 - -22.5% from soma (µm)

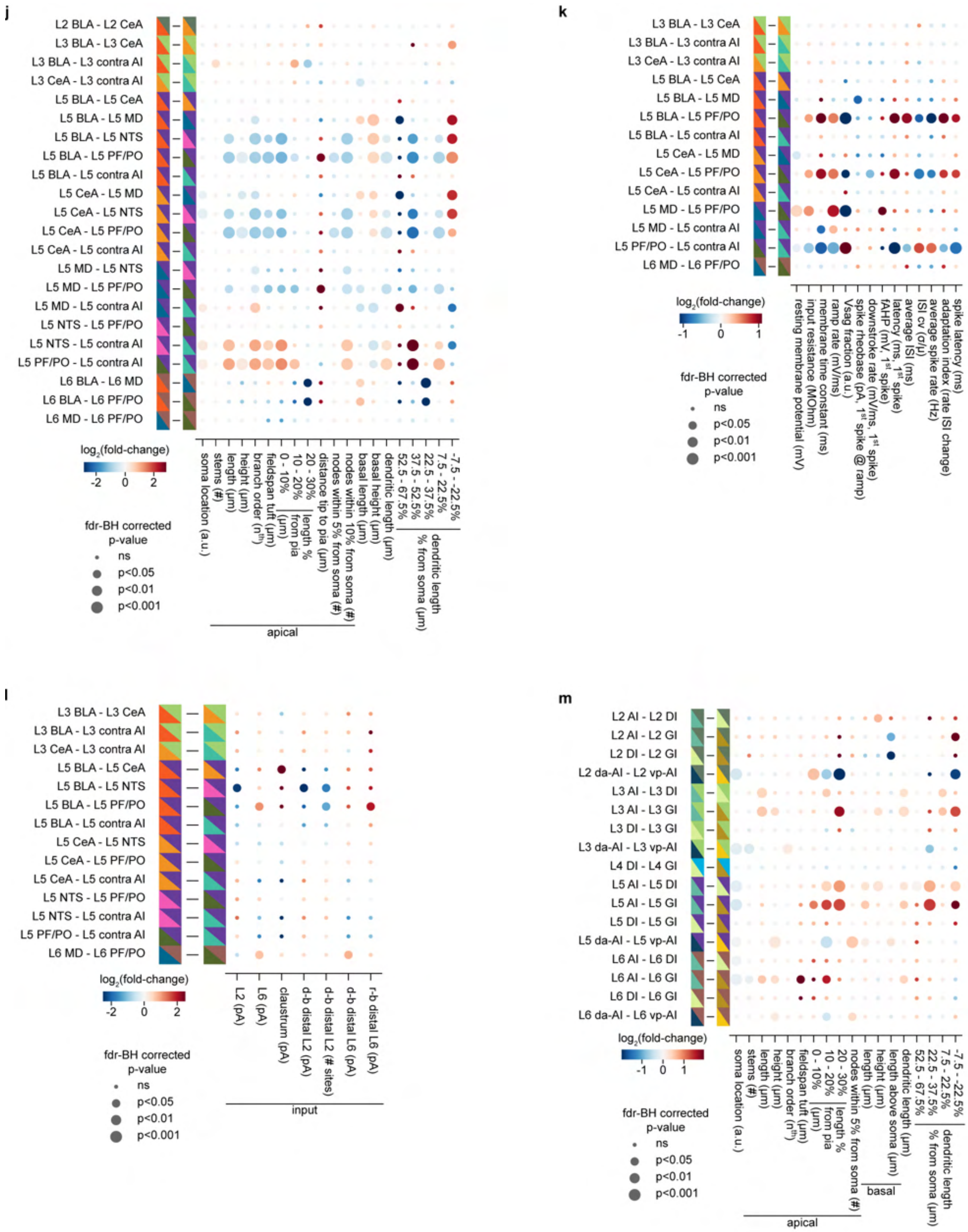
j
L2 BLA - L2 CeA
L3 BLA - L3 CeA
L3 BLA - L3 contra AI
L3 CeA - L3 contra AI
L5 BLA - L5 CeA
L5 BLA - L5 MD
L5 BLA - L5 NTS
L5 BLA - L5 PF/PO
L5 BLA - L5 contra AI
L5 CeA - L5 MD
L5 CeA - L5 NTS
L5 CeA - L5 PF/PO
L5 CeA - L5 contra AI
L5 MD - L5 NTS
L5 MD - L5 PF/PO
L5 MD - L5 contra AI
L5 NTS - L5 PF/PO
L5 NTS - L5 contra AI
L5 PF/PO - L5 contra AI
L6 BLA - L6 MD
L6 BLA - L6 PF/PO
L6 MD - L6 PF/PO
log2(fold-change)
-2 0 2
fdr-BH corrected p-value
ns
p<0.05
p<0.01
p<0.001
soma location (a.u.)
stems (#)
length (µm)
height (µm)
branch order (nth)
fieldspan tuft (µm)
0 - 10%
10 - 20%
20 - 30%
length % from pia (µm)
distance tip to pia (µm)
nodes within 5% from soma (#)
nodes within 10% from soma (#)
basal length (µm)
basal height (µm)
dendritic length (µm)
52.5 - 67.5%
37.5 - 52.5%
22.5 - 37.5%
7.5 - 22.5%
-7.5 - -22.5%
dendritic length % from soma (µm)
apical
k
L3 BLA - L3 CeA
L3 BLA - L3 contra AI
L3 CeA - L3 contra AI
L5 BLA - L5 CeA
L5 BLA - L5 MD
L5 BLA - L5 PF/PO
L5 BLA - L5 contra AI
L5 CeA - L5 MD
L5 CeA - L5 PF/PO
L5 CeA - L5 contra AI
L5 MD - L5 PF/PO
L5 MD - L5 contra AI
L5 PF/PO - L5 contra AI
L6 MD - L6 PF/PO
log2(fold-change)
-1 0 1
fdr-BH corrected p-value
ns
p<0.05
p<0.01
p<0.001
resting membrane potential (mV)
input resistance (MOhm)
membrane time constant (ms)
ramp rate (mV/ms)
Vsag fraction (a.u.)
spike rheobase (pA, 1st spike @ ramp)
downstroke rate (mV/ms, 1st spike)
fAHP (mV, 1st spike)
latency (ms, 1st spike)
average ISI (ms)
ISI cv (σ/µ)
average spike rate (Hz)
adaptation index (rate ISI change)
spike latency (ms)
l
L3 BLA - L3 CeA
L3 BLA - L3 contra AI
L3 CeA - L3 contra AI
L5 BLA - L5 CeA
L5 BLA - L5 NTS
L5 BLA - L5 PF/PO
L5 BLA - L5 contra AI
L5 CeA - L5 NTS
L5 CeA - L5 PF/PO
L5 CeA - L5 contra AI
L5 NTS - L5 PF/PO
L5 NTS - L5 contra AI
L5 PF/PO - L5 contra AI
L6 MD - L6 PF/PO
log2(fold-change)
-2 0 2
fdr-BH corrected p-value
ns
p<0.05
p<0.01
p<0.001
L2 (pA)
L6 (pA)
claustrum (pA)
d-b distal L2 (pA)
d-b distal L2 (# sites)
d-b distal L6 (pA)
r-b distal L6 (pA)
input
m
L2 AI - L2 DI
L2 AI - L2 GI
L2 DI - L2 GI
L2 da-AI - L2 vp-AI
L3 AI - L3 DI
L3 AI - L3 GI
L3 DI - L3 GI
L3 da-AI - L3 vp-AI
L4 DI - L4 GI
L5 AI - L5 DI
L5 AI - L5 GI
L5 DI - L5 GI
L5 da-AI - L5 vp-AI
L6 AI - L6 DI
L6 AI - L6 GI
L6 DI - L6 GI
L6 da-AI - L6 vp-AI
log2(fold-change)
-1 0 1
fdr-BH corrected p-value
ns
p<0.05
p<0.01
p<0.001
soma location (a.u.)
stems (#)
length (µm)
height (µm)
branch order (nth)
fieldspan tuft (µm)
0 - 10%
10 - 20%
20 - 30%
length % from pia (µm)
nodes within 5% from soma (#)
length (µm)
height (µm)
length above soma (µm)
dendritic length (µm)
52.5 - 67.5%
22.5 - 37.5%
7.5 - 22.5%
-7.5 - -22.5%
dendritic length % from soma (µm)
apical
basal

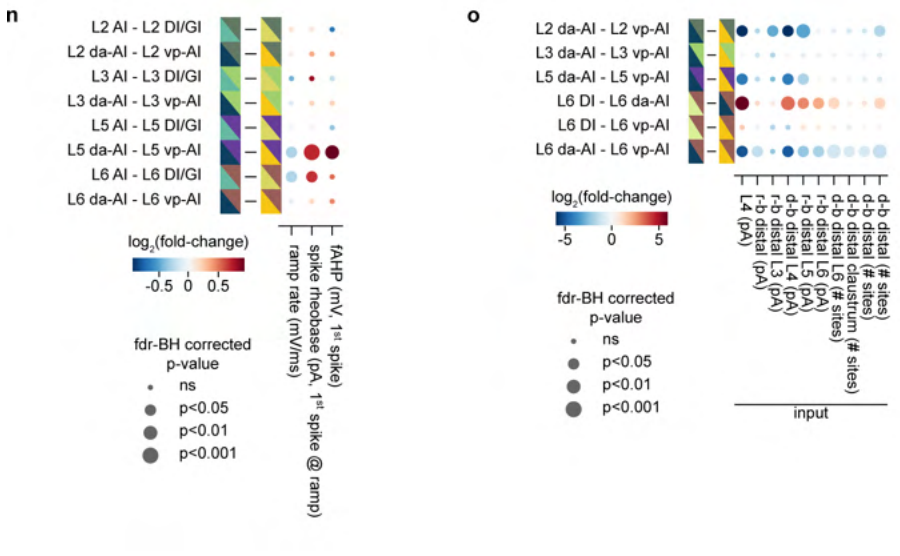

**Supplementary Fig. 3: Informative features for pair-wise comparisons between cell types across all modalities. a–o**, Pair-wise cross-modal feature comparisons between cell types (n ≥ 3 cells per type) of morphological (**a**–**c**), electrical (**d**–**f**), input (**g**–**l**), and topographical (**m**–**o**) modalities. In each row, the left and right cell types are color-coded as defined earlier (Figs. 1c, 2e, 3d, 4a, and 5d). For cell types based on the intersection of two modalities (**h**–**o**), e.g., layer allocation and input class (**h**), colors corresponding to each modality are shown in triangles. All features used for the iCCA are included (Extended Data Figs. 5e, 8b, and 10b). Bubble plots show fold difference calculated as $\log_2$ of mean feature value of the left cell type divided by that of the right cell type. Red hue: the mean value of the left cell type > mean value of the right cell type; blue hue: the reverse. All comparisons were statistically tested using a Mann-Whitney U test with false-discovery rate Benjamini-Hochberg corrections (fdr-BH corr.), indicated by four bubble sizes. Only features with a significant difference between any cell type pair are shown. For distal input features, the distal input border was defined by boundaries based on either 95% dendritic density (density-based, d-b) or maximum dendritic reach (reach-based, r-b).

| Modality | Category | Feature | Unit | Package | Description |
|---|---|---|---|---|---|
| Morphology | Soma | Soma location | relative | BCS | Soma location relative to lct. |
| | Soma | Soma diameter | µm | BCS | Soma diameter. |
| | Apical / basal / apical + basal | Length | µm | L-measure | Total length of dendrite. |
| | Apical / basal | Nodes | nodes | L-measure | Total number of branch points. |
| | Apical / basal | Nodes within r pct from soma | nodes | BCS | Number of branch points within a radius (r) percentage of lct around the soma. |
| | Apical / basal | Normalized nodes within r pct from soma | relative | BCS | Number of branch points within a radius (r) percentage of lct around the soma relative to total branch points. |
| | Apical / basal / apical + basal | Branches | branches | L-measure | Total number of segments, a segment is a stretch of dendrite between two branch points or between branch point and end point. |
| M or | Apical / basal | Tips | tips | L-measure | Total number of end points. |

| | | | | | |
|---|---|---|---|---|---|
| | Apical / basal / apical + basal | Width | µm | L-measure | Dendritic extent parallel to the pia (X-axis). Distance between the outer points of the 95 percentiles of all traced points along the X-axis. |
| | Apical / basal / apical + basal | Height | µm | L-measure | Dendritic extent perpendicular to the pia (Y-axis). Distance between the outer points of the 95 percentiles of all traced points along the Y-axis. |
| | Apical | Normalized height | relative | BCS | Dendritic height relative to the lct. |
| | Apical / basal | Euclidean distance | µm | L-measure | Summed Euclidean distance between soma to each traced point. |
| | Apical / basal | Path distance | µm | L-measure | Summed path length, distance between soma to each traced point. |
| | Apical / basal | Partition asymmetry | relative | L-measure | Averaged partition asymmetry, per bifurcation the asymmetry is calculated as the difference in total tips located at each of the two branches / (total number of tips at both branches – 2). |
| Morphology | Apical / basal | Bifurcation angle local | degrees | L-measure | Averaged angle between two segments after bifurcation. |

| | | | | | |
|---|---|---|---|---|---|
| | Apical / basal | Bifurcation angle remote | degrees | L-measure | Averaged angle between two bifurcation points or between bifurcation point and end point or between two end points. |
| | Apical / basal | Contraction | ratio | L-measure | Averaged ratio between Euclidean and path distance of a branch. |
| | Apical / basal | Branch order | order | L-measure | Averaged order of the branch with respect to the soma for each segment. |
| | Apical | Field span near pia | µm | BCS | Dendritic extent parallel to the pia (X-axis) located within the first 5% lct from pia. |
| | Apical | Normalized length Y1 - Y2 pct from pia | relative | BCS | Local dendritic length located in the specified cortical depth range, Y1 to Y2 percentage of lct away from the pia. Local dendritic length is normalized to total apical length. |
| Morphology | Apical | Normalized reach to pia | relative | BCS | Distance between pia and farthest dendritic extent along Y-axis towards pia relative to the lct. (1 - farthest dendritic extent Y) / lct, 0: dendrite reaches pia. |

| | | | | | |
|---|---|---|---|---|---|
| | Apical | Reach potential to pia | relative | BCS | Normalized reach to pia relative to soma location. Soma location - (normalized reach to pia / soma location). |
| | Apical | Fraction below soma | relative | BCS | Dendritic length located below the soma relative to the total apical length. |
| | Basal | Fraction above soma | relative | BCS | Dendritic length located above the soma relative to the total basal length. |
| | Basal | Stems | stems | L-measure | Total number of basal stems attached to the soma. |
| | Apical + basal | Overlap binary per 18 µm lamina | relative | BCS | Number of layer bins, 18 µm along Y-axis, in which both apical and basal dendrites are present, relative to total number of layer bins at the lct. [a] |
| Morphology | Apical + basal | Overlap binary per 18 µm voxel | relative | BCS | Number of voxels (18 × 18 × 18 µm) in which both apical and basal dendrites are present, relative to total number of voxels with a dendrite present. [a] |

| | | | | | |
|---|---|---|---|---|---|
| | Apical + basal | Overlap density per 18 µm lamina | ratio | BCS | Ratio between dendritic density located in layer bins, 18 µm along Y-axis, in which both apical and basal dendrites are present, and the total dendritic length. [a, b]. |
| | Apical + basal | Overlap density per 18 µm voxel | ratio | BCS | Ratio between dendritic density located within voxels (18 × 18 ×18 µm) in which both apical and basal dendrites are present, and the total dendritic length. [a,b] |
| | Apical + basal | Ratio length | ratio | BCS | Ratio between total apical and total basal dendritic length. |
| | Apical + basal | Ratio width | ratio | BCS | Ratio between total apical and total basal dendritic width. |
| | Apical + basal | Ratio height | ratio | BCS | Ratio between total apical and total basal dendritic height. |
| Morphology | Basal | Ratio height / width | ratio | BCS | Ratio between total basal dendritic height and total basal dendritic width. |
| | Apical + basal | Ratio height / width | ratio | BCS | Ratio between total dendritic height and total dendritic width. |
| | Apical + basal | Normalized length Y1 - Y2 pct from soma | relative | BCS | Local dendritic length located in the specified cortical depth range, Y1 to Y2 percentage of lct away |

| | | | | | |
|---|---|---|---|---|---|
| | | | | | from the soma. Local dendritic length is normalized to lct. |
| Electrophysiology | Passive properties | Resting membrane potential | mV | BCS | Resting membrane potential during the first 100 ms of the trace or until start stimulation. Median of the first 20 recorded traces (~ first 100 seconds of whole-cell patch-clamp recording). |
| | Passive properties | Input resistance | Mohm | BCS | Slope of the linear fit to maximum hyperpolarized membrane potential during ≥ 200 ms long current steps of 25 pA ranging between -150 to 0 pA. If not available the input resistance was estimated based on the test pulses (-2.5 to -5 mV) recorded in voltage-clamp traces: voltage step / steady-state current response. |
| Electrophysiology | Passive properties | Membrane time constant | ms | BCS | Single-exponential fit to the rising voltage trace. Median of all traces with ≥ 200 ms long current injections of -150 to -25 pA. |

| | | | | | |
|---|---|---|---|---|---|
| | Passive properties | Ramp rate | mV/s | BCS | Membrane charging velocity during linear current injection (165pA/s). Slope of the linear fit between start ramp until spike threshold. Median of all ramping current injections. |
| | Rectifying properties | voltage sag | relative | BCS | Voltage difference between maximum hyperpolarized and steady-state membrane potentials relative to steady-state membrane potential. Median of all traces with ≥ 200 ms long current injections of -300 pA. |
| | Rectifying properties | fAHP | mV | BCS | Membrane potential difference between spike threshold and maximum hyperpolarization within 5ms from peak of first AP. Median of all traces with ≥ 200 ms long current injections at rheobase. |
| Electrophysiology | Rectifying properties | AHP | mV | BCS | Membrane potential difference between spike threshold of first AP and maximum hyperpolarization between first and second APs. Median of all traces with ≥ 200 ms long current injections at rheobase. |

| | | | | | |
|---|---|---|---|---|---|
| | Rectifying properties | AHP second AP during spike train | mV | BCS | Membrane potential difference between spike threshold of second AP and maximum hyperpolarization between second and third APs. Median of all traces with ≥ 1 ms long current injections at rheobase + 125 pA. |
| | AP waveform | AP threshold | mV | BCS | Membrane potential at AP threshold. Threshold is defined as the first moment membrane potential exceeds 5% of the average maximum upstroke rate. |
| | AP waveform | AP height | mV | BCS | AP height between threshold and peak of the first AP. |
| | AP waveform | AP width | ms | BCS | Full-width at half-max of AP peak of the first AP. |
| | AP waveform | AP upstroke rate | mV/ms | BCS | Maximum velocity of upstroke of the first AP. |
| Electrophysiology | AP waveform | AP downstroke rate | mV/ms | BCS | Maximum velocity of downstroke of the first AP. |

| | | | | | |
|---|---|---|---|---|---|
| | AP waveform | AP upstroke to downstroke ratio | ratio | BCS | Ratio between up and down stroke rates for the first AP. |
| | Spiking behavior | Rheobase | pA | BCS | Current required to elicit first AP with a square-pulse depolarizing current injection. |
| | Spiking behavior | Rheobase during ramp | pA | BCS | Current required to elicit first AP during a ramping depolarizing current injection. |
| | Spiking behavior | Charge at rheobase | pA/s | BCS | Charge required to elicit first AP with a square-pulse current injection. [b] |
| | Spiking behavior | AP latency at rheobase | ms | BCS | Latency between stimulation onset and threshold of the first AP during depolarizing current injections. |
| | Spiking behavior | First AP ISI | ms | BCS | Inter-spike interval (ISI) between first and second APs. |
| | Spiking behavior | AP ISI | ms | BCS | ISI between all APs. |
| Electrophysiology | Spiking behavior | ISI CV | a.u. | BCS | Coefficient of variation of ISI. [b] |
| | Spiking behavior | AP rate | spike/s | BCS | Number of AP / current injection time. |
| | Spiking behavior | Spike adaptation | a.u. | BCS | Rate at which spiking speeds up or slows down. 1 / (number of APs - |

| | | | | | |
|---|---|---|---|---|---|
| | | | | | 1) × sum(ISI[$AP_n$ + 1] - ISI[$AP_n$] / ISI[$AP_{n+1}$] + ISI[$AP_n$]). 0: regular spiking; 0 >: spiking slows down; 0 <: spiking speeds up. |
| | Spiking behavior | AP firing to current injection rate | Hz/pA | BCS | Slope of linear fit to the spike rate over current injection amplitude. [b] |
| | Spiking behavior | AP latency at rheobase + 125 pA | ms | BCS | Latency between stimulation onset and threshold of the first AP. |
| | Spiking behavior | AP height desensitization | ratio | BCS | Ratio between first and last AP height. |
| | Spiking behavior | AP width desensitization | ratio | BCS | Ratio between first and last AP width. |
| | Spiking behavior | Fraction short ISI | relative | BCS | Fraction of short ISIs (< 10 ms) over all ISIs. |
| | Spiking behavior | Longest AP burst | ms | BCS | Time that a neuron spends in bursting state (inter-spike interval < 10 ms). [b] |
| Input | Layer-stratified input | Input strength by layer | relative | BCS | Summed input strengths of all locations allocated to layer x relative to total input strength. |
| | Layer-stratified input | Distal input strength by layer | relative | BCS | Summed input strength of all locations allocated to layer x beyond a lateral distance from the soma relative to total input |

| | | | | | |
|---|---|---|---|---|---|
| | | | | | strength of layer x. Lateral distance threshold was based on dataset's 95% dendritic reach (675µm), or 95% dendritic density (350 µm). |
| | Layer-stratified input | Distal input location by layer | relative | BCS | Summed input locations allocated to layer x beyond a lateral distance from the soma relative to total input locations in layer x. Lateral distance threshold was based on dataset's 95% dendritic reach (675µm), or 95% dendritic density (350 µm). |
| | Input relative to soma | Input strength above soma | relative | BCS | Input strength above soma relative to total input strength. |
| | Input relative to soma | Farthest input location anterior and posterior | µm | BCS | Farthest input location relative to soma on the anterior and posterior side. |
| | Input relative to soma | Lateral input location span | µm | BCS | Distance farthest anterior to posterior input locations. |
| Input | Input relative to soma | Anterior input location bias | µm | BCS | -1 × farthest input location anterior + farthest input location posterior |
| | Input relative to soma | Anterior input strength | relative | BCS | Anterior input strength relative to total input strength |
| | Input relative to soma | Distance-weighted | relative | BCS | Distance-weighted anterior input strength relative to total distance-weighted input strength |

| | | |
|---|---|---|
| | | anterior input strength |

**Supplementary Table 1: Overview of morphology, electrophysiology, and input properties/features.** Several features were extracted for multiple categories. For example, for the morphology feature ‘length’, the length of either the basal or apical dendrite alone as well as the summed basal + apical dendrite length was extracted. Features were either extracted using L-measure[97] or BrainCellSuite (BCS). For all spiking behavior category features the median value over all traces with ≥ 1 s long current injections at rheobase + 125 pA were used. For all action potential (AP) waveform category features the median value of all first APs during ≥ 200 ms long current injections at rheobase were used. For all input features individual input maps were first interpolated along the Y-axis between pia and corpus callosum and redistributed among 22 cortical bins. All stimulation locations in the interpolated map were matched to the Nissl atlas to allocate each location to a layer. [a] Dimension was based on averaged soma diameter ~18 µm (18.07 µm). [b] Skewed features were $log_2$ transformed; for electrophysiology features a small offset (0.1 unit) was added. ISI: Inter-spike interval, lct: local cortical thickness, pct: percentage.

| **Short name** | **Strain** | **RRID** | **Catalog number** | **Supplier** |
|---|---|---|---|---|
| wild-type (WT) | C57BL/6NCrl (WT) | IMSR_CR:027 | 027 | Charles river |
| PV-Cre | B6.129P2-*Pvalbtm1(cre)Arbr*/J | IMSR_JAX:008069 | 8069 | The Jackson Laboratory |
| SST-Cre | STOCK Ssttm2.1(cre)Zjh/J | IMSR_JAX:013044 | 13044 | The Jackson Laboratory |
| Gad2-Cre | Gad2tm2(cre)Zjk/J | IMSR_JAX:010802 | 10802 | The Jackson Laboratory |
| Drd1-EGFP | STOCK Tg(Drd1-EGFP)X60Gsat/Mmmh | MMRRCC_000297-MU | 000297-MU | Gensat |
| Dat-Cre | B6.SJL-Slc6a3tm1.1(cre)Bkmn/J | IMSR_JAX:006660 | 6660 | The Jackson Laboratory |
| Ai32 | B6;129S-Gt(ROSA)26Sortm32(CAG-COP4*H134R/EYFP)Hze/J | IMSR_JAX:012569 | 12569 | The Jackson Laboratory |
| Ai9 | B6.Cg-Gt(ROSA)26Sortm9(CAG-tdTomato)Hze/J | IMSR_JAX:007909 | 7909 | The Jackson Laboratory |

**Supplementary Table 2: Mouse strains included in this database.**

| Genotype (short names) | Female | Male |
|---|---|---|
| *Dat-Cre*$^{+/-}$ | 0 / 0 | 4 / 1 |
| *Dat-Cre*$^{+/-}$;*Ai32*$^{+/-}$ | 9 / 2 | 0 / 0 |
| *Drd1-EGFP*$^{+/-}$ | 1 / 1 | 1 / 1 |
| *Gad2-Cre*$^{+/-}$ | 3 / 3 | 2 / 2 |
| *Gad2-Cre*$^{+/-}$;*Ai9*$^{+/-}$ | 8 / 2 | 1 / 1 |
| *PV-Cre*$^{+/-}$ | 4 / 3 | 1 / 1 |
| *PV-Cre*$^{+/-}$;*Ai32*$^{+/-}$ | 76 / 20 | 36 / 12 |
| *PV-Cre*$^{+/-}$;*Ai9*$^{+/-}$ | 153 / 46 | 340 / 65 |
| *SST*$^{+/-}$;*Ai32*$^{+/-}$ | 3 / 1 | 24 / 5 |
| WT | 239 / 53 | 170 / 59 |
| genotyped-WT* | 5 / 2 | 13 / 3 |

**Supplementary Table 3: Number of observations in this dataset tabulated by mouse strain and sex**. * Genotyped-WT are offspring of the strains mentioned in this tabulation with genotyped homozygous wildtype (C57BL) alleles at the transgene location.
